# Genetic Disruption of WASHC4 Drives Endo-lysosomal Dysfunction and Cognitive-Movement Impairments in Mice and Humans

**DOI:** 10.1101/2020.08.06.239517

**Authors:** Jamie L. Courtland, Tyler W. A. Bradshaw, Greg Waitt, Erik J. Soderblom, Tricia Ho, Anna Rajab, Ricardo Vancini, Hwan Kim, Scott H. Soderling

**Affiliations:** Department of Neurobiology, Duke University School of Medicine, Durham, NC 27710, USA; Department of Cell Biology, Duke University School of Medicine, Durham, NC 27710, USA; Proteomics and Metabolomics Shared Resource, Duke University School of Medicine, Durham, NC 27710, USA; Burjeel Hospital, VPS Healthcare, Muscat, Oman; Department of Pathology, Duke University School of Medicine, Durham, NC 27710, USA; Department of Anatomy and Neurobiology, University of Tennessee Heath Science Center, Memphis, TN 38163, USA

## Abstract

Mutation of the WASH complex subunit, SWIP, is implicated in human intellectual disability, but the cellular etiology of this association is unknown. We identify the neuronal WASH complex proteome, revealing a network of endosomal proteins. To uncover how dysfunction of endosomal SWIP leads to disease, we generate a mouse model of the human *WASHC4^c.3056C>G^* mutation. Quantitative spatial proteomics analysis of SWIP^P1019R^ mouse brain reveals that this mutation destabilizes the WASH complex and uncovers significant perturbations in both endosomal and lysosomal pathways. Cellular and histological analyses confirm that SWIP^P1019R^ results in endo-lysosomal disruption and uncover indicators of neurodegeneration. We find that SWIP^P1019R^ not only impacts cognition, but also causes significant progressive motor deficits in mice. Remarkably, a retrospective analysis of SWIP^P1019R^ patients confirms motor deficits in humans. Combined, these findings support the model that WASH complex destabilization, resulting from SWIP^P1019R^, drives cognitive and motor impairments via endo-lysosomal dysfunction in the brain.

## INTRODUCTION

Neurons maintain precise control of their subcellular proteome using a sophisticated network of vesicular trafficking pathways that shuttle cargo throughout their elaborate processes. Endosomes function as a central hub in this vesicular relay system by coordinating protein sorting between multiple cellular compartments, including surface receptor endocytosis and recycling, as well as degradative shunting to the lysosome. How endosomal trafficking is modulated in neurons remains a vital area of research due to the unique degree of spatial segregation between organelles in neurons, and its strong implication in neurodevelopmental and neurodegenerative diseases.

In non-neuronal cells, an evolutionarily conserved complex, the Wiskott-Aldrich Syndrome protein and SCAR Homology (WASH) complex, coordinates endosomal trafficking (Derivery and Gautreau, 2010; Linardopoulou et al., 2007). WASH is composed of five core protein components: WASHC1 (aka WASH1), WASHC2 (aka FAM21), WASHC3 (aka CCDC53), WASHC4 (aka SWIP), and WASHC5 (aka Strumpellin) (encoded by genes *Washc1-Washc5*, respectively), which are broadly expressed in multiple organ systems (Alekhina et al., 2017; Kustermann et al., 2018; McNally et al., 2017; Simonetti and Cullen, 2019; Thul et al., 2017). The WASH complex plays a central role in non-neuronal endosomal trafficking by activating Arp2/3-dependent actin branching at the outer surface of endosomes to influence cargo sorting and vesicular scission (Gomez and Billadeau, 2009; Lee et al., 2016; Phillips-Krawczak et al., 2015; Piotrowski et al., 2013; Simonetti and Cullen, 2019). WASH also interacts with at least three main cargo adaptor complexes — the Retromer, Retriever, and COMMD/CCDC22/CCDC93 (CCC) complexes — all of which associate with distinct sorting nexins to select specific cargo and enable their trafficking to other cellular locations (Binda et al., 2019; Farfán et al., 2013; McNally et al., 2017; Phillips-Krawczak et al., 2015; Seaman and Freeman, 2014; Singla et al., 2019). Loss of the WASH complex in non-neuronal cells has detrimental effects on endosomal structure and function, as its loss results in aberrant endosomal tubule elongation and cargo mislocalization (Bartuzi et al., 2016; Derivery et al., 2009; Gomez et al., 2012; Gomez and Billadeau, 2009; Phillips-Krawczak et al., 2015; Piotrowski et al., 2013). However, whether the WASH complex performs an endosomal trafficking role in neurons remains an open question, as no studies have addressed neuronal WASH function to date.

Consistent with the association between the endosomal trafficking system and pathology, dominant missense mutations in *WASHC5* (protein: Strumpellin) are associated with hereditary spastic paraplegia (SPG8) (De Bot et al., 2013; Valdmanis et al., 2007), and autosomal recessive point mutations in *WASHC4* (protein: SWIP) and *WASHC5* are associated with syndromic and non-syndromic intellectual disabilities (Assoum et al., 2020; Elliott et al., 2013; Ropers et al., 2011). In particular, an autosomal recessive mutation in *WASHC4* (c.3056C>G; p.Pro1019Arg) was identified in a cohort of children with non-syndromic intellectual disability (Ropers et al., 2011). Cell lines derived from these patients exhibited decreased abundance of WASH proteins, leading the authors to hypothesize that the observed cognitive deficits in SWIP^P1019R^ patients resulted from disruption of neuronal WASH signaling (Ropers et al., 2011). However, whether this mutation leads to perturbations in neuronal endosomal integrity, or how this might result in cellular changes associated with disease, are unknown.

Here we report the analysis of neuronal WASH and its molecular role in disease pathogenesis. We use *in vivo* proximity proteomics (iBioID) to uncover the neuronal WASH proteome and demonstrate that it is highly enriched for components of endosomal trafficking. We then generate a mouse model of the human *WASHC4^c.3056c>g^* mutation (SWIP^P1019R^) (Ropers et al., 2011) to discover how this mutation may alter neuronal trafficking pathways and test whether it leads to phenotypes congruent with human patients. Using an adapted spatial proteomics approach (Geladaki et al., 2019), coupled with a systems-level analysis of protein covariation networks, we find strong evidence for substantial disruption of neuronal endosomal and lysosomal pathways *in vivo*. Cellular analyses confirm a significant impact on neuronal endo-lysosomal trafficking *in vitro* and *in vivo*, with evidence of lipofuscin accumulation and progressive apoptosis activation, molecular phenotypes that are indicative of neurodegenerative pathology. Behavioral analyses of SWIP^P1019R^ mice at adolescence and adulthood confirm a role of WASH in cognitive processes, and reveal profound, progressive motor dysfunction. Importantly, retrospective examination of SWIP^P1019R^ patient data confirms motor dysfunction coincident with cognitive impairments in humans. Our results establish that impaired WASH complex function leads to altered neuronal endo-lysosomal function, which manifests behaviorally as cognitive and movement impairments.

## RESULTS

### Identification of the WASH complex proteome *in vivo* confirms a neuronal role in endosomal trafficking

While multiple mutations within the WASH complex have been identified in humans (Assoum et al., 2020; Elliott et al., 2013; Ropers et al., 2011; Valdmanis et al., 2007), how these mutations lead to neurological dysfunction remains unknown (Figure 1A). Given that previous work in non-neuronal cultured cells and non-mammalian organisms have established that the WASH complex functions in endosomal trafficking, we first aimed to determine whether this role was conserved in the mouse nervous system (Alekhina et al., 2017; Billadeau et al., 2010; Derivery et al., 2009; Gomez et al., 2012; Gomez and Billadeau, 2009). To discover the likely molecular functions of the neuronal WASH complex, we utilized an *in vivo* BioID (iBioID) paradigm developed in our laboratory to identify the WASH complex proteome from brain tissue (Uezu et al., 2016). BioID probes were generated by fusing a component of the WASH complex, WASH1 (gene: *Washc1*), with the promiscuous biotin ligase, BioID2 (WASH1-BioID2, Figure 1B), or by expressing BioID2 alone (negative control, solubleBioID2) under the neuron-specific, human Synapsin-1 promoter (Kim et al., 2016). We injected adenoviruses (AAV) expressing these constructs into the cortex of wild-type postnatal day zero (P0) mice (Figure 1B). Two weeks post-injection, we administered daily subcutaneous biotin for seven days to biotinylate *in vivo* substrates. The viruses displayed efficient expression and activity in brain tissue, as evidenced by colocalization of the WASH1-BioID2 viral epitope (HA) and biotinylated proteins (Streptavidin) (Figures 1C-F). For label-free quantitative high-mass accuracy LC-MS/MS analyses, whole brain samples were collected at P22, snap-frozen, and processed as previously described (Uezu et al., 2016). A total of 2,311 proteins were identified across all three experimental replicates, which were further analyzed for those with significant enrichment in WASH1-BioID2 samples over solubleBioID2 negative controls (Table S1).

**Figure 1.**
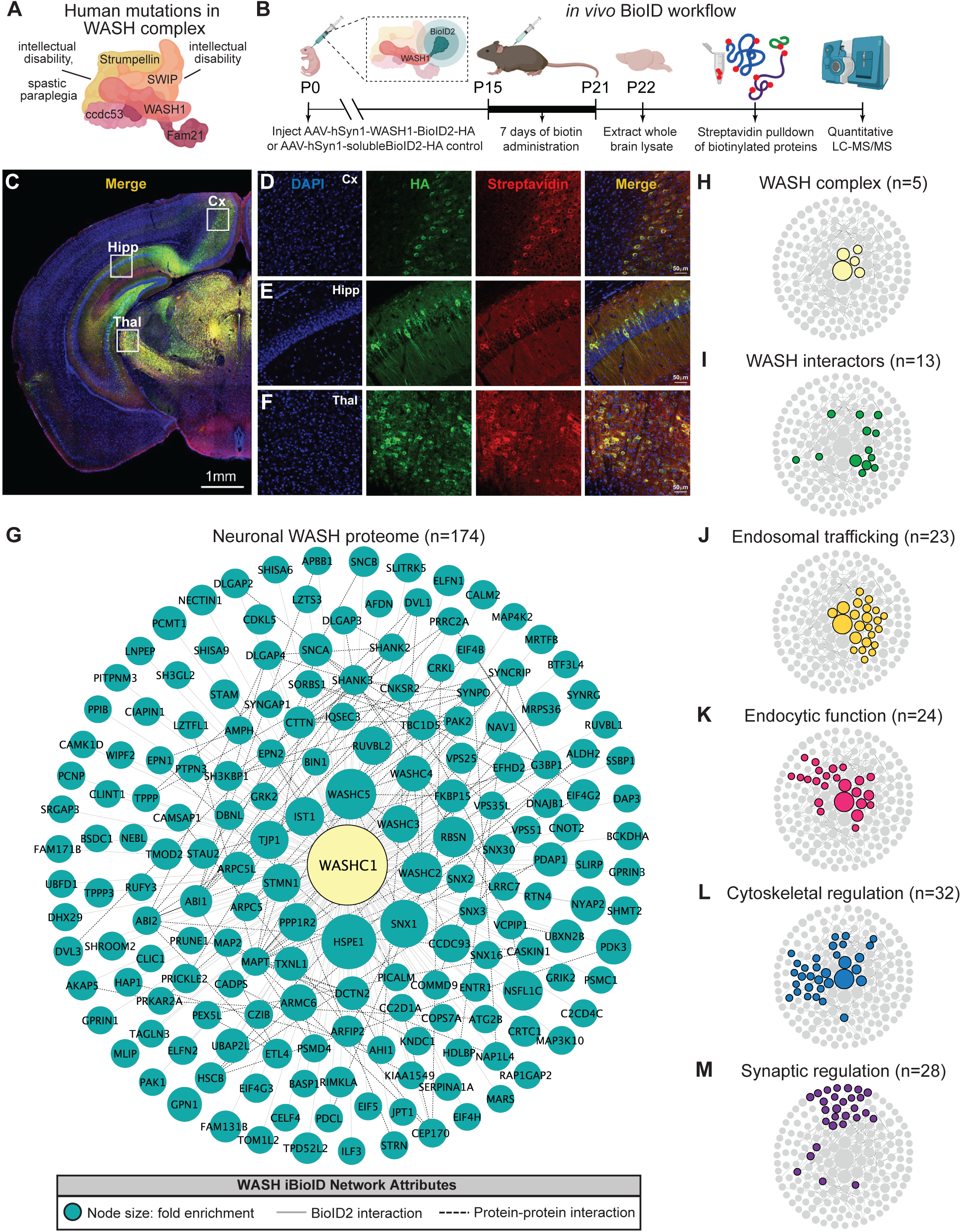
Identification of the WASH complex proteome *in vivo* confirms a neuronal role in endosomal trafficking. (A) The WASH complex is composed of five subunits, *Washc1* (WASH1), *Washc2* (FAM21), *Washc3* (CCDC53), *Washc4* (SWIP), and *Washc5* (Strumpellin). Human mutations in these components are associated with spastic paraplegia (De Bot et al., 2013; Jahic et al., 2015; Valdmanis et al., 2007), Ritscher-Schinzel Syndrome (Elliott et al., 2013), and intellectual disability (Assoum et al., 2020; Ropers et al., 2011). (B) A BioID2 probe was attached to the c-terminus of WASH1 and expressed under the human synapsin-1 (hSyn1) promoter in an AAV construct for in vivo BioID (iBioID). iBioID probes (WASH1-BioID2-HA, or negative control solubleBioID2-HA) were injected into wild-type mouse brain at P0 and allowed to express for two weeks. Subcutaneous biotin injections (24 mg/kg) were administered over seven days for biotinylation, and then brains were harvested for isolation and purification of biotinylated proteins. LC-MS/MS identified proteins significantly enriched in all three replicates of WASH1-BioID2 samples over soluble-BioID2 controls. (C) Representative image of WASH1-BioID2-HA expression in a mouse coronal brain section (Cx = cortex, Hipp = hippocampus, Thal = thalmus). Scale bar, 1 mm. (D) Representative image of WASH1-BioID2-HA expression in mouse cortex (inset from C). Individual panels show nuclei (DAPI, blue), AAV construct HA epitope (green), and biotinylated proteins (Streptavidin, red). Merged image shows colocalization of HA and Streptavidin (yellow). Scale bar, 50 µm. (E) Representative image of WASH1-BioID2-HA expression in mouse hippocampus (inset from C). Scale bar, 50 µm. (F) Representative image of WASH1-BioID2-HA expression in mouse thalamus (inset from C). Scale bar, 50 µm. (G) iBioID identified known and unknown proteins interactors of the WASH complex in murine neurons. Nodes size represents protein abundance fold-enrichment over negative control (range: 3 to 181.7), solid grey edges delineate iBioID interactions between the WASHC1 probe (seen in yellow at the center) and identified proteins, dashed edges indicate known protein-protein interactions from HitPredict database (López et al., 2015). (H-I) Clustergrams of: (H) All five WASH complex proteins identified by iBioID. (I) Previously reported WASH interactors (13/174), including the CCC and Retriever complexes. (J) Endosomal trafficking proteins (23/174 proteins). (K) Endocytic proteins (24/174). (L) Proteins involved in cytoskeletal regulation (32/174), including Arp2/3 subunit ARPC5. (M)Synaptic proteins (28/174). Clustergrams were annotated by hand and cross-referenced with Metascape (Zhou et al., 2019) GO enrichment of WASH1 proteome constituents over all proteins identified in the BioID experiment.

The resulting neuronal WASH proteome included 174 proteins that were significantly enriched (Fold-change ≥ 3.0, Benjamini-Hochberg P-Adjust < 0.1, Figure 1G). Of these proteins, we identified all five WASH complex components (Figure 1H), as well as 13 previously reported WASH complex interactors (Figure 1I) (McNally et al., 2017; Phillips-Krawczak et al., 2015; Simonetti and Cullen, 2019; Singla et al., 2019), which provided strong validity for our proteomic approach and analyses. Additional bioinformatic analyses of the neuronal WASH proteome identified a network of proteins implicated in vesicular trafficking, including 23 proteins enriched for endosomal functions (Figure 1J) and 24 proteins enriched for endocytic functions (Figure 1K). Among these endosomal and endocytic proteins were components of the recently identified endosomal sorting complexes, CCC (CCDC93 and COMMD9) and Retriever (VPS35L) (Phillips-Krawczak et al., 2015; Singla et al., 2019), as well as multiple sorting nexins important for recruitment of trafficking regulators to the endosome and cargo selection, such as SNX1-3, and SNX16 (Kvainickas et al., 2017; Maruzs et al., 2015; Simonetti et al., 2017). These data demonstrated that the WASH complex interacts with many of the same proteins in neurons as it does in yeast, amoebae, flies, and mammalian cell lines. Furthermore, there were 32 proteins enriched for cytoskeletal regulatory functions (Figure 1L), including actin-modulatory molecules such as the Arp2/3 complex subunit ARPC5, which is consistent with WASH’s role in activating this complex to stimulate actin polymerization at endosomes for vesicular scission (Billadeau et al., 2010; Derivery et al., 2009). The WASH1-BioID2 isolated complex also contained 28 proteins known to localize to the excitatory post-synapse (Figure 1M). This included many core synaptic scaffolding proteins, such as SHANK2-3 and DLGAP2-4 (Chen et al., 2011; Mao et al., 2015; Monteiro and Feng, 2017; Wan et al., 2011), as well as modulators of synaptic receptors such as SYNGAP1 and SHISA6 (Barnett et al., 2006; Clement et al., 2012; Kim et al., 2003; Klaassen et al., 2016), which was consistent with the idea that vesicular trafficking plays an important part in synaptic function and regulation. Taken together, these results support a major endosomal trafficking role of the WASH complex in mouse brain.

### SWIP^P1019R^ does not incorporate into the WASH complex, reducing its stability and levels *in vivo*

To determine how disruption of the WASH complex may lead to disease, we generated a mouse model of a human missense mutation found in children with intellectual disability, *WASHC4^c.3056c>g^* (protein: SWIP^P1019R^) (Ropers et al., 2011). Due to the sequence homology of human and mouse *Washc4* genes, we were able to introduce the same point mutation in exon 29 of murine *Washc4* using CRISPR (Derivery and Gautreau, 2010; Ropers et al., 2011). This C>G point mutation results in a Proline>Arginine substitution at position 1019 of SWIP’s amino acid sequence (Figure 2A), a region thought to be critical for its binding to the WASH component, Strumpellin (Jia et al., 2010; Ropers et al., 2011). Western blot analysis of brain lysate from adult homozygous SWIP^P1019R^ mutant mice (referred to from here on as MUT mice) displayed significantly decreased abundance of two WASH complex members, Strumpellin and WASH1 (Figure 2B). These results phenocopied data from the human patients (Ropers et al., 2011) and suggested that the WASH complex is unstable in the presence of this SWIP point mutation *in vivo*. To test whether this mutation disrupted interactions between WASH complex subunits, we compared the ability of wild-type SWIP (WT) and SWIP^P1019R^ (MUT) to co-immunoprecipitate with Strumpellin and WASH1 in HEK cells.

**Figure 2.**
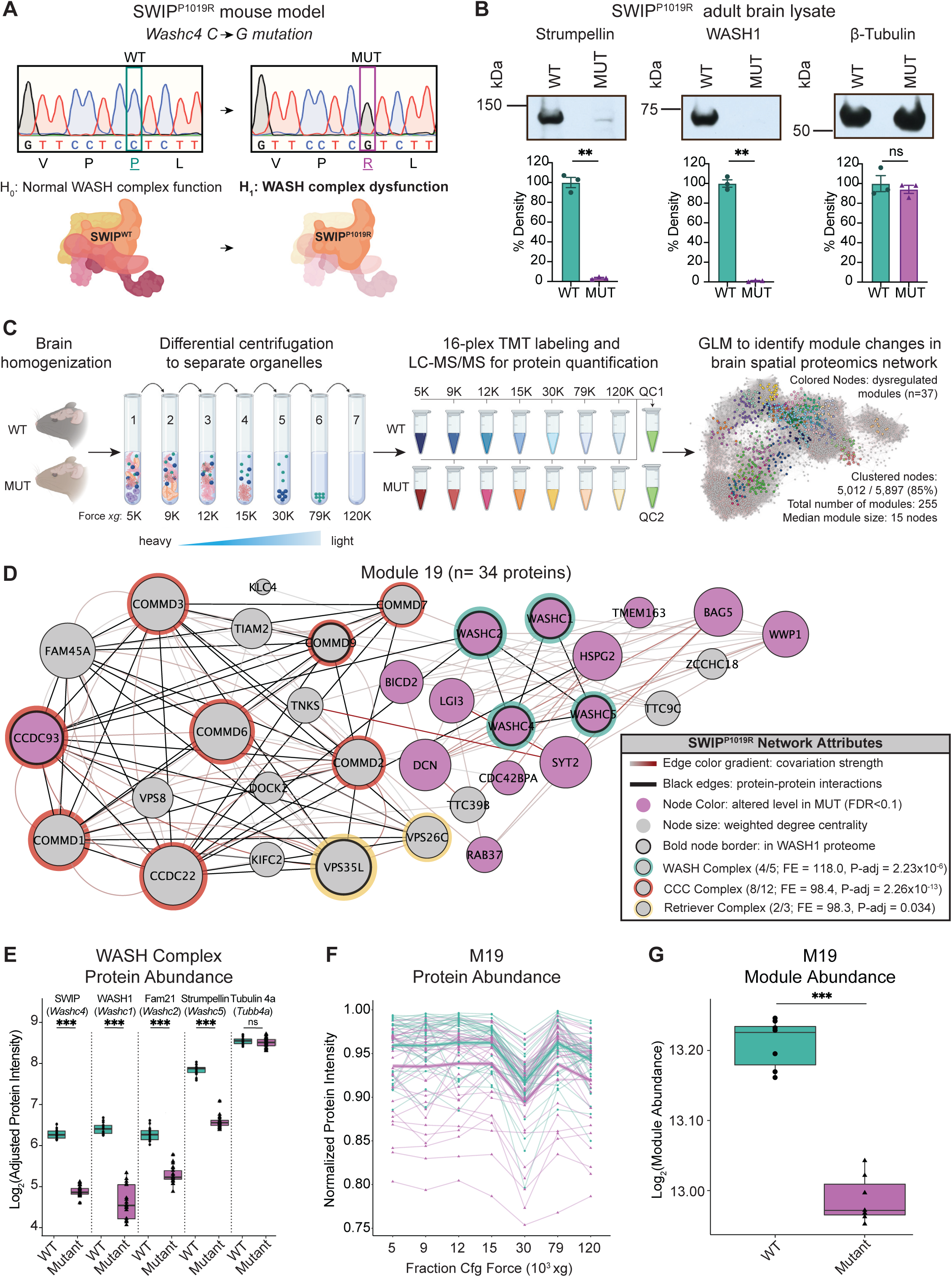
Spatial proteomics and network covariation analysis reveal significant disruptions to the WASH complex and an endosomal module in SWIP^P1019R^ mutant mouse brain. (A) Mouse model of the human SWIP^P1019R^ missense mutation created using CRISPR. A C>G point mutation was introduced into exon29 of murine *Washc4*, leading to a P1019R amino acid substitution. We hypothesize (H_1_) that this mutation causes instability of the WASH complex. (B) Representative western blot and quantification of WASH components, Strumpellin and WASH1 (predicted sizes in kDa: 134 and 72, respectively), as well as loading control β-Tubulin (55kDa) from whole adult whole brain lysate prepared from SWIP WT (*Washc4*^C/C^) and SWIP homozygous MUT (*Washc4*^G/G)^ mice. Bar plots show quantification of band intensities relative to WT (n=3 mice per genotype). Strumpellin (WT 100.0 ± 5.2%, MUT 3.5 ± 0.7%, t_2.1_=18.44, p=0.0024) and WASH1 (WT 100.0 ± 3.8%, MUT 1.1 ± 0.4%, t_2.1_=25.92, p=0.0013) were significantly decreased. Equivalent amounts of protein were analyzed in each condition (β-Tubulin: WT 100.0 ± 8.2%, MUT 94.1 ± 4.1%, U=4, p>0.99). (C) Spatial TMT proteomics experimental design. 7 subcellular fractions were prepared from one WT and one MUT mouse (10mo). These samples, as well as two pooled quality control (QC) samples, were labeled with unique TMT tags and concatenated for simultaneous LC-MS/MS analysis. This experiment was repeated three times (3 WT and 3 MUT brains total). To detect network-level changes, proteins were clustered into modules, and general linearized models (GLMs) were used to identify differences in module abundance between WT and MUT samples. The network shows an overview of the spatial proteomics graph in which the 37 differentially abundant modules are indicated by colored nodes. (D) Protein module 19 (M19) contains subunits of the WASH, CCC, and Retriever complexes. Node size denotes its weighted degree centrality (∼importance in module); purple node color indicates proteins with altered abundance in MUT brain relative to WT; black node border denotes proteins identified in the WASH1-BioID proteome (Figure 1); red, yellow, and green borders highlight protein components of the CCC, Retriever, and WASH complexes; black edges indicate known protein-protein interactions; and grey-red edges denote the relative strength of protein covariation within a module (gray = weak, red = strong). P-adjust values represent enrichment of proteins identified in the CORUM database adjusted for multiple comparisons (Giurgiu et al., 2019). (E) Difference in normalized protein abundance for four WASH proteins found in M19 (SWIP: WT 6.28 ± 0.41, MUT 4.89 ± 0.28, p=3.98×10^-28^; WASH1: WT 6.41 ± 0.47, MUT 4.65 ± 0.62, p=7.92×10^-18^; FAM21: WT 6.29 ± 0.48, MUT 5.29 ± 0.49, p=3.27×10^-16^; Strumpellin: WT 7.85 ± 0.52, MUT 6.59 ± 0.53, p=9.06×10^-25^) and one control (Tubulin 4a: WT 8.57 ± 0.52, MUT 8.52 ± 0.58, p>0.99) across all three experimental replicates (n=3 independent experiments), presented as log_2_(adjusted protein intensities). (F) Normalized average intensities for every protein within M19 across all seven subcellular fractions analyzed. Teal lines delineate protein levels in WT samples, purple lines delineate protein levels in MUT samples (averaged across three experimental replicates). Bolded lines demarcate the fitted intensity values for WT and MUT proteins (n=3 independent experiments). (G) Difference in M19 abundance, adjusted for fraction differences and presented as log_2_(adjusted module abundance) (WT 13.21 ± 0.003, MUT 12.99 ± 0.003, p=0.0007; n=3 independent experiments).

Compared to WT, MUT SWIP co-immunoprecipitated significantly less Strumpellin and WASH1 (IP: 54.8% and 41.4% of WT SWIP, respectively), suggesting that the SWIP^P1019R^ mutation hinders WASH complex formation (Figure 2-figure supplement 1). Together these data support the notion that SWIP^P1019R^ is a damaging mutation that not only impairs its function, but also results in significant reductions of the WASH complex as a whole.

### Spatial proteomics and unbiased network covariation analysis reveal significant disruptions in the endo-lysosomal pathway of SWIP^P1019R^ mutant mouse brain

Next, we aimed to understand the impact of the SWIP^P1019R^ mutation on the subcellular organization of the mouse brain proteome. We performed spatial proteomics by following the protocol established by Geladaki *et al*., with modifications for homogenization of brain tissue (Geladaki et al., 2019; Hallett et al., 2008). We isolated seven subcellular fractions from brain tissue and quantified proteins in these samples using 16-plex TMT proteomics. Using this spatial proteomics dataset, we developed a data-driven clustering approach to classify proteins into subcellular compartments. This approach, which differs from the support vector machine learning algorithm employed by Geladaki *et al*. (2019), was motivated by the lack of a large corpus of brain-specific protein subcellular localization information, and the greater complexity of brain tissue compared to cultured cells. In addition to evaluating differential protein abundance between WT and SWIP^P1019R^ MUT brain, we utilized this spatial proteomics dataset to analyze network-level changes in groups of covarying proteins to better understand WASH’s function and explore the cellular mechanisms by which SWIP^P1019R^ causes disease.

Brains from 10-month-old mice were gently homogenized to release intact organelles, followed by successive centrifugation steps to enrich subcellular compartments into different fractions based on their density (Figure 2C) (Geladaki et al., 2019). Seven WT and seven MUT fractions (each prepared from one brain, 14 samples total) were labeled with unique isobaric tandem-mass tags and concatenated. We also included two sample pooled quality controls (SPQCs), which allowed us to assess experimental variability and perform normalization between experiments. By performing this experiment in triplicate, deep coverage of the mouse brain proteome was obtained— across all 48 samples we quantified 86,551 peptides, corresponding to 7,488 proteins. After data pre-processing, normalization, and filtering we retained 5,897 reproducibly quantified proteins in the final dataset (Table S2).

We used generalized linear models (GLMs) to assess differential protein abundance for intra-fraction comparisons between WT and MUT genotypes, and for overall comparisons between WT and MUT groups, adjusted for baseline differences in subcellular fraction. In the first analysis, there were 85 proteins with significantly altered abundance in at least one of the 7 subcellular fractions (Benjamini-Hochberg P-Adjust < 0.1, Table S2 and Figure 2-figure supplement 2). Five proteins were differentially abundant between WT and MUT in all 7 fractions, including four WASH proteins and RAB21A—a known WASH interactor that functions in early endosomal trafficking (WASHC1, WASHC2, WASHC4, WASHC5, Figure 2E) (Del Olmo et al., 2019; Simpson et al., 2004). The abundance of the remaining WASH complex protein, WASHC3, was found to be very low and was not retained in the final dataset due to its sparse quantification. These data affirm that the SWIP^P1019R^ mutation destabilizes the WASH complex. Next, to evaluate global differences between WT and MUT brain, we analyzed the average effect of genotype on protein abundance across all fractions. At this level, there were 687 differentially abundant proteins between WT and MUT brain (Bonferroni P-Adjust < 0.05) (Table S2). We then aimed to place these differentially abundant proteins into a more meaningful biological context using a systems-based approach.

For network-based analyses, we clustered the protein covariation network defined by pairwise correlations between all 5,897 proteins. Our data-driven, quality-based approach used Network Enhancement (Wang et al., 2018) to remove biological noise from the covariation network and employed the Leiden algorithm (Traag et al., 2019) to identify optimal partitions of the graph. We enforced module quality by permutation testing (Ritchie et al., 2016) to ensure that identified modules exhibited a non-random topology. Clustering of the protein covariation graph identified 255 modules of proteins that strongly covaried together (see Methods for complete description of clustering approach).

To test for module-level differences between WT and MUT brain, we summarized modules for each biological replicate (a single subcellular fraction prepared from either a WT or MUT mouse) as the sum of their proteins, and extended our GLM framework to identify changes in module abundance (adjusted for fraction differences) between genotypes. 37 of the 255 modules exhibited significant differences in WT versus MUT brain (Bonferroni P-Adjust < 0.05; Table S3). Of note, the module containing the WASH complex, M19, was predicted to have endosomal function by annotation of protein function, and was enriched for proteins identified by WASH1-BioID2 (hypergeometric test P-Adjust < 0.05, bold node edges, Figure 2D). Similar to the WASH iBioID proteome (Figure 1), M19 contained components of the CCC (CCDC22, CCDC93, COMMD1-3, COMMD6-7, and COMMD9) and Retriever sorting complexes (VPS26C and VPS35L), but not the Retromer sorting complex, suggesting that in the brain, the WASH complex may not interact as closely with Retromer as it does in other cells (Figure 2D). Across all fractions, the abundance of M19 was significantly lower in MUT brain compared to WT, providing evidence that the SWIP^P1019R^ mutation reduces the stability of this protein subnetwork and impairs its function (Figure 2F-G).

In contrast to the decreased abundance of the WASH complex/endosome module, M19, we observed three modules (M2, M159, and M213) which were enriched for lysosomal protein components (Geladaki et al., 2019), and exhibited increased abundance in MUT brain (Figure 3). M159 (Figure 3B) contained the lysosomal protease Cathepsin A (CTSA), while M213 (Figure 3D) contained Cathepsin B (CTSB), as well as two key lysosomal hydrolases GLB1 and MAN2B2, and M2 (Figure 3C) contained two Cathepsins (CTSS and CTSL) and several lysosomal hydrolases (e.g. GNS, GLA, and MAN2B1) (Eng and Desnick, 1994; Mayor et al., 1993; Mok et al., 2003; Moon et al., 2016; Patel et al., 2018; Regier and Tifft, 1993; Rosenbaum et al., 2014). Notably, M2 also contained the lysosomal glycoprotein progranulin (GRN), which is integral to proper lysosome function and whose loss is widely linked with neurodegenerative pathologies (Baker et al., 2006; Pottier et al., 2016; Tanaka et al., 2017; Zhou et al., 2018). In addition, M2 contained the hydrolase IDS, whose loss causes a lysosomal storage disorder that can present with neurological symptoms (Hopwood et al., 1993; Schröder et al., 1994). The overall increase in abundance of modules M2, M159, and M213, and these key lysosomal proteins (Figure 3E-G), may therefore reflect an increase in flux through degradative lysosomal pathways in SWIP^P1019R^ brain.

**Figure 3.**
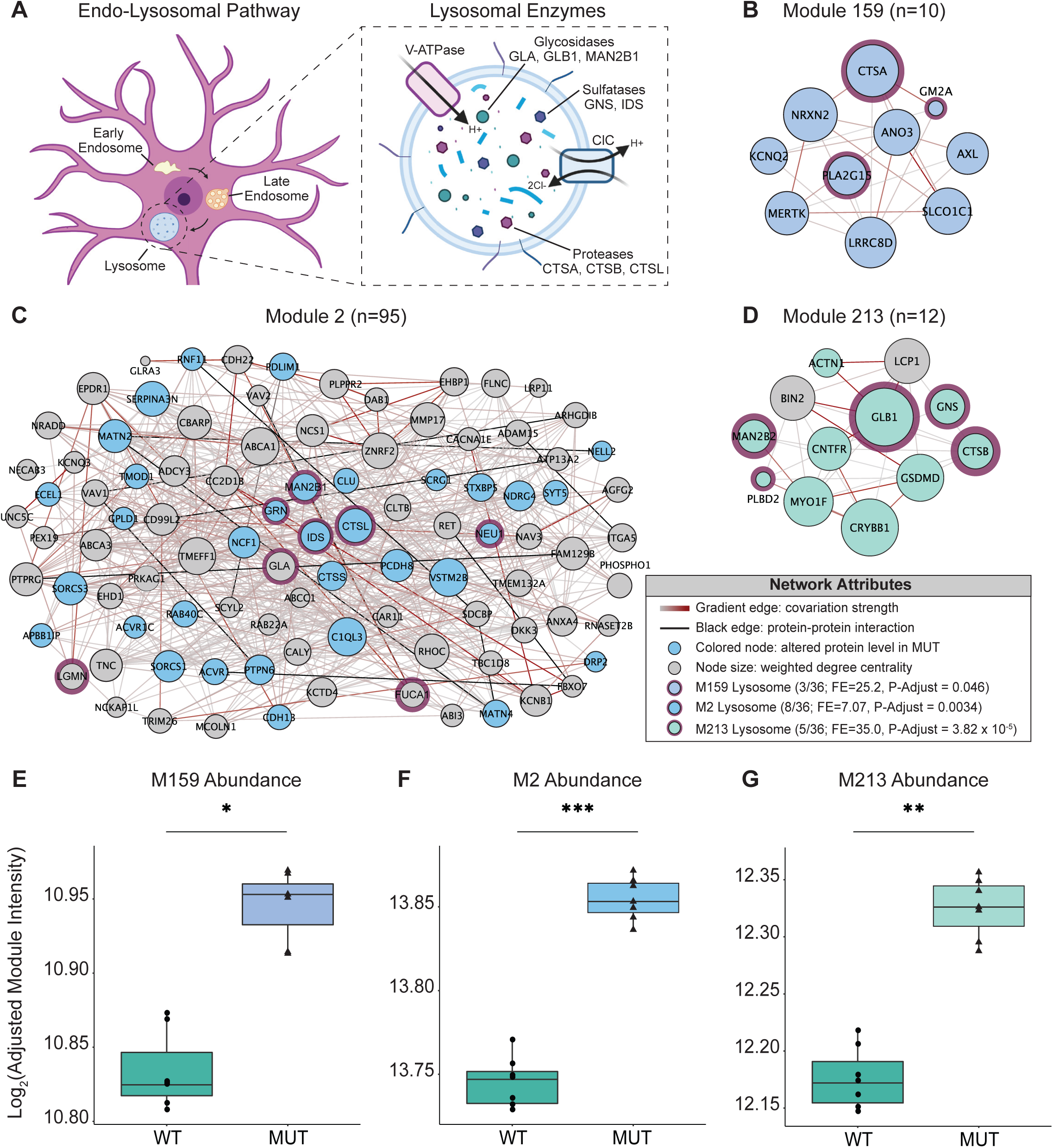
Disruption of lysosomal protein networks in SWIP^P1019R^ mutant brain. (A) Simplified schematic of the endo-lysosomal pathway in neurons. Inset depicts representative lysosomal enzymes, such as proteases (CTSA, CTSB, CTSL), glycosidases (GLA, GLB1, MAN2B1), and sulfatases (GNS, IDS). (B) Network graph of module 159 (M159). All proteins in M159 exhibit altered abundance in MUT brain, including lysosomal proteins, CTSA, PLA2G15, and GM2A. (C) Module 2 (M2) contains multiple lysosomal proteins with increased abundance in MUT brain compared to WT, including CTSS, CTSL, GRN, IDS, MAN2B1. (D) Module 213 (M213) contains multiple proteins with increased abundance in MUT brain, including lysosomal proteins, GLB1, GNS, CTSB, MAN2B2, and PLBD2. Network attributes (B-D): Node size denotes its weighted degree centrality (∼importance in module), node color indicates proteins with altered abundance in MUT brain relative to WT, purple outline highlight proteins identified as lysosomal in (Geladaki et al., 2019), black edges indicate known protein-protein interactions, and grey-red edges denote the relative strength of protein covariation within a module (gray = weak, red = strong). P-Adjust values represent enrichment of proteins identified as lysosomal in Geladaki et al., 2019. (E) The overall effect of genotype on M159 module abundance (WT 10.83 ± 0.002, MUT 10.94 ± 0.002, p=0.031). (F) The overall effect of genotype on M2 module abundance (WT 13.74 ± 0.001, MUT 13.85 ± 0.0009, p=0.0006). (G) The overall effect of genotype on M213 abundance (WT 12.17 ± 0.002, MUT 12.33 ± 0.002, p=0.0037). Data reported as mean ± SEM, error bars are SEM. *p<0.05, ** p<0.01, ***p<0.001, empirical Bayes quasi-likelihood F-test with Bonferroni correction (E-G).

Furthermore, Module 2 (Figure 3C) included multiple membrane proteins and extracellular proteins, such as ITGA5 (an integrin shown to be upregulated and redistributed upon loss of WASH1), ATP13A2 (a cation transporter whose loss causes a Parkinsonian syndrome), and MMP17 (an extracellular metalloprotease), suggesting a link between these proteins and lysosomal enzymatic function (English et al., 2000; Ramirez et al., 2006; Zech et al., 2011). Increased abundance of these M2 proteins in MUT brain may indicate that WASH complex disruption alters their cellular localization. Taken together, these changes appear to reflect a pathological condition characterized by distorted lysosomal metabolism and altered cellular trafficking.

In addition to these endo-lysosomal changes, network alterations were evident for an endoplasmic reticulum (ER) module (M83), supporting a shift in the proteostasis of mutant neurons (Figure 2-figure supplement 3B). Notably, within the ER module, M83, there was increased abundance of chaperones (e.g. HSPA5, PDIA3, PDIA4, PDIA6, and DNAJC3) that are commonly engaged in presence of misfolded proteins (Bartels et al., 2019; Kim et al., 2020; Montibeller and de Belleroche, 2018; Synofzik et al., 2014; Wang et al., 2016). This elevation of ER stress modulators can be indicative of neurodegenerative states, in which the unfolded protein response (UPR) is activated to resolve misfolded species (Garcia-Huerta et al., 2016; Hetz and Saxena, 2017). These data demonstrate that loss of WASH function not only alters endo-lysosomal trafficking, but also causes increased stress on cellular homeostasis.

Finally, besides these endo-lysosomal and homeostatic changes, we also observed two synaptic modules (M35 and M248) that were reduced in MUT brain (Figure 2-figure supplement 3C-D). These included mostly excitatory post-synaptic proteins such as HOMER2 and DLG4 (also identified in WASH1-BioID, Figure 1), consistent with endosomal WASH influencing synaptic regulation. Decreased abundance of these modules indicates that loss of the WASH complex may result in failure of these proteins to be properly trafficked to the synapse.

### Mutant neurons display structural abnormalities in endo-lysosomal compartments *in vitro*

Combined, the proteomics data strongly suggested that endo-lysosomal pathways are altered in adult SWIP^P1019R^ mutant mouse brain. Next, we analyzed whether structural changes in this system were evident in primary neurons. Cortical neurons from littermate WT and MUT P0 pups were cultured for 15 days *in vitro* (DIV15, Figure 4A), then fixed and stained for established markers of early endosomes (Early Endosome Antigen 1, EEA1; Figures 4B and 4C) and lysosomes (Cathepsin D, CathD; Figures 4D and 4E). Reconstructed three-dimensional volumes of EEA1 and Cathepsin D puncta revealed that MUT neurons display larger EEA1+ somatic puncta than WT neurons (Figures 4G and 4J), but no difference in the total number of EEA1+ puncta (Figure 4F). This finding is consistent with a loss-of-function mutation, as loss of WASH activity prevents cargo scission from endosomes and leads to cargo accumulation (Bartuzi et al., 2016; Gomez et al., 2012). Conversely, MUT neurons exhibited significantly less Cathepsin D+ puncta than WT neurons (Figure 4H), but the remaining puncta were significantly larger than those of WT neurons (Figures 4I and 4K). These data support the finding that the SWIP^P1019R^ mutation results in both molecular and morphological abnormalities in the endo-lysosomal pathway.

**Figure 4.**
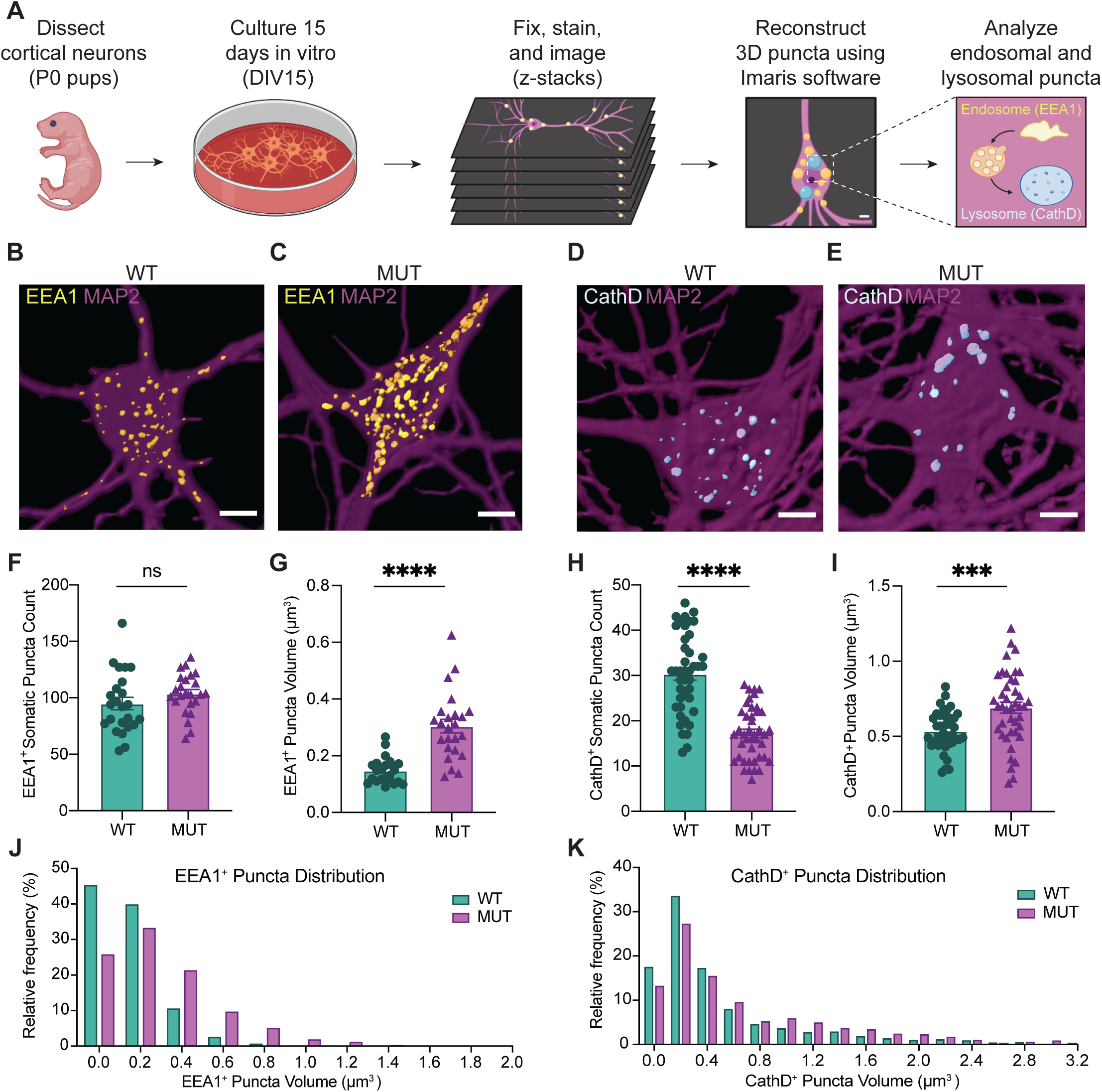
SWIP^P1019R^ mutant neurons display structural abnormalities in endo-lysosomal compartments *in vitro*. (A) Experimental design. Cortices were dissected from P0 pups, and neurons were dissociated and cultured on glass coverslips for 15 days. Cultures were fixed, stained, and imaged using confocal microscopy. 3D puncta volumes were reconstructed from z-stack images using Imaris software. (B-C) Representative 3D reconstructions of WT and MUT DIV15 neurons (respectively) stained for EEA1 (yellow) and MAP2 (magenta). (D-E) Representative 3D reconstructions of WT and MUT DIV15 neurons (respectively) stained for Cathepsin D (cyan) and MAP2 (magenta). (F) Graph of the average number of EEA1+ volumes per soma in each image (WT 95.0 ± 5.5, n=24 neurons; MUT 103.7 ± 3.7, n=24 neurons; t_40.2_=1.314, p=0.1961). (G) Graph of the average EEA1+ volume size per soma shows larger EEA1+ volumes in MUT neurons (WT 0.15 ± 0.01 µm^3^, n=24 neurons; MUT 0.30 ± 0.02 µm^3^, n=24 neurons; U=50, p<0.0001). (H) Graph of the average number of Cathepsin D+ volumes per soma illustrates less Cathepsin D+ volumes in MUT neurons (WT 30.4 ± 1.4, n=42; MUT 17.2 ± 0.9, n=42; t_71_=7.943, p<0.0001). (I) Graph of the average Cathepsin D+ volume size per soma demonstrates larger Cathepsin D+ volumes in MUT neurons (WT 0.54 ± 0.02 µm^3^, n=42; MUT 0.69 ± 0.04 µm^3^, n=42; t_63_=3.701, p=0.0005). (J) Histogram of EEA1+ volumes illustrate differences in size distributions between MUT and WT neurons. (K) Histogram of CathD+ volumes show differences in size distributions between MUT and WT neurons. Analyses included at least three separate culture preparations. Scale bars, 5 µm (B-E). Data reported as mean ± SEM, error bars are SEM. ***p<0.001, ****p<0.0001, two-tailed t-tests or Mann-Whitney U test (G).

### SWIP^P1019R^ mutant brains exhibit markers of abnormal endo-lysosomal structures and cell death *in vivo*

As there is strong evidence that dysfunctional endo-lysosomal trafficking and elevated ER stress are associated with neurodegenerative disorders, adolescent (P42) and adult (10 month-old, 10mo) WT and MUT brain tissue were analyzed for the presence of cleaved caspase-3, a marker of apoptotic pathway activation, in four brain regions (Boatright and Salvesen, 2003; Porter and Jänicke, 1999). Very little cleaved caspase-3 staining was present in WT and MUT mice at adolescence (Figures 5A, 5B, and Figure 5-figure supplement 1). However, at 10mo, the MUT motor cortices displayed significantly greater cleaved caspsase-3 staining compared to age-matched WT littermate controls (Figures 5D, 5E, and 5H). Furthermore, this difference appeared to be selective for the motor cortex, as we did not observe significant differences in cleaved caspase-3 staining at either age for hippocampal, striatal, or cerebellar regions (Figure 5-figure supplement 1). These data suggested that neurons of the motor cortex were particularly susceptible to disruption of endo-lysosomal pathways downstream of SWIP^P109R^, perhaps because long-range corticospinal projections require high fidelity of trafficking pathways (Blackstone et al., 2011; Slosarek et al., 2018; Wang et al., 2014).

**Figure 5.**
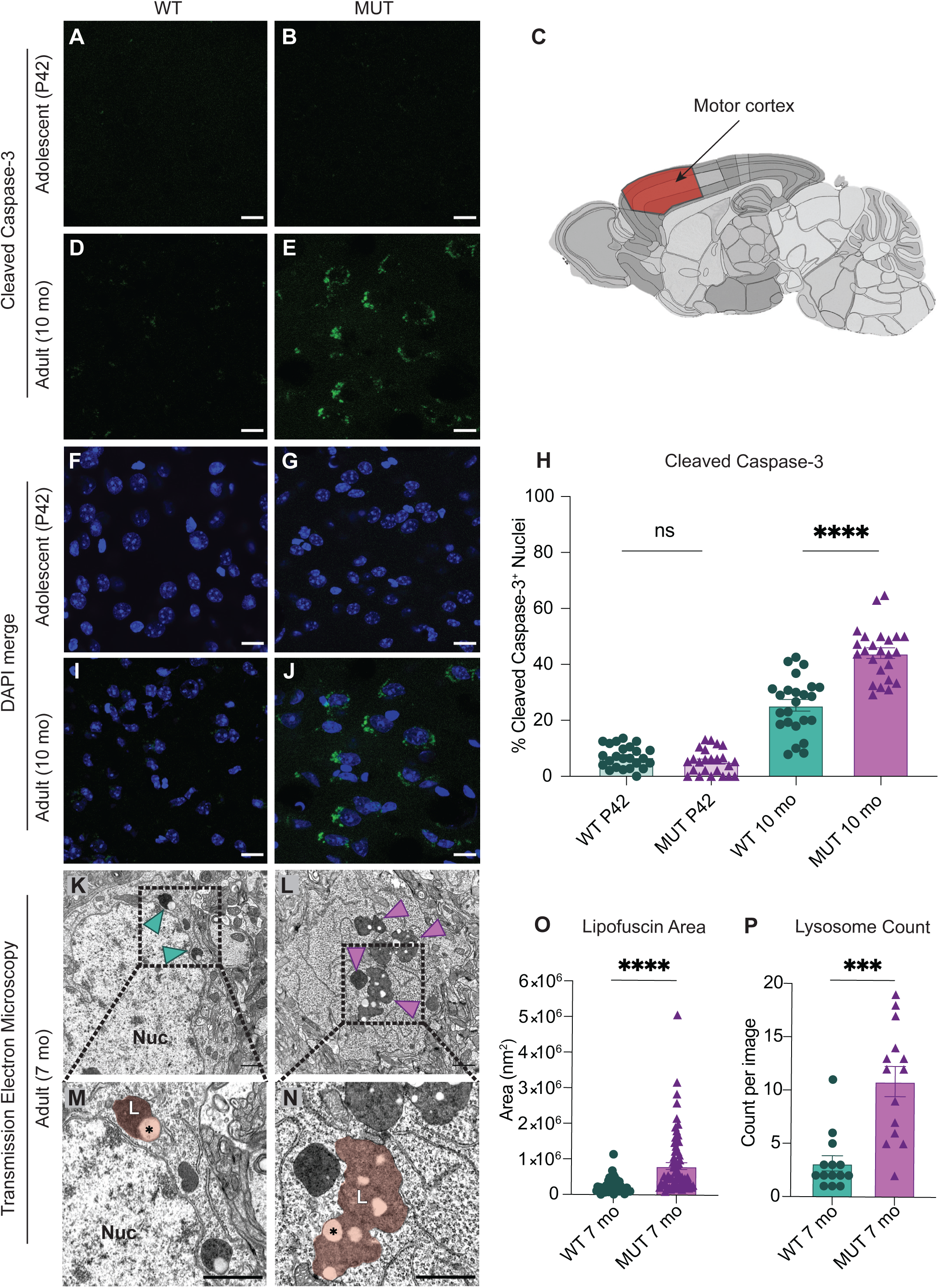
SWIP^P1019R^ mutant brains exhibit markers of abnormal endo-lysosomal structures and cell death *in vivo*. (A-B) Representative images of adolescent (P42) WT and MUT motor cortex stained with cleaved caspase-3 (CC3, green). (C) Anatomical representation of mouse brain with motor cortex highlighted in red, adapted from the Allen Brain Atlas (Oh et al., 2014). (D-E) Representative image of adult (10 mo) WT and MUT motor cortex stained with CC3 (green). (F, G, I, and J) DAPI co-stained images for (A, B, D, and E, respectively). Scale bar for (A-J), 15 µm. (H) Graph depicting the normalized percentage of DAPI+ nuclei that are positive for CC3 per image. No difference is seen at P42, but the amount of CC3+ nuclei is significantly higher in aged MUT mice (P42 WT 6.97 ± 0.80%, P42 MUT 5.26 ± 0.90%, 10mo WT 25.38 ± 2.05%, 10mo MUT 44.01 ± 1.90%, n=24 images per genotype taken from 4 different mice, H=74.12, p<0.0001). We observed no difference in number of nuclei per image between genotypes. (K) Representative transmission electron microscopy (TEM) image taken of soma from adult (7mo) WT motor cortex. Arrowheads delineate electron-dense lipofuscin material, Nuc = nucleus. (L) Representative transmission electron microscopy (TEM) image taken of soma from adult (7mo) MUT motor cortex. (M) Inset from (K) highlights lysosomal structure in WT soma. Pseudo-colored region depicts lipofuscin area, demarcated as L. (N) Inset from (L) highlights large lipofuscin deposit in MUT soma (L, pseudo-colored region) with electron-dense and electron-lucent lipid-like (asterisk) components. (O) Graph of areas of electron-dense regions of interest (ROI) shows increased ROI size in MUT neurons (WT 2.4×10^5^ ± 2.8×10^4^ nm^2^, n=50 ROIs; MUT 8.2×10^5^ ± 9.7 ×10^4^ nm^2^, n=75 ROIs; U=636, p<0.0001). (P) Graph of the average number of presumptive lysosomes with associated electron-dense material reveals increased number in MUT samples (WT 3.14 ± 0.72 ROIs, n=14 images; MUT 10.86 ± 1.42 ROIs, n=14 images; U=17, p<0.0001). For (N) and (O), images were taken from multiple TEM grids, prepared from n=3 animals per genotype. Scale bar for all TEM images, 1 µm. Data reported as mean ± SEM, error bars are SEM. ***p<0.001, ****p<0.0001, Kruskal-Wallis test (F), Mann-Whitney U test (O-P).

To further examine the morphology of primary motor cortex neurons at a subcellular resolution, samples from age-matched 7-month-old WT and MUT mice (7mo, 3 animals each) were imaged by transmission electron microscopy (TEM). Strikingly, we observed large electron-dense inclusions in the cell bodies of MUT neurons (arrows, Figure 5L; pseudo-colored region, 5N). These dense structures were associated electron-lucent lipid-like inclusions (asterisk, Figure 5N), and were visually consistent with lipofuscin accumulation at lysosomal residual bodies (Poët et al., 2006; Valdez et al., 2017; Yoshikawa et al., 2002). Lipofuscin is a by-product of lysosomal breakdown of lipids, proteins, and carbohydrates, which naturally accumulates over time in non-dividing cells such as neurons (Höhn and Grune, 2013; Moreno-García et al., 2018; Terman and Brunk, 1998). However, excessive lipofuscin accumulation is thought to be detrimental to cellular homeostasis by inhibiting lysosomal function and promoting oxidative stress, often leading to cell death (Brunk and Terman, 2002; Powell et al., 2005). As a result, elevated lipofuscin is considered a biomarker of neurodegenerative disorders, including Alzheimer’s disease, Parkinson’s disease, and Neuronal Ceroid Lipofuscinoses (Moreno-García et al., 2018). Therefore, the marked increase in lipofuscin area and number seen in MUT electron micrographs (Figures 5O and 5P, respectively) is consistent with the increased abundance of lysosomal pathways observed by proteomics, and likely reflects an increase in lysosomal breakdown of cellular material. Together these data indicate that SWIP^P1019R^ results in pathological lysosomal function that could lead to neurodegeneration.

### SWIP^P1019R^ mutant mice display persistent deficits in cued fear memory recall

To observe the functional consequences of the SWIP^P1019R^ mutation, we next studied WT and MUT mouse behavior. Given that children with homozygous SWIP^P1019R^ point mutations display intellectual disability (Ropers et al., 2011) and SWIP^P1019R^ mutant mice exhibit endo-lysosomal disruptions implicated in neurodegenerative processes, behavior was assessed at two ages: adolescence (P40-50), and mid-late adulthood (5.5-6.5 mo). Interestingly, MUT mice performed equivalently to WT mice in episodic and working memory paradigms, including novel object recognition and Y-maze alternations (Figure 6-figure supplement 1). However, in a fear conditioning task, MUT mice displayed a significant deficit in cued fear memory (Figure 6). This task tests the ability of a mouse to associate an aversive event (a mild electric footshock) with a paired tone (Figure 6A). Freezing behavior of mice during tone presentation is attributed to hippocampal or amygdala-based fear memory processes (Goosens and Maren, 2001; Maren and Holt, 2000; Vazdarjanova and McGaugh, 1998). Forty-eight hours after exposure to the paired tone and footshock, MUT mice showed a significant decrease in conditioned freezing to tone presentation compared to their WT littermates (Figures 6B and 6C). To ensure that this difference was not due to altered sensory capacities of MUT mice, we measured the startle response of mice to both electric foot shock and presented tones. In line with intact sensation, MUT mice responded comparably to WT mice in these tests (Figure 6-figure supplement 2). These data demonstrate that although MUT mice perceive footshock sensations and auditory cues, it is their memory of these paired events that is significantly impaired. Additionally, this deficit in fear response was evident at both adolescence and adulthood (top panels, and bottom panels, respectively, Figures 6B and 6C). These changes are consistent with the hypothesis that SWIP^P109R^ is the cause of cognitive impairments in humans.

**Figure 6.**
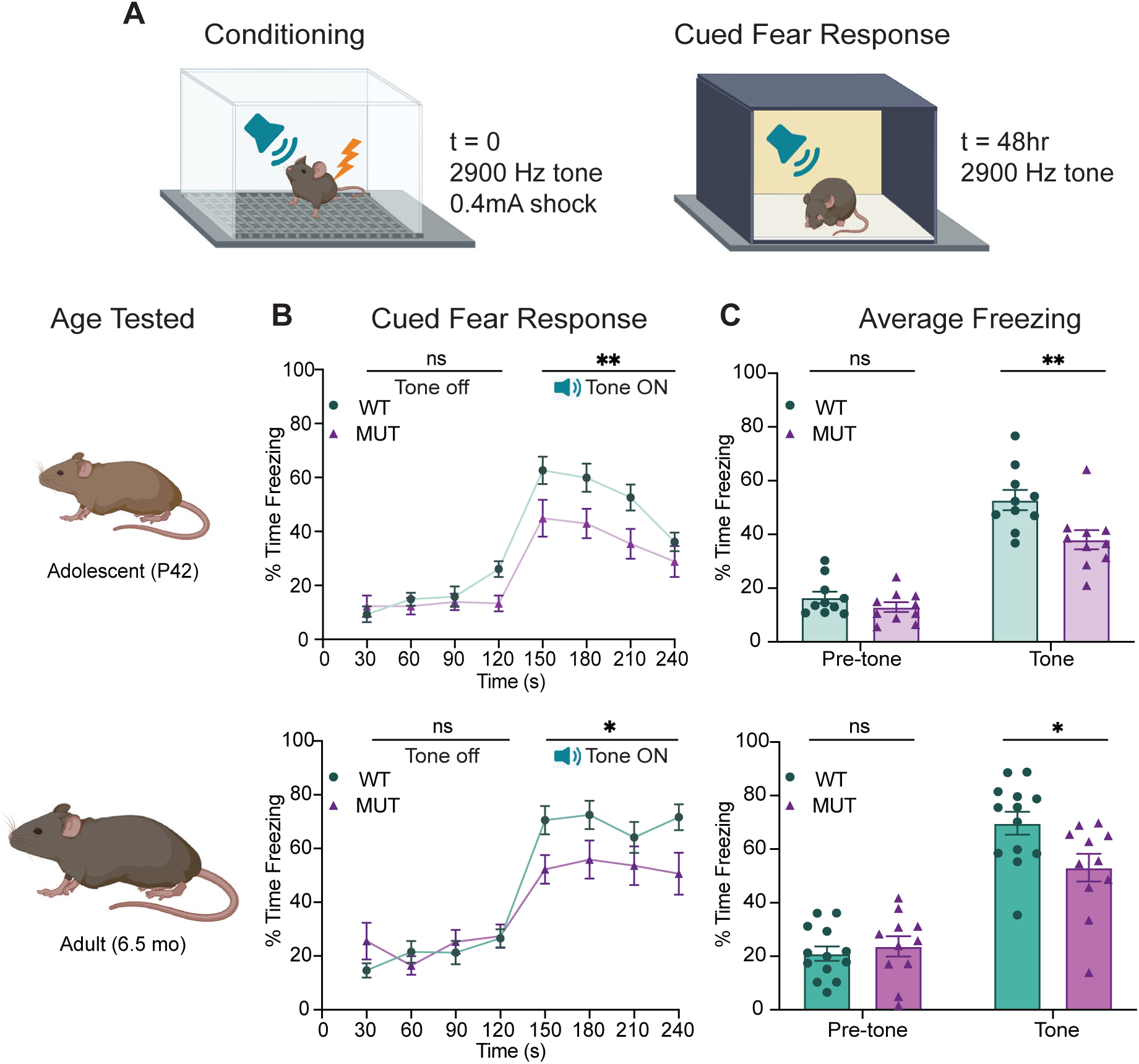
SWIP^P1019R^ mutant mice display persistent deficits in cued fear memory recall. (A) Experimental fear conditioning paradigm. After acclimation to a conditioning chamber, mice received a mild aversive 0.4mA footshock paired with a 2900Hz tone. 48 hours later, the mice were placed in a chamber with different tactile and visual cues. The mice acclimated for two minutes and then the 2900Hz tone was played (no footshock) and freezing behavior was assessed. (B) Line graphs of WT and MUT freezing response during cued tone memory recall. Data represented as average freezing per genotype in 30 s time bins. The tone is presented after t = 120 s, and remains on for 120 seconds (Tone ON). Two different cohorts of mice were used for age groups P42 (top) and 6.5mo (bottom). Two-way ANOVA analysis of average freezing during Pre-Tone and Tone periods reveal a Genotype x Time effect at P42 (WT n=10, MUT n=10, F_1,18_=4.944, p=0.0392) and 6.5mo (WT n=13, MUT n=11, F_1,22_= 13.61, p=0.0013). (C) Graphs showing the average %time freezing per animal before and during tone presentation. Top: freezing is reduced by 20% in MUT adolescent mice compared to WT littermates (Pre-tone WT 16.5 ± 2.2%, n=10; Pre-tone MUT 13.0 ± 1.8%, n=10; t_36_=0.8569, p=0.6366; Tone WT 52.8 ± 3.8%, n=10; Tone MUT 38.0 ± 3.6%, n=10; t_36_=3.539, p=0.0023), Bottom: freezing is reduced by over 30% in MUT adult mice compared to WT littermates (Pre-tone WT 21.1 ± 2.7%, n=13; Pre-tone MUT 23.7 ± 3.8%, n=11; t_44_=0.4675, p=0.8721; Tone WT 69.7 ± 4.3%, n=13; Tone MUT 53.1 ± 5.2%, n=11; t_44_=2.921, p=0.0109). Data reported as mean ± SEM, error bars are SEM. *p<0.05, **p<0.01, two-way ANOVAs (B) and Sidak’s post-hoc analyses (C).

### SWIP^P1019R^ mutant mice exhibit surprising motor deficits that are confirmed in human patients

Because SWIP^P1019R^ results in endo-lysosomal pathology consistent with neurodegenerative disorders in the motor cortex, we next analyzed motor function of the mice over time. First, we tested the ability of WT and MUT mice to remain on a rotating rod for five minutes (Rotarod, Figures 7A-7C). At both adolescence and adulthood, MUT mice performed markedly worse than WT littermate controls (Fig 7C). Mouse performance was not significantly different across trials, which suggested that this difference in retention time was not due to progressive fatigue, but more likely due to an overall difference in motor control (Mann and Chesselet, 2015).

**Figure 7.**
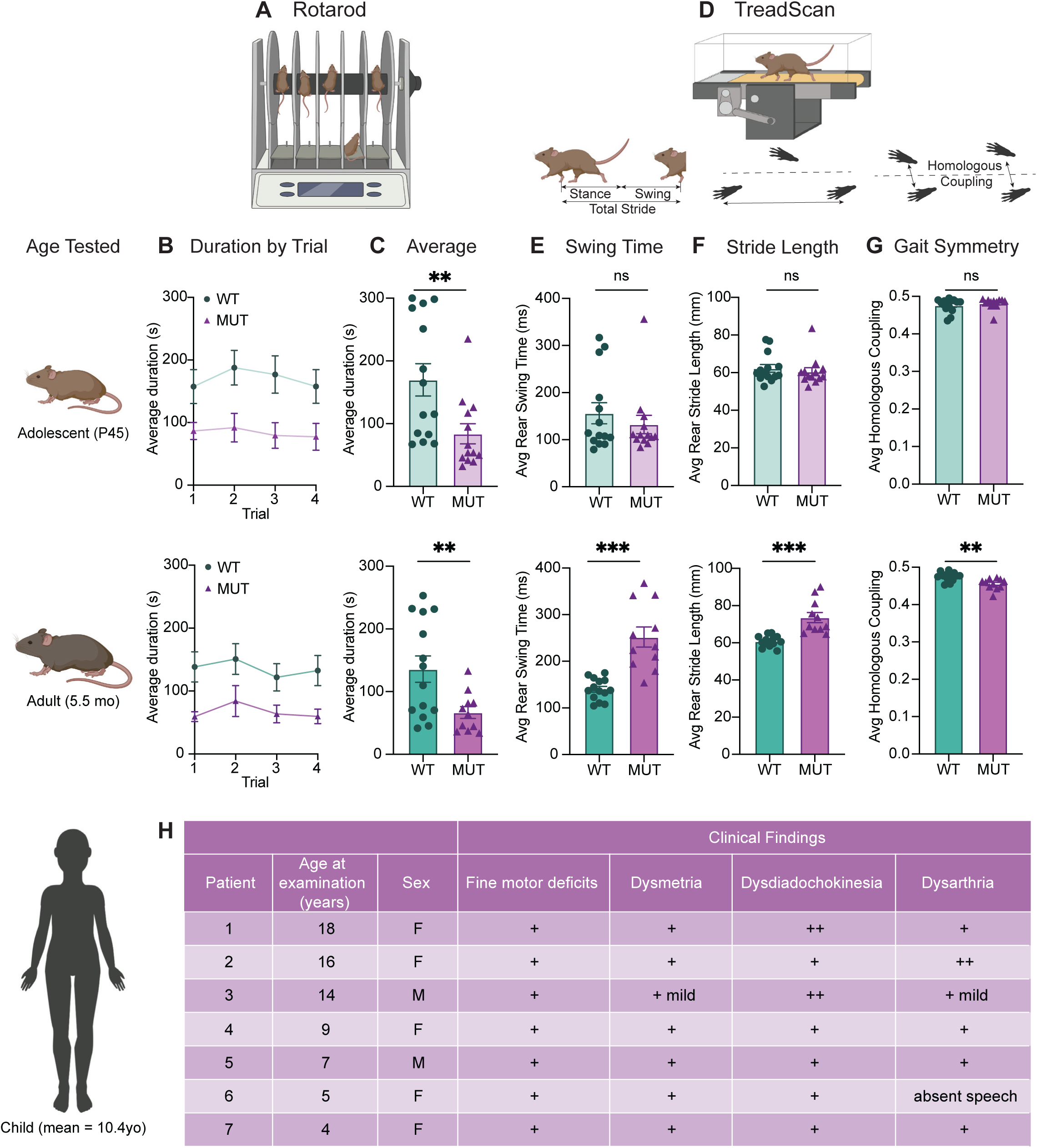
SWIP^P1019R^ mutant mice exhibit surprising motor deficits that are confirmed in human patients. (A) Rotarod experimental setup. Mice walked atop a rod rotating at 32rpm for 5 minutes, and the duration of time they remained on the rod before falling was recorded. (B) Line graph of average duration animals remained on the rod per genotype across four trials, with an inter-trial interval of 40 minutes. The same cohort of animals was tested at two different ages, P45 (top) and 5.5 months (bottom). Genotype had a significant effect on task performance at both ages (top, P45: genotype effect, F_1,25_=7.821, p=0.0098. bottom, 5.5mo: genotype effect, F_1,23_= 7.573, p=0.0114). (C) Graphs showing the average duration each animal remained on the rod across trials. At both ages, the MUT mice exhibited an almost 50% reduction in their ability to remain on the rod (Top, P45: WT 169.9 ± 25.7 s, MUT 83.8 ± 15.9 s, U=35, p=0.0054. Bottom, 5.5mo: WT 135.9 ± 20.9 s, MUT 66.7 ± 9.5 s, t_18_=3.011, p=0.0075). (D) TreadScan task. Mice walked on a treadmill for 20 s while their gate was captured with a high-speed camera. Diagrams of gait parameters measured in (E-G) are shown below the TreadScan apparatus. (E) Average swing time per stride for hindlimbs. At P45 (top), there is no significant difference in rear swing time (WT 156.2 ± 22.4 ms, MUT 132.3 ± 19.6 ms, U=83, p=0.7203). At 5.5mo (bottom), MUT mice display significantly longer rear swing time (WT 140 ± 6.2 ms, MUT 252.0 ± 21.6 ms, t_12_=4.988, p=0.0003). (F) Average stride length for hindlimbs. At P45 (top), there is no significant difference in stride length (WT 62.3 ± 2.0 mm, MUT 60.5 ± 2.1 mm, U=75, p=0.4583). At 5.5mo (bottom), MUT mice take significantly longer strides with their hindlimbs (WT 60.8 ± 0.8 mm, MUT 73.6 ± 2.7 mm, t_11.7_=4.547, p=0.0007). (G) Average homologous coupling for front and rear limbs. Homologous coupling is 0.5 when the left and right feet are completely out of phase. At P45 (top), WT and MUT mice exhibit normal homologous coupling (WT 0.48 ± 0.005, MUT 0.48 ± 0.004, U=76.5, p=0.4920). At 5.5 mo (bottom), MUT mice display decreased homologous coupling, suggestive of abnormal gait symmetry (WT 0.48 ± 0.003, MUT 0.46 ± 0.004, t_18.8_=3.715, p=0.0015). At P45: n=14 WT, n=13 MUT; At 5.5mo: n=14 WT, n=11 MUT. (H) Table of motor findings in clinical exam of human patients with the homozygous SWIP^P1019R^ mutation (n=7). All patients exhibit motor dysfunction (+ = symptom present). Data reported as mean ± SEM, error bars are SEM.*p<0.05, **p<0.01, ***p<0.001, ****p<0.0001, two-way repeated measure ANOVAs (B), Mann-Whitney U tests and two-tailed t-tests (C-G).

To study the animals’ movement at a finer scale, the gait of WT and MUT mice was also analyzed using a TreadScan system containing a high-speed camera coupled to a transparent treadmill (Figure 7D) (Beare et al., 2009). Interestingly, while the gait parameters of mice were largely indistinguishable across genotypes at adolescence, a striking difference was seen when the same mice were aged to adulthood (Figures 7E-7G). In particular, MUT mice took slower (Figure 7E), longer strides (Figure 7F), stepping closer to the midline of their body (track width, Figure 7-figure supplement 1), and their gait symmetry was altered so that their strides were no longer perfectly out of phase (out of phase=0.5, Figure 7G). While these differences were most pronounced in the rear limbs (as depicted in Figure 7E-7G), the same trends were present in front limbs (Figure 7-figure supplement 1). These findings demonstrate that SWIP^P1019R^ results in progressive motor function decline that was detectable by the rotarod task at adolescence, but which became more prominent with age, as both gait and strength functions deteriorated.

These marked motor findings prompted us to re-evaluate the original reports of human SWIP^P1019R^ patients (Ropers et al., 2011). While developmental delay or learning difficulties were the primary impetus for medical evaluation, all patients also exhibited motor symptoms (mean age = 10.4 years old, Figure 7H). The patients’ movements were described as “clumsy” with notable fine motor difficulties, dysmetria, dysdiadochokinesia, and mild dysarthria on clinical exam (Figure 7H). Recent communication with the parents of these patients, who are now an average of 21 years old, revealed no notable symptom exacerbation. It is therefore possible that the SWIP^P1019R^ mouse model either exhibits differences from human patients or may predict future disease progression for these individuals, given that we observed significant worsening at 5-6 months old in mice (which is thought to be equivalent to ∼30-35 years old in humans) (Dutta and Sengupta, 2016; Zhang et al., 2019).

## DISSCUSSION

Taken together, the data presented here support a mechanistic model whereby SWIP^P1019R^ causes a loss of WASH complex function, resulting in endo-lysosomal disruption and accumulation of neurodegenerative markers, such as upregulation of unfolded protein response modulators and lysosomal enzymes, as well as build-up of lipofuscin and cleaved caspase-3 over time. To our knowledge, this study provides the first mechanistic evidence of WASH complex impairment having direct and indirect organellar effects that lead to cognitive deficits and progressive motor impairments (Figure 8).

**Figure 8.**
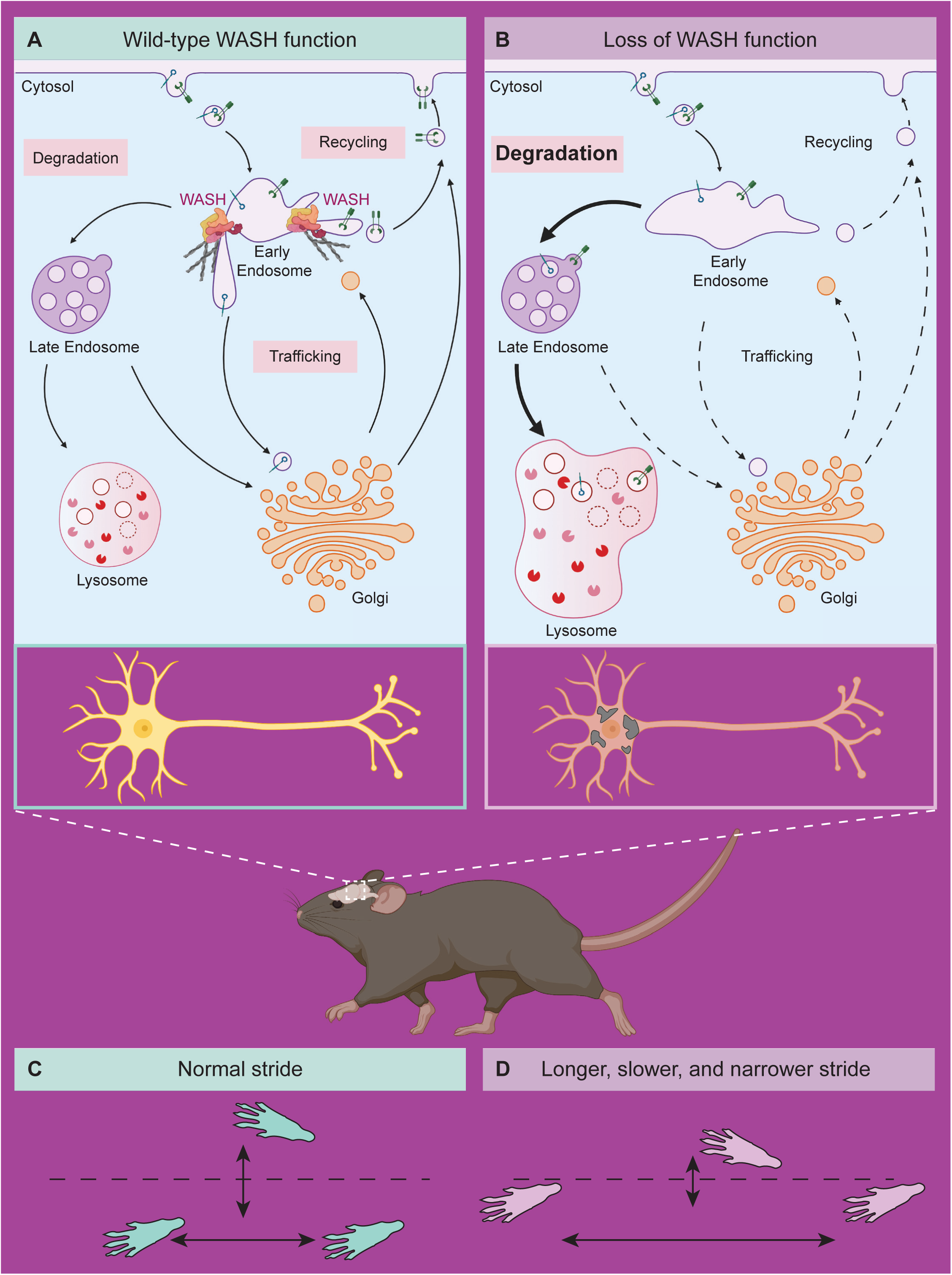
Model of neuronal endo-lysosomal pathology in SWIP^P1019R^ mutant mice. (A) Wild-type WASH function in mouse brain. Under normal conditions, the WASH complex interacts with many endosomal proteins and cytoskeletal regulators, such as the Arp2/3 complex. These interactions enable restructuring of the endosome surface (actin in gray) and allow for cargo segregation and scission of vesicles. Substrates are transported to the late endosome for lysosomal degradation, to the Golgi network for modification, or to the cell surface for recycling. (B) Loss of WASH function leads to increased lysosomal degradation in mouse brain. Destabilization of the WASH complex leads to enlarged endosomes and lysosomes, with increased substrate accumulation at the lysosome. This suggests an increase in flux through the endo-lysosomal pathway, possibly as a result of mis-localized endosomal substrates. (C) Wild-type mice exhibit normal motor function. (D) SWIP^P1019R^ mutant mice display progressive motor dysfunction in association with these subcellular alterations.

Using *in vivo* proximity-based proteomics in wild-type mouse brain, we identify that the WASH complex interacts with the CCC (COMMD9 and CCDC93) and Retriever (VPS35L) cargo selective complexes (Bartuzi et al., 2016; Singla et al., 2019). Interestingly, we did not find significant enrichment of the Retromer sorting complex, a well-known WASH interactor, suggesting that it may play a minor role in neuronal WASH-mediated cargo sorting (Figure 1). These data are supported by our TMT proteomics and covariation network analyses of SWIP^P1019R^ mutant brain, which clustered the WASH, CCC, and Retriever complexes together in M19, but not the Retromer complex, which was found in endosomal module M14 (Figure 2 and Figure 2-figure supplement 3A). Systems-level protein covariation analyses also revealed that disruption of these WASH-CCC-Retriever interactions may have multiple downstream effects on the endosomal machinery, since endosomal modules displayed significant changes in SWIP^P1019R^ brain (including both M19, Figure 2, as well as M14, Figure 2-figure supplement 3A), with corresponding decreases in the abundance of endosomal proteins including Retromer subunits (VPS29 and VPS35), associated sorting nexins (e.g. SNX17 and SNX27), known WASH interactors (e.g. RAB21 and FKBP15), and cargos (e.g. LRP1 and ITGA3) (Figure 2-figure supplements 2 and 3) (Del Olmo et al., 2019; Farfán et al., 2013; Fedoseienko et al., 2018; Halff et al., 2019; Harbour et al., 2012; McNally et al., 2017; Pan et al., 2010; Ye et al., 2020; Zimprich et al., 2011). While previous studies have indicated that Retromer and CCC influence endosomal localization of WASH (Harbour et al., 2012; Phillips-Krawczak et al., 2015; Singla et al., 2019), our findings of altered endosomal networks containing decreased Retromer, Retriever, and CCC protein levels in SWIP^P1019R^ mutant brain point to a possible feedback mechanism wherein WASH impacts the protein abundance and/or stability of these interactors. Future studies defining the hierarchical interplay between the WASH, Retromer, Retriever, and CCC complexes in neurons could provide clarity on how these mechanisms are organized.

In addition to highlighting the neuronal roles of WASH in CCC- and Retriever-mediated endosomal sorting, our proteomics approach also identified protein modules with increased abundance in SWIP^P1019R^ mutant brain. The proteins in these modules fell into two interesting categories: lysosomal enzymes and proteins involved in the endoplasmic reticulum (ER) stress response. Of note, some of the lysosomal enzymes with elevated levels in MUT brain (GRN, M2; IDS, M2; and GNS, M213; Figure 3) are also implicated in lysosomal storage disorders, where they generally have decreased, rather than increased, function or expression (Hopwood et al., 1993; Mok et al., 2003; Schröder et al., 1994; Ward et al., 2017). We speculate that loss of WASH function in our mutant mouse model may lead to increased accumulation of cargo and associated machinery at early endosomes (as seen in Figure 4, enlarged EEA1^+^ puncta), eventually overburdening early endosomal vesicles and triggering transition to late endosomes for subsequent fusion with degradative lysosomes (Figure 8). This would effectively increase delivery of endosomal substrates to the lysosome compared to baseline, resulting in enlarged, overloaded lysosomal structures, and elevated demand for degradative enzymes. For example, since mutant neurons display increased lysosomal module protein abundance (Figure 3), and larger lysosomal structures (Figures 4 and 5), they may require higher quantities of progranulin (GRN, M2; Figure 3) for sufficient lysosomal acidification (Tanaka et al., 2017).

Our findings that SWIP^P1019R^ results in reduced WASH complex stability and function, which may ultimately drive lysosomal dysfunction, are supported by studies in non-mammalian cells. For example, expression of a dominant-negative form of WASH1 in amoebae impairs recycling of lysosomal V-ATPases (Carnell et al., 2011) and loss of WASH in *Drosophila* plasmocytes affects lysosomal acidification (Gomez et al., 2012; Nagel et al., 2017; Zech et al., 2011). Moreover, mouse embryonic fibroblasts lacking WASH1 display abnormal lysosomal morphologies, akin to the structures we observed in cultured SWIP^P1019R^ MUT neurons (Gomez et al., 2012).

In addition to lysosomal dysfunction, endoplasmic reticulum (ER) stress is commonly observed in neurodegenerative states, where accumulation of misfolded proteins disrupts cellular proteostasis (Cai et al., 2016; Hetz and Saxena, 2017; Montibeller and de Belleroche, 2018). This cellular strain triggers the adaptive unfolded protein response (UPR), which attempts to restore cellular homeostasis by increasing the cell’s capacity to retain misfolded proteins within the ER, remedy misfolded substrates, and trigger degradation of persistently misfolded species. Involved in this process are ER chaperones that we identified as increased in SWIP^P1019R^ mutant brain including BiP (HSPA5), calreticulin (CALR), calnexin (CANX), and the protein disulfide isomerase family members (PDIA1, PDIA4, PDIA6) (M83; Figure 2-supplement 3B) (Garcia-Huerta et al., 2016). Many of these proteins were identified in the ER protein module found to be significantly altered in MUT mouse brain (M83), supporting a network-level change in the ER stress response (Figure 2-supplement 3B). One notable exception to this trend was endoplasmin (HSP90B1, M136), which exhibited significantly decreased abundance in SWIP^P1019R^ mutant brain (Table S2). This is surprising given that endoplasmin has been shown to coordinate with BiP in protein folding (Sun et al., 2019), however it may highlight a possible compensatory mechanism. Additionally, prolonged UPR can stimulate autophagic pathways in neurons, where misfolded substrates are delivered to the lysosome for degradation (Cai et al., 2016). These data highlight a relationship between ER and endo-lysosomal disturbances as an exciting avenue for future research.

Strikingly, we observed modules enriched for resident proteins corresponding to all 10 of the major subcellular compartments mapped by Geladaki *et al*. (2019; nucleus, mitochondria, golgi, ER, peroxisome, proteasome, plasma membrane, lysosome, cytoplasm, and ribosome; Supplementary File 1). The greatest dysregulations we observed were in lysosomal, endosomal, ER, and synaptic modules, supporting the hypothesis that SWIP^P1019R^ primarily results in disrupted endo-lysosomal trafficking. While analysis of these dysregulated modules informs the pathobiology of SWIP^P1019R^, our spatial proteomics approach also identified numerous biologically cohesive modules, which remained unaltered (Supplementary File 1). Given that many of these modules contained proteins of unknown function, we anticipate that future analyses of these modules and their protein constituents have great potential to inform our understanding of protein networks and their influence on neuronal cell biology.

It has become clear that preservation of the endo-lysosomal system is critical to neuronal function, as mutations in mediators of this process are implicated in neurological diseases such as Parkinson’s disease, Huntington’s disease, Alzheimer’s disease, Frontotemporal Dementia, Neuronal Ceroid Lipofuscinoses (NCLs), and Hereditary Spastic Paraplegia (Baker et al., 2006; Connor-Robson et al., 2019; Edvardson et al., 2012; Follett et al., 2019; Harold et al., 2009; Mukherjee et al., 2019; Pal et al., 2006; Quadri et al., 2013; Seshadri et al., 2010; Tachibana et al., 2019; Valdmanis et al., 2007). These genetic links to predominantly neurodegenerative conditions have supported the proposition that loss of endo-lysosomal integrity can have compounding effects over time and contribute to progressive disease pathologies. In particular, NCLs are lysosomal storage disorders primarily found in children, with heterogenous presentations and multigenic causations (Mukherjee et al., 2019). The majority of genes implicated in NCLs affect lysosomal enzymatic function or transport of proteins to the lysosome (Mukherjee et al., 2019; Poët et al., 2006; Ramirez-Montealegre and Pearce, 2005; Yoshikawa et al., 2002). Most patients present with marked neurological impairments, such as learning disabilities, motor abnormalities, vision loss, and seizures, and have the unifying feature of lysosomal lipofuscin accumulation upon pathological examination (Mukherjee et al., 2019). While the human SWIP^P1019R^ mutation has not been classified as an NCL (Ropers et al., 2011), findings from our mutant mouse model suggest that loss of WASH complex function leads to phenotypes bearing strong resemblance to NCLs, including lipofuscin accumulation (Figures 4-7). As a result, our mouse model could provide the opportunity to study these pathologies at a mechanistic level, while also enabling preclinical development of treatments for their human counterparts.

Currently there is an urgent need for greater mechanistic investigations of neurodegenerative disorders, particularly in the domain of endo-lysosomal trafficking. Despite the continual increase in identification of human disease-associated genes, our molecular understanding of how their protein equivalents function and contribute to pathogenesis remains limited. Here we employ a systems-level analysis of proteomic datasets to uncover biological perturbations linked to SWIP^P1019R^. We demonstrate the power of combining *in vivo* proteomics and systems network analyses with *in vitro* and *in vivo* functional studies to uncover relationships between genetic mutations and molecular disease pathologies. Applying this platform to study organellar dysfunction in other neurodegenerative and neurodevelopmental disorders may facilitate the identification of convergent disease pathways driving brain disorders.

---

**Figure 2-figure supplement 1.**
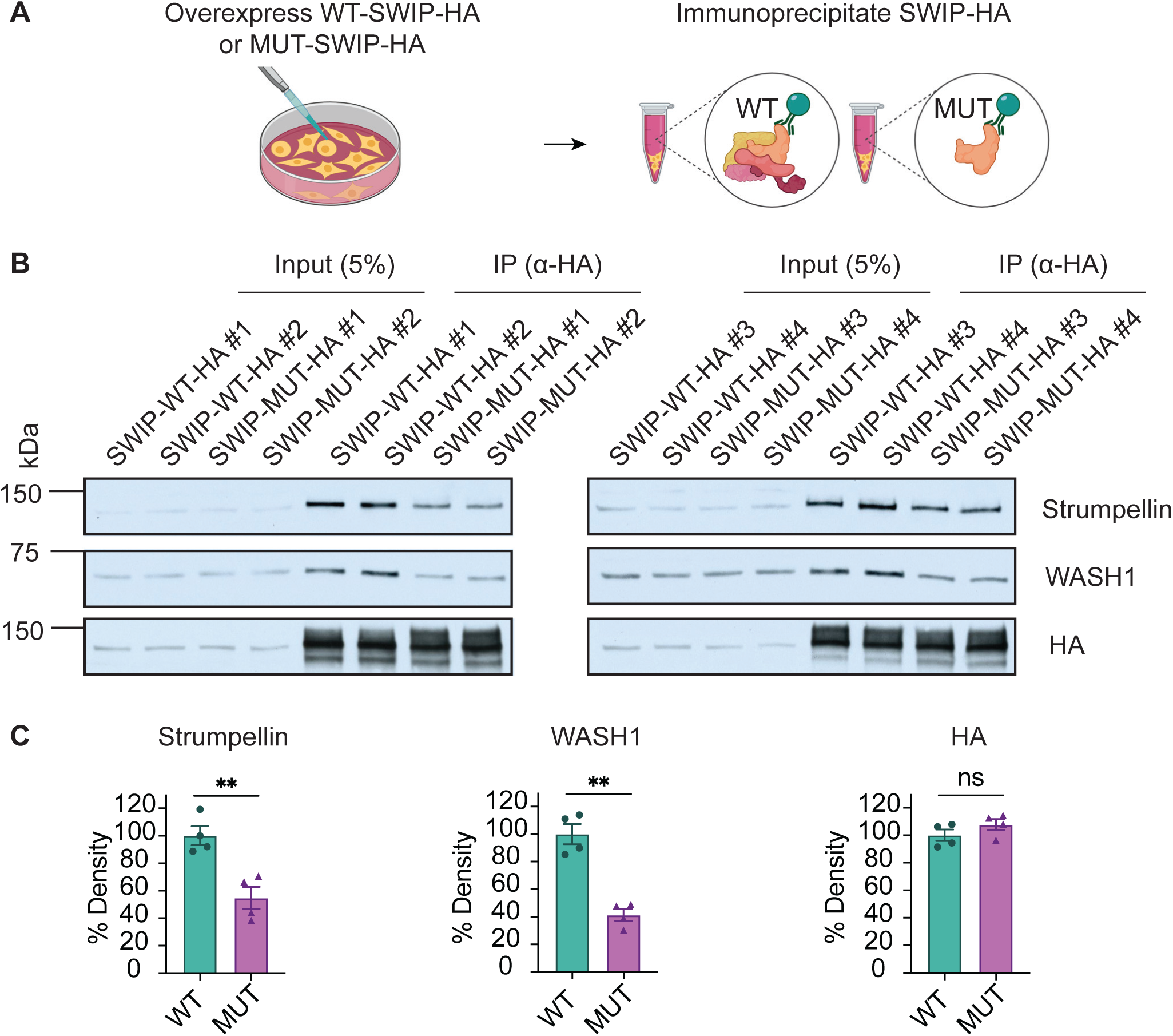
Overexpression of SWIP^P1019R^ decreases WASH complex binding in cultured cells; related to Figure 2. (A) Schematic showing overexpression of WT or MUT SWIP^P1019R^ in HEK293T cells followed by immunoprecipitation. (B) Western blots of input (5%, left) and immunoprecipitated (IP, right) protein. Two samples per condition were run on two separate gels, n=4 biological replicates from separate experiments. (C) Quantification of B normalized to WT. Strumpellin (WT 100.0 ± 6.8%, MUT 54.8 ± 8.0%, t_5.9_=4.290, p=0.0054), WASH1 (WT 100.0 ± 7.3%, MUT 41.4 ± 4.4%, t_4.9_=6.902, p=0.0011), HA (WT 100.0 ± 4.1%, MUT 107.8 ± 4.1%, t_6.0_=1.344, p=0.2275). Data reported as mean ± SEM, error bars are SEM. **p<0.01, two-tailed t-tests.

**Figure 2-figure supplement 2.**
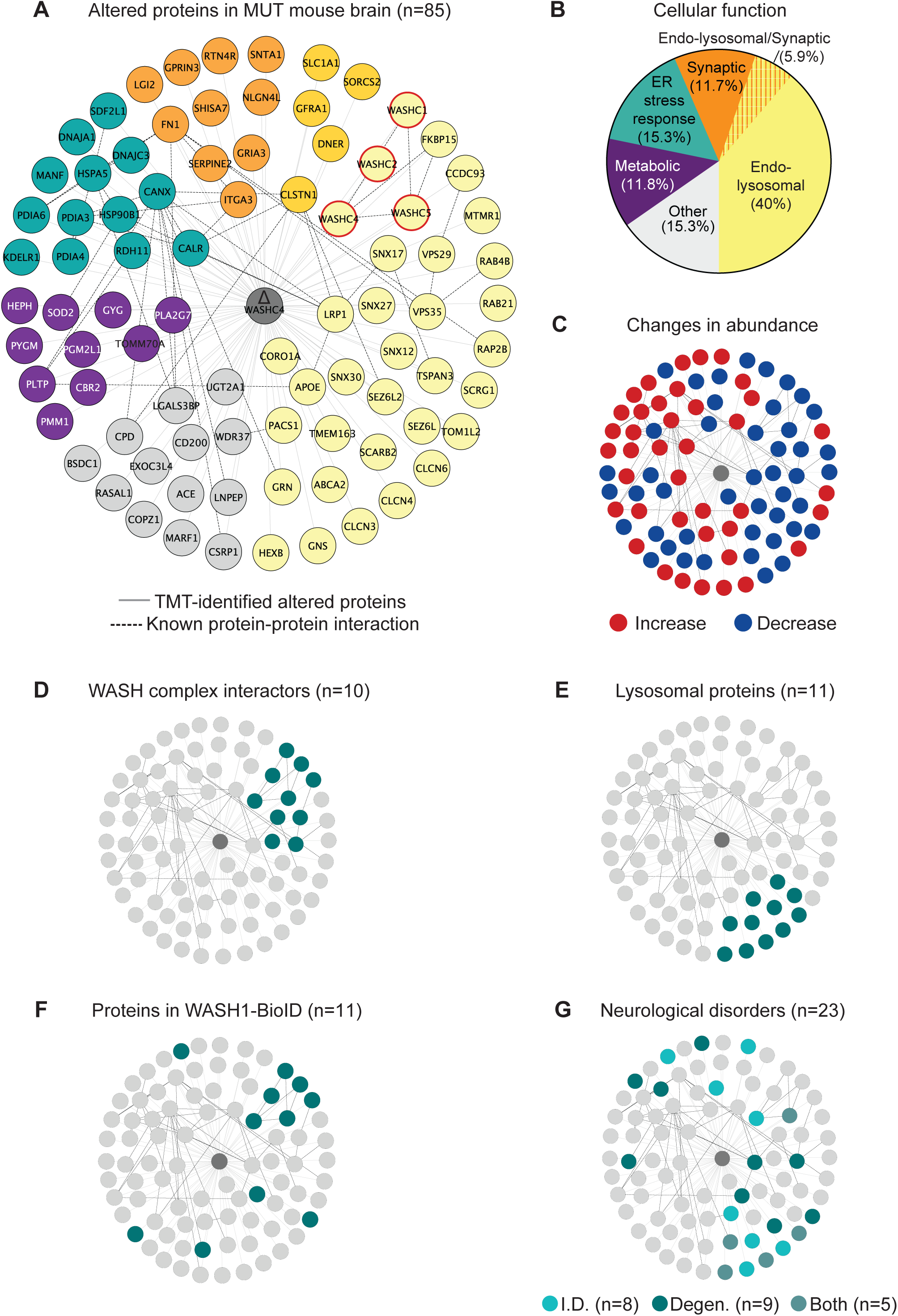
SWIP^P1019R^ MUT brain displays significant alterations in protein abundance compared to WT; related to Figures 2 and 3. (A) Interactome of altered proteins. Nodes reflect protein name, light gray lines delineate proteins identified as different in MUT compared to WT (ΔWASHC4 in center), dark dashed lines indicate known protein-protein interactions from HitPredict database (López et al., 2015). Color reflects cellular function seen in B. Nodes with red borders delineate WASH complex proteins. (B) Cellular function of proteins in A, as reflected in published literature. % reflects the percentage of proteins in a category out of the total 85 altered proteins. (C-G) Clustergrams of: (C) Proteins with increased (red) or decreased (blue) abundance in MUT brains compared to WT. (D) Known protein components of the WASH complex and their previously reported interactors. (E) Proteins with lysosomal function. (F) Proteins identified in the WASH1-BioID2 proteome (Figure 1). (G) Proteins with links to intellectual disability (I.D.) or neurodegeneration (Degen.).

**Figure 2-figure supplement 3.**
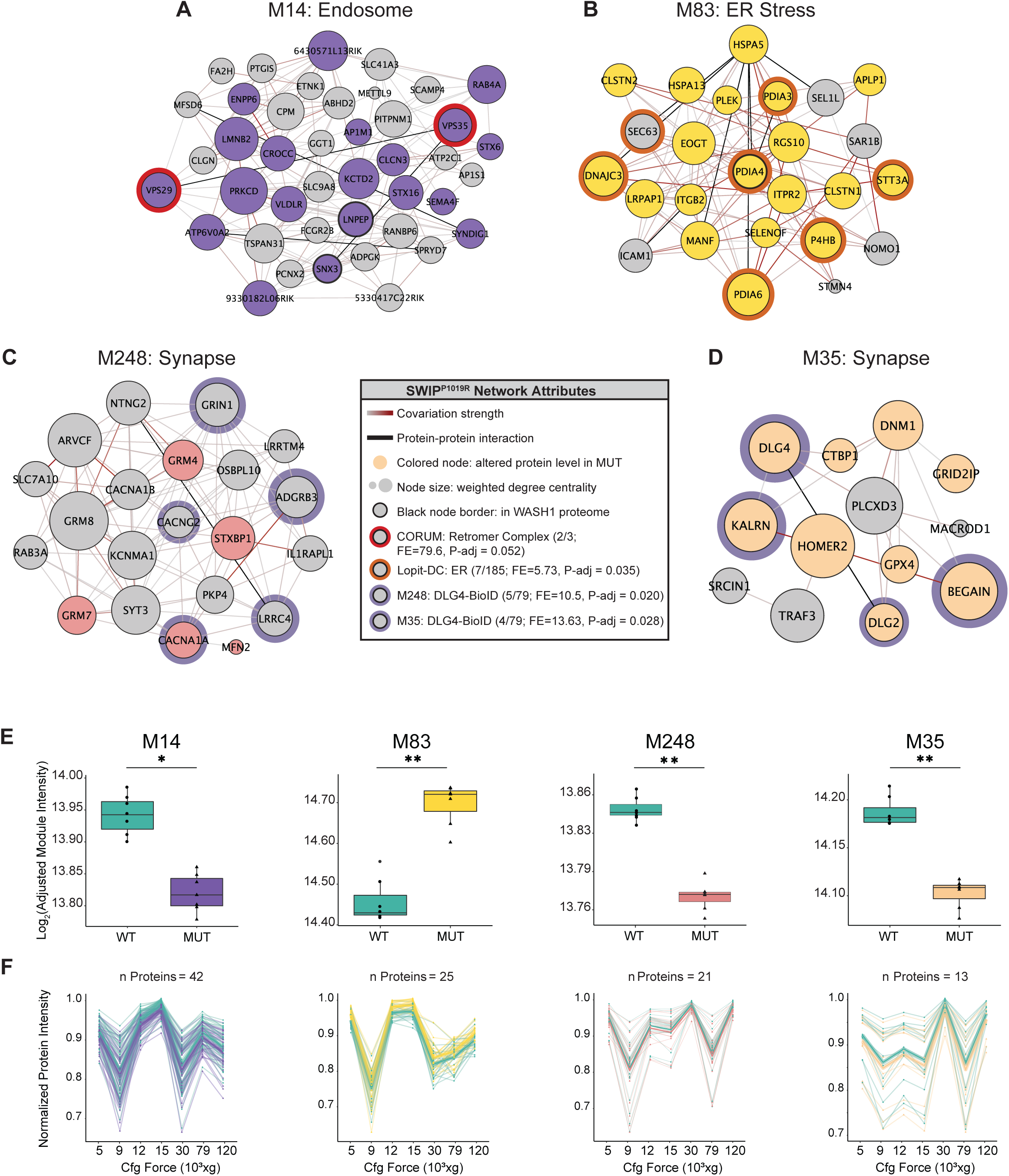
Multiple protein networks display significant alterations in MUT brain compared to WT; related to Figures 2 and 3. (A) Module 14 (M14) containing endosomal proteins. Two of the three retromer sorting complex subunits are highlighted with red borders, with enrichment calculated relative to the CORUM database (Giurgiu et al., 2019). (B) Module 83 (M83) containing endoplasmic reticulum (ER) stress response proteins. Seven of the 185 proteins identified to have ER function by LOPIT-DC spatial proteomics are highlighted with orange borders (Geladaki et al., 2019). (C-D) Modules 248 and 35 (M248, M35, respectively) containing synaptic proteins. Network attributes for graphs in B-E: Node size denotes its weighted degree centrality (∼importance in module), colored node indicates altered abundance in MUT brain relative to WT, black node border denotes proteins identified in the WASH1-BioID proteome (Figure 1), colored node border denotes proteins identified in other datasets, black edges indicate known protein-protein interactions, and grey-red edges denote the relative strength of protein covariation within a module (gray = weak, red = strong). P-adjust enrichment values are calculated relative to the CORUM database (B), Lopit-DC dataset (C), or excitatory postsynapse proteome (DLG4-BioID, D-E) (Geladaki et al., 2019; Giurgiu et al., 2019; Uezu et al., 2016). (E) Summary box plots of module abundance for all seven brain fractions analyzed presented as log2(adjusted module intensities). All modules display significant differences between WT and MUT groups (M14: WT 13.94 ± 0.002 MUT 13.82 ± 0.002, p=0.0207; M83: WT 14.46 ± 0.004, MUT 14.70 ± 0.003, p=0.0088; M248: WT 13.83 ± 0.001, MUT 13.77 ± 0.001, p=0.00134; M35: WT 14.19 ± 0.001, MUT 14.10 ± 0.001, p=0.0030). (F) Normalized protein abundance for all proteins in each module across the seven subcellular fractions analyzed. Plots correspond to modules seen in F, and data colors reflect genotypes in F. Thick lines represent the mean protein intensity per genotype. Number of proteins in each module are indicated at the top of each graph. Data reported as mean ± SEM, error bars are SEM. *p<0.05, **p<0.01, ***p<0.001, GLM (model: 0 ∼ Fraction + Module) and empirical Bayes quasi-likelihood F-test with Bonferroni correction (F).

**Figure 5-figure supplement 1.**
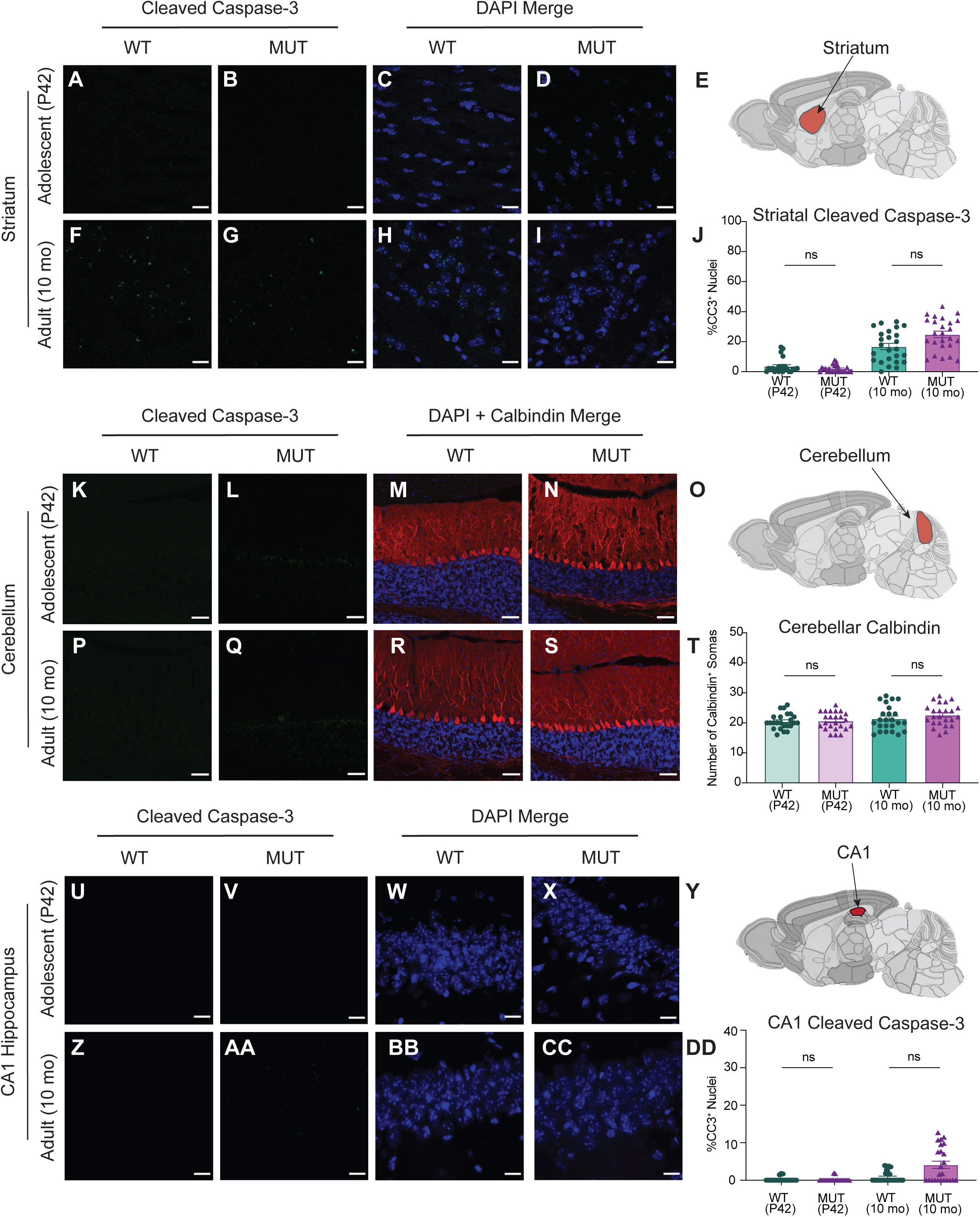
There is no significant difference in striatal, cerebellar, or hippocampal cell death between WT and MUT mice; related to Figure 5. (A) Representative image of adolescent (P42) WT striatum stained with cleaved caspase-3 (CC3, green). (B) Representative image of adolescent (P42) MUT striatum stained with cleaved caspase-3 (CC3, green). (C and D) DAPI co-stained images of A and B, respectively. (E) Anatomical representation of mouse brain with striatum highlighted in red, adapted from the Allen Brain Atlas (Oh et al., 2014). (F) Representative image of adult (10 mo) WT striatum stained with CC3 (green). (G) Representative image of adult (10 mo) MUT striatum stained with CC3 (green). (H and I) DAPI co-stained images of F and G, respectively. Scale bars for A-I are 15 µm. (J) Graph depicting the normalized % of DAPI+ nuclei that are positive for CC3 per image. No difference is seen between genotypes at either age (P42 WT 3.70 ± 0.99%, P42 MUT 1.95 ± 0.49%, 10mo WT 16.77 ± 2.09%, 10mo MUT 24.86 ± 2.17%, H=61.87, p<0.0001). (K) Representative image of adolescent (P42) WT cerebellum stained with cleaved caspase-3 (CC3, green). (L) Representative image of adolescent (P42) MUT cerebellum stained with cleaved caspase-3 (CC3, green). (M and N) DAPI and Calbindin co-stained images of K and L, respectively. (O) Anatomical representation of mouse cerebellum, adapted from the Allen Brain Atlas (Oh et al., 2014). Red region highlights area used for imaging. (P) Representative image of adult (10 mo) WT cerebellum stained with CC3 (green). (Q) Representative image of adult (10 mo) MUT cerebellum stained with CC3 (green). No significant CC3 staining is observed at either age. (R and S) DAPI and Calbindin co-stained images of P and Q, respectively. Scale bars for K-S are 50 µm. (T) Graph depicting the number of Calbindin+ somas per image, a marker for Purkinje cells. No difference is seen between genotypes at either age (P42 WT 20.50 ± 0.53, P42 MUT 20.67 ± 0.59, 10mo WT 21.42 ± 0.85, 10mo MUT 22.63 ± 0.74, H=4.891, p=0.1799). (U) Representative image of adolescent (P42) WT cerebellum stained with cleaved caspase-3 (CC3, green). (V) Representative image of adolescent (P42) MUT cerebellum stained with cleaved caspase-3 (CC3, green). (W and X) DAPI and Calbindin co-stained images of U and V, respectively. (Y) Anatomical representation of mouse hippocampus CA1, adapted from the Allen Brain Atlas (Oh et al., 2014). Red region highlights area used for imaging. (Z) Representative image of adolescent (P42) WT cerebellum stained with cleaved caspase-3 (CC3, green). (AA) Representative image of adolescent (P42) MUT cerebellum stained with cleaved caspase-3 (CC3, green). (BB and CC) DAPI and Calbindin co-stained images of U and V, respectively. (DD) Graph of the normalized % of DAPI+ nuclei that are positive for CC3 per image. Very little CC3 staining is seen at either age, regardless of genotype (P42 WT 0.21 ± 0.11%, P42 MUT 0.15 ± 0.11%, 10mo WT 0.81 ± 0.28%, 10mo MUT 4.13 ± 0.96%, H=20.27, p=0.0001). Data obtained from four animals per condition, and reported as mean ± standard error of the mean (SEM), with error bars as SEM. Kruskal-Wallis test (J,T, and DD).

**Figure 6-figure supplement 1.**
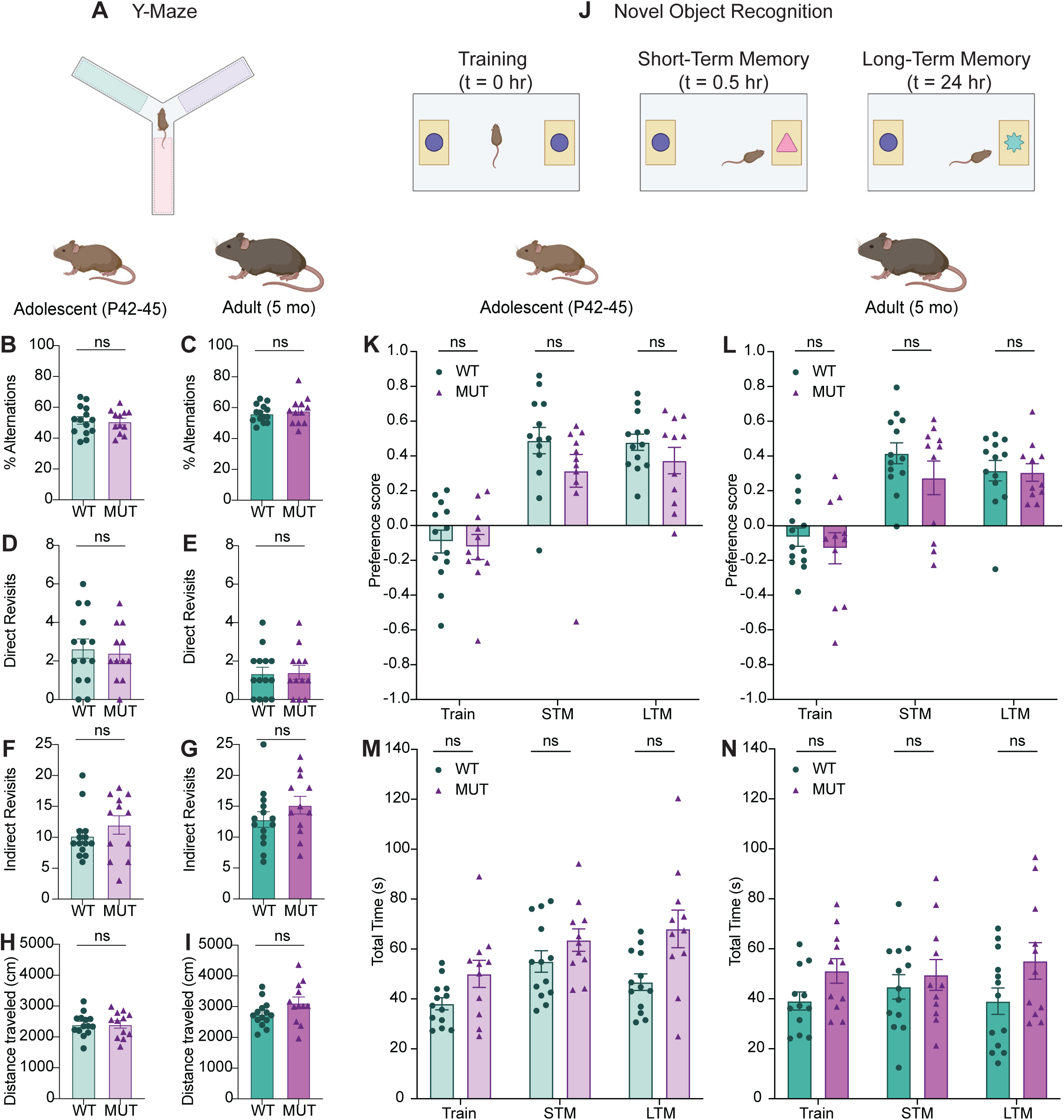
SWIP^P1019R^ mutant mice do not display deficits in spatial working memory or novel object recognition; related to Figure 6. (A) Y-maze paradigm. Mice were placed in the center of the maze and allowed to explore all three arms freely for five minutes. Each arm had distinct visual cues. (B) Graph depicting the percent alternations achieved for each mouse at adolescence (WT 51.48 ± 2.47%, MUT 50.81 ± 2.19%, t_24_=0.2036, p=0.8404). (C) Graph depicting the percent alternations achieved for each mouse at adulthood (WT 56.12 ±1.53%, MUT 57.93 ± 2.56%, t_18_=0.6074, p=0.5511). (D) Graphs of the number of direct revisits mice made to the arm they just explored reveal no difference between genotypes at adolescence (WT 2.64 ± 0.50, MUT 2.42 ± 0.42, t_24_=0.3482, p=0.7308). (E) Graphs of the number of direct revisits mice made at adulthood (WT 1.36 ± 0.33, MUT 1.42 ± 0.36, t_24_=0.1231, p=0.9031). (F) Similar to D-E, there were no differences in the number of indirect revisits (ex: arm A◊ arm B◊arm A) between genotypes at adolescence (WT 10.21 ± 1.03, MUT 12.00 ± 1.48, U=69, p=0.4515) (G) Indirect revisits at adulthood (WT 12.86 ± 1.26, MUT 15.17 ± 1.43, t_23_=1.211, p=0.2380). (H) There were no significant differences in total distance travelled at adolescence, suggesting that motor function did not affect Y-maze performance (WT 2401 ± 98.9 cm, MUT 2406 ± 121.0 cm, t_22_=0.03281, p=0.9741). (I) No difference in distanced travelled at adulthood (WT 2761 ± 111.6 cm, MUT 3124 ± 191.1 cm, t_18_=1.638, p=0.1189). (J) Novel object task. Mice first performed a 5-minute trial in which they were placed in an arena with two identical objects and allowed to explore freely, while their behavior was tracked with video software. Mice were returned to their home cage, and then re-introduced to the arena a half an hour later, where one of the objects had been replaced with a novel object, and their behavior was again tracked. Twenty-four hours later the same test was performed, but the novel object was replaced with another new object. (K) Graph depicting adolescent animals’ preference for the novel object during the three phases of the task, training (Train), short-term memory (STM), and long-term memory (LTM). No significant difference in object preference is seen between genotypes for any phase (Train WT −0.092 ± 0.065, Train MUT −0.123 ± 0.072, STM WT 0.488 ± 0.076, STM MUT 0.315 ± 0.094, LTM WT 0.479 ± 0.046, LTM MUT 0.373 ± 0.076, F_1,22_=1.840, p=0.1887). (L) Graph depicting adult animals’ preference for the novel object. (Train WT −0.066 ± 0.053, Train MUT −0.130 ± 0.089, STM WT 0.416 ± 0.060, STM MUT 0.274 ± 0.096, LTM WT 0.316 ± 0.059, LTM MUT 0.306 ± 0.050, F_1,22_=0.9735, p=0.3345). (M) Graph depicting the total amount of time (in seconds) adolescent animals spent exploring both objects in each phase of the task. No significant difference in exploration time was observed across genotypes, suggesting that genotype does not hinder object exploration (Train P44 WT 38.10 ± 2.45 s, Train P44 MUT 50.04 ± 5.44 s, STM P44 WT 55.02 ± 4.31 s, STM P44 MUT 63.60 ± 4.50 s, LTM P45 WT 46.75 ± 3.34 s, LTM P45 MUT 68.08 ± 7.54 s, F_1,22_=7.373, p=0.0126). (N) Graph depicting the total time adult animals spent exploring both objects in each the task (Train WT 39.10 ± 3.62 s, Train MUT 51.15 ± 4.91 s, STM WT 44.75 ± 4.87 s, STM MUT 49.56 ± 6.16 s, LTM WT 39.02 ± 5.29 s, LTM MUT 55.15 ± 7.34 s, F_1,22_=2.936, p=0.1007). For all Y-maze measures, WT n=14, MUT n=12. For all novel object measures, WT n=13, MUT n=11. Data reported as mean ± SEM, error bars are SEM. Two-tailed t-tests or Mann-Whitney U tests (B-I), two-way ANOVAs (K-N).

**Figure 6-figure supplement 2.**
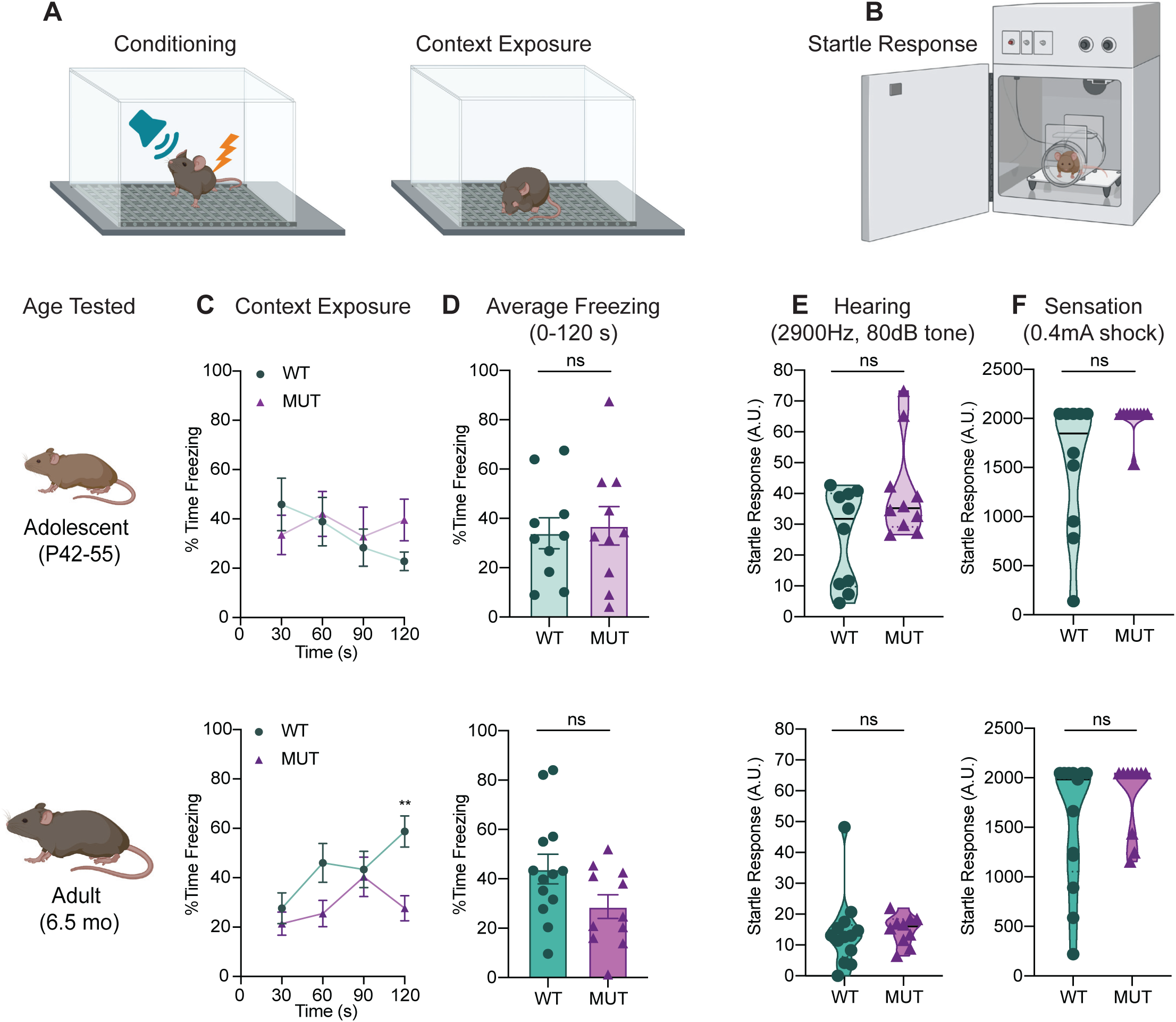
SWIP^P1019R^ mutant mice do not have significant deficits in contextual fear memory recall, auditory perception, or tactile sensation; related to Figure 6. (A) Experimental fear conditioning scheme. After acclimation, mice received a mild aversive 0.4mA footshock paired with a 2900Hz tone in a conditioning chamber. 24 hours later, the mice were placed back in the same chamber to assess freezing behavior (without footshock or tone). (B) Experimental startle response setup used to assess hearing and somatosensation. Mice were placed in a plexiglass tube atop a load cell that measured startle movements in response to stimuli. (C) Line graphs of WT and MUT freezing response during the contextual memory recall task. Data represented as average freezing per genotype in 30 second time bins. The task was performed with two different cohorts for the different ages, P42 (top) and 6.5mo (bottom). Top: no significant difference in freezing at P42 (Two-way repeated measure ANOVA, Genotype effect, F_1,18_=0.088, p=0.7698. Sidak’s post-hoc analysis, 30 s p=0.8388, 60 s p=0.9990, 90 s p=0.9964, 120 s p=0.3281), Bottom: no significant difference in freezing at 6.5mo (Two-way repeated measure ANOVA, Genotype effect, F_1,22_= 3.723, p=0.0667. Sidak’s post-hoc analysis, 30 s p=0.8977, 60 s p=0.1636, 90 s p=0.9979, 120 s p=0.0037). (D) Graphs showing the average total freezing time per animal during context exposure. Top: no significant difference is seen between WT and MUT mice at P42 (WT 34.01 ± 6.32%, MUT 36.99 ± 7.81%, t_17_=2.985, p=0.7699). Bottom: no significant different is seen between genotypes at 6.5mo (WT 43.94 ± 6.00%, MUT 28.73 ± 4.80%, t_21.6_=1.980, p=0.0606). (E) Graphs of individual animals’ startle response to a 2900Hz tone played at 80dB. MUT mice were not significantly more reactive to the tone than WT at P50 (WT 25.96 ± 4.95, MUT 40.68 ± 5.05, U=35, p=0.2799), or at 6.5mo (WT 14.07 ± 3.27, MUT 14.85 ± 1.49, U=47, p=0.2768). (F) Graphs of individual animals’ startle response to a 0.4mA footshock. No significant difference observed between genotypes at either age (P55 WT 1527 ± 215.7, P55 MUT 1996 ± 51.0, U=28.50, p=0.0542; 6.5mo WT 1545 ± 179.5, 6.5mo MUT 1817 ± 119.1, U=47, p=0.2360). Startle response reported in arbitrary units (A.U.). For all adolescent measures: WT n=10, MUT n=10. For adult freezing measures: WT n=13, MUT n=11. For adult startle responses: WT n=13, MUT n=10. Data reported as mean ± SEM, error bars are SEM.*p<0.05, **p<0.01, two-way repeated measure ANOVAs (C), two-tailed t-tests (D), and Mann-Whitney U tests (E-F).

**Figure 7-figure supplement 1.**
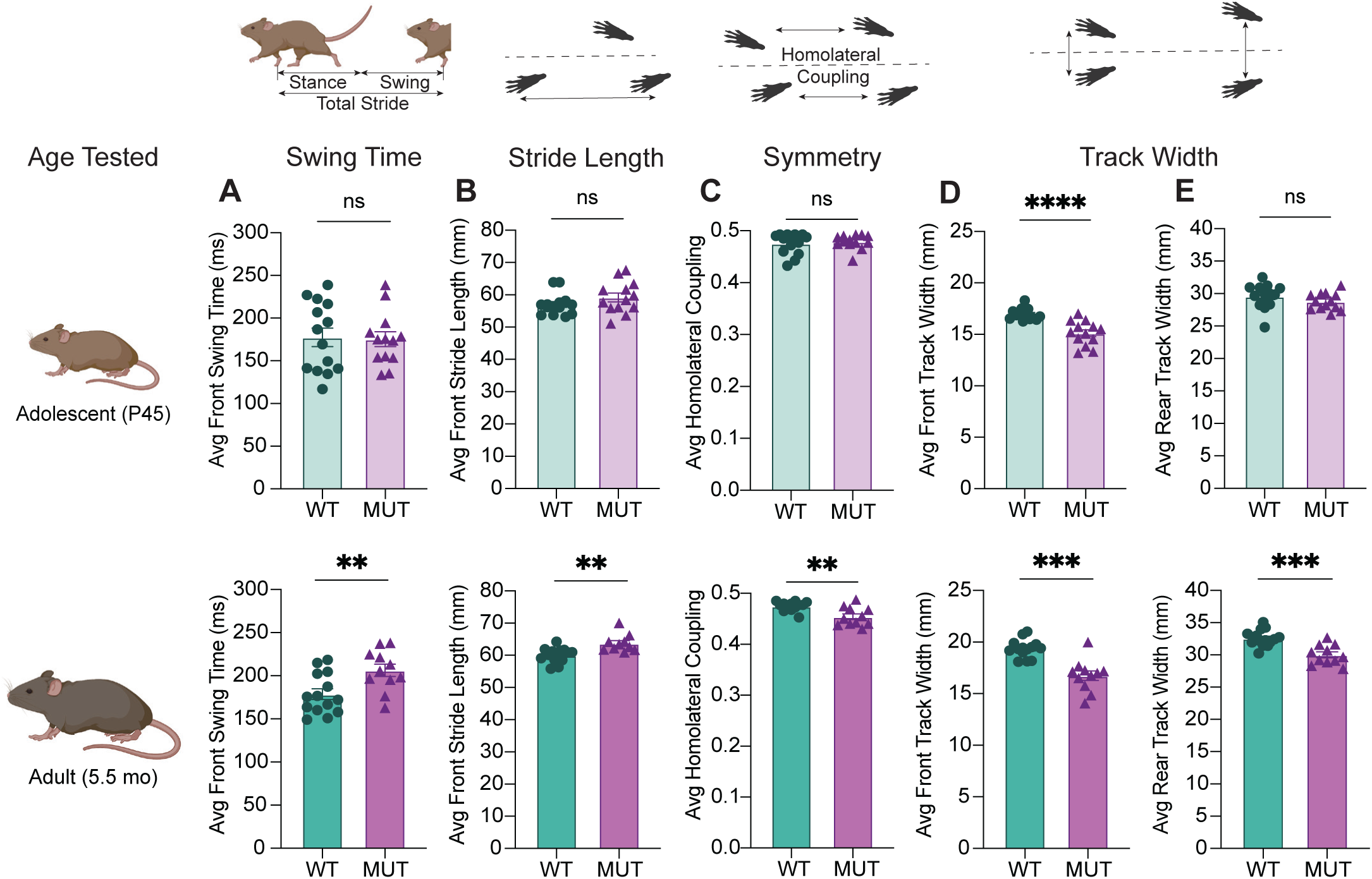
Progressive gait changes in SWIP^P1019R^ mutant mice are not restricted to rear limbs; related to Figure 7. (A) Graph of average swing time per stride for front limbs. At P45 (top), there is no significant difference in front swing time (WT 177.5 ± 10.9 ms, MUT 175.3 ± 8.7 ms, t_24_=0.1569, p=0.8766). At 5.5 mo (bottom), MUT mice take significantly longer to swing their forelimbs (WT 178.6 ± 6.2 ms, MUT 206.3 ± 7.2 ms, t_21.4_=2.927, p=0.0079). (B) Graph of average stride length for front limbs. At P45 (top), there is no difference in WT and MUT stride length (WT 57.0 ± 0.9 mm, MUT 59.2 ± 1.4 mm, U=68, p=0.2800). At 5.5mo (bottom), MUT mice take significantly longer strides with their forelimbs (WT 60.0 ± 0.6 mm, MUT 63.7 ± 0.9 mm, t_17_=3.545, p=0.0024). (C) Graph of average homolateral coupling, the fraction of a reference foot’s stride when its ipsilateral foot starts its stride. At P45, there is no significant difference in homolateral coupling (WT 0.48 ± 0.005, MUT 0.48 ± 0.004, U=90, p=0.9713), but at 5.5mo, MUT mice display decreased homolateral coupling (WT 0.48 ± 0.002, MUT 0.45 ± 0.005, t_13.5_=3.469, p=0.0039). (D) Graph of average track width between front limbs. At P45 (top), there is a significantly narrower front track width in MUT compared to WT (WT 16.99 ± 0.15 mm, MUT 15.12 ± 0.33 mm, t_17_=5.192, p<0.0001). This difference persists into adulthood at 5.5mo (WT 19.36 ± 0.23 mm, MUT 16.74 ± 0.46 mm, t_15_=5.055, p=0.0001). (E) Graph of average track width between rear limbs. At P45 (top), there is no difference in WT and MUT rear track widths (WT 29.58 ± 0.51 mm, MUT 28.77 ± 0.36 mm, t_23_=1.292, p=0.2091). At 5.5mo (bottom), mutants display significantly narrower rear track widths (WT 32.59 ± 0.34 mm, MUT 30.01 ± 0.46 mm, t_19.4_=4.502, p=0.0002). For P45 measures: WT n=14, MUT n=13; for 5.5mo measures: WT n=14, MUT n=11. Data reported as mean ± SEM, error bars are SEM. **p<0.01, ***p<0.001, ****p<0.0001, two-tailed t-tests or Mann-Whitney U tests.

## ACKNOWLEDGEMENTS

We are very grateful for the human *WASHC4^c.3056C>G^* patients’ contributions to this study. We also thank the Duke Transgenic Core Facility for their work in generating the SWIP^P1019R^ mutant mice, as well as the Duke Behavioral Core Facility, the Duke Proteomics and Metabolomics Shared Resource, the Duke Electron Microscopy Core Facility, and the Duke Light Microscopy Core Facility for their support in completing these experiments. We also greatly appreciate everyone who provided advice on this project and reviewed this manuscript, including Drs. Alicia Purkey, Shataakshi Dube, Anne West, Cagla Eroglu, and Nicole Calakos. We generated experimental schematics using a personal academic <u>BioRender.com</u> license. This work was supported by a Translating Duke Health Neuroscience Initiative Grant to SHS, NIH grants (MH111684 and DA047258) to SHS, an NIH grant (MH117429) and NARSAD young investigator grant (25163) to IHK, NIH F30 fellowship funding (MH117851) and MSTP training grant support (GM007171) for JLC, and NIH F31 fellowship funding (5F31NS113738-03) to TWAB.

## AUTHOR CONTRIBUTIONS

Conceptualization: JLC, TWAB, IHK, and SHS; Methodology: JLC, TWAB, IHK, and SHS; Investigation: JLC, TWAB, IHK, GW, TH, RV, ES, and AR; Resources: ES, RV, and SHS; Writing—Original Draft: JLC and SHS; Writing—Editing: JLC, TWAB, IHK, and SHS; Visualization: JLC; Funding Acquisition: JLC, TWAB, IHK, and SHS. All authors discussed the results and commented on the manuscript.

## RESOURCE AVAILABILITY

### Lead Contact

Further information and requests for resources and reagents should be directed to and will be fulfilled by the Lead Contact, Scott Soderling.

### Materials Availability

Plasmids generated by this study are available upon request from corresponding author Scott H. Soderling.

### Data and Code Availability

The data and source code used in this study are available online at https://github.com/twesleyb/SwipProteomics.

## MATERIALS AND METHODS

### Animals

We generated *Washc4* mutant (SWIP^P1019R^) mice in collaboration with the Duke Transgenic Core Facility to mimic the *de novo* human variant at amino acid 1019 of human *WASHc4.* A CRISPR-induced CCT>CGT point mutation was introduced into exon 29 of *Washc4*. 50ng/µl of the sgRNA (5’-ttgagaatactcacaagaggagg-3’), 100ng/µl Cas9 mRNA, and 100ng/µl of a repair oligonucleotide containing the C>G mutation were injected into the cytoplasm of B6SJLF1/J mouse embryos (Jax #100012) (See Table S4 for the sequence of the repair oligonucleotide). Mice with germline transmission were then backcrossed into a C57BL/ 6J background (Jax #000664). At least 5 backcrosses were obtained before animals were used for behavior. We bred heterozygous SWIP^P1019R^ mice together to obtain age-matched mutant and wild-type genotypes for cell culture and behavioral experiments. Genetic sequencing was used to screen for germline transmission of the C>G point mutation (*FOR:* 5’-tgcttgtagatgtttttcct-3’, *REV*: 5’-gttaacatgatcctatggcg-3’). All mice were housed in the Duke University′s Division of Laboratory Animal Resources or Behavioral Core facilities at 2-5 animals/cage on a 14:10h light:dark cycle. All experiments were conducted with a protocol approved by the Duke University Institutional Animal Care and Use Committee in accordance with NIH guidelines.

### Human Subjects

We retrospectively analyzed clinical findings from seven children with homozygous *WASHC4*^c.3056C>G^ mutations (obtained by Dr. Rajab in 2010 at the Royal Hospital, Muscat, Oman). The original report of these human subjects and parental consent for data use can be found in (Ropers et al., 2011).

### Cell Lines

HEK293T cells (ATCC #CRL-11268) were purchased from the Duke Cell Culture facility. and were tested for mycoplasma contamination. HEK239T cells were used for co-immunoprecipitation experiments and preparation of AAV viruses.

### Primary Neuronal Culture

Primary neuronal cultures were prepared from P0 mouse cortex. P0 mouse pups were rapidly decapitated and cortices were dissected and kept individually in 5ml Hibernate A (Thermo #A1247501) supplemented with 2% B27(Thermo #17504044) at 4°C overnight to allow for individual animal genotyping before plating. Neurons were then treated with Papain (Worthington #LS003120) and DNAse (VWR #V0335)-supplemented Hibernate A for 18min at 37°C and washed twice with plating medium (plating medium: Neurobasal A (Thermo #10888022) supplemented with 10% horse serum, 2% B-27, and 1% GlutaMAX (Thermo #35050061)), and triturated before plating at 250,000 cells/well on poly-L-lysine-treated coverslips (Sigma #P2636) in 24-well plates. Plating medium was replaced with growth medium (Neurobasal A, 2% B-27, 1% GlutaMAX) 2 hours later. Cell media was supplemented and treated with AraC at DIV5 (5uM final concentration/well). Half-media changes were then performed every 4 days.

### Plasmid DNA Constructs

For immunoprecipitation experiments, a pmCAG-SWIP-WT-HA construct was generated by PCR amplification of the human *WASHC4* sequence, which was then inserted between NheI and SalI restriction sites of a pmCAG-HA backbone generated in our lab. Site-directed mutagenesis (Agilent #200517) was used to introduce a C>G point mutation into this pmCAG-SWIP-WT-HA construct for generation of a pmCAG-SWIP-MUT-HA construct *(FOR:* 5’-ctacaaagttgagggtcagacggggaacaattatatagaaa-3’, *REV:* 5’-tttctatataattgttccccgtctgaccctcaactttgtag-3’). For iBioID experiments, an AAV construct expressing hSyn1-WASH1-BioID2-HA was generated by cloning a *Washc1* insert between SalI and HindIII sites of a pAAV-hSyn1-Actin Chromobody-Linker-BioID2-pA construct (replacing Actin Chromobody) generated in our lab. This backbone included a 25nm GS linker-BioID2-HA fragment from Addgene #80899, generated by Kim *et al*. (Kim et al., 2016). An hSyn1-solubleBioID2-HA construct was created similarly, by removing Actin Chromobody from the above construct. Oligonucleotide sequences are reported in Table S4. Sequences of the plasmid DNA constructs are available online (see Key Resources Table).

### AAV Viral Preparation

AAV preparations were performed as described previously(Uezu et al., 2016). The day before transfection, HEK293T cells were plated at a density of 1.5×10^7^ cells per 15cm^2^ plate in DMEM media with 10% fetal bovine serum and 1% Pen/Strep (Thermo #11965-092, Sigma #F4135, Thermo #15140-122). Six HEK293T 15cm^2^ plates were used per viral preparation. The next day, 30µg of pAd-DeltaF6 helper plasmid, 15µg of AAV2/9 plasmid, and 15µg of an AAV plasmid carrying the transgene of interest were mixed in OptiMEM with PEI-MAX (final concentration 80µg/ml, Polysciences #24765). 2ml of this solution were then added dropwise to each of the 6 HEK293T 15cm^2^ plates. Eight hours later, the media was replaced with 20ml DMEM+10%FBS. 72 hours post-transfection, cells were scraped and collected in the media, pooled, and centrifuged at 1,500rpm for 5min at RT. The final pellet from the 6 cell plates was resuspended in 5ml of cell lysis buffer (15 mM NaCl, 5 mM Tris-HCl, pH 8.5), and freeze-thawed three times using an ethanol/dry ice bath. The lysate was then treated with 50U/ml of Benzonase (Novagen #70664), for 30min in a 37°C water bath, vortexed, and then centrifuged at 4,500rpm for 30min at 4°C. The resulting supernatant containing AAV particles was added to the top of an iodixanol gradient (15%, 25%, 40%, 60% top to bottom) in an Optiseal tube (Beckman Coulter #361625). The gradient was then centrifuged using a Beckman Ti-70 rotor in a Beckman XL-90 ultracentrifuge at 67,000rpm for 70min, 18°C. The purified viral solution was extracted from the 40%/60% iodixanol interface using a syringe, and placed into an Amicon 100kDa filter unit (#UFC910024). The viral solution was washed in this filter 3 times with 1X ice-cold PBS by adding 5ml of PBS and centrifuging at 4,900rpm for 45min at 4°C to obtain a final volume of approximately 200µl of concentrated virus that was aliquoted into 5-10µl aliquots and stored at −80°C until use.

### Immunocytochemistry

Primary antibodies: Rabbit anti-EEA1 (Cell Signaling Technology #C45B10, 1:500), Rat anti-CathepsinD (Novus #204712, 1:250), Guinea Pig anti-MAP2 (Synaptic Systems #188004, 1:500) Secondary antibodies: Goat anti-Rabbit Alexa Fluor 568 (Invitrogen #A11036, 1:1000), Goat anti-Guinea Pig Alexa Fluor 488 (Invitrogen #A11073, 1:1000), Goat anti-Rat Alexa Fluor 488 (Invitrogen #A11006, 1:1000), Goat anti-Guinea Pig Alexa Fluor 555 (Invitrogen #A21435, 1:1000) At DIV15, neurons were fixed for 15 minutes using ice-cold 4%PFA/4% sucrose in 1X PBS, pH 7.4 (for EEA1 staining), or 30 minutes with 50% Bouin’s solution/4% sucrose (for CathepsinD staining, Sigma #HT10132), pH 7.4(Cheng et al., 2018). Fixed neurons were washed with 1X PBS, then permeabilized with 0.25% TritonX-100 in PBS for 8 minutes at room temperature (RT), and blocked with 5%normal goat serum/0.2%Triton-X100 in PBS (blocking buffer) for 1 hour at RT with gentle rocking. For EEA1/MAP2 staining, samples were incubated with primary antibodies diluted in blocking buffer at RT for 1 hour. For CathepsinD/MAP2 staining, samples were incubated with primary antibodies diluted in blocking buffer overnight at 4°C. For both conditions, samples were washed three times with 1X PBS, and incubated for 30min at RT with secondary antibodies, protected from light. After secondary antibody staining, coverslips were washed three times with 1X PBS, and mounted with FluoroSave mounting solution (Sigma #345789). See antibody section for primary and secondary antibody concentrations.

### Immunohistochemistry

Primary antibodies: Rabbit anti-Cleaved Caspase-3 (Cell Signaling Technology #9661, 1:2000), Mouse anti-Calbindin (Sigma #C9848, 1:2000), Rat anti-HA 3F10 (Sigma #12158167001, 1:500) Secondary antibodies: Donkey anti-Rabbit Alexa Fluor 488 (Invitrogen #A21206, 1:2000), Goat anti-Mouse Alexa Fluor 594 (Invitrogen #A11032, 1:2000), Goat anti-Rat Alexa Fluor 488 (Invitrogen #A11006, 1:5000), Streptavidin Alexa Fluor 594 conjugate (Invitrogen #S32356, 1:5000), 4′,6-diamidino-2-phenylindole (DAPI, Sigma #D9542, 1:1000 for 10min at RT) Mice were deeply anesthetized with isoflurane and then transcardially perfused with ice-cold heparinized PBS (25U/ml) by gravity flow. After clearing of liver and lungs (∼2min), perfusate was switched to ice-cold 4% PFA in 1X PBS (pH 7.4) for 15 minutes. Brains were dissected, post-fixed in 4%PFA overnight at 4°C, and then cryoprotected in 30% sucrose/1X PBS for 48hr at 4°C. Fixed brains were then mounted in OTC (Sakura TissueTek #4583) and stored at −20°C until cryosectioning. Every third sagittal section (30 µm thickness) was collected from the motor cortex and striatal regions. Free-floating sections were then permeabilized with 1%TritonX-100 in 1X PBS at RT for 2 hr, and blocked in 1X blocking solution (Abcam #126587) diluted in 0.2%TritonX-100 in 1X PBS for 1hr at RT. Sections were then incubated in primary antibodies diluted in the 1X blocking solution for two overnights at 4°C. After three washes with 0.2%TritonX-100 in 1X PBS, the sections were then incubated in secondary antibodies diluted in 1X blocking buffer for one overnight at 4°C. Sections were then washed four times with 0.2%TritonX-100 in 1X PBS at RT, and mounted onto coverslips with FluoroSave mounting solution (Sigma #345789).

### Western Blotting

Primary antibodies: Rabbit anti-Strumpellin (Santa Cruz #sc-87442, 1:500), Rabbit anti-WASH1 c-terminal (Sigma #SAB4200373, 1:500), Mouse anti-Beta Tubulin III (Sigma #T8660, 1:10,000), Mouse anti-HA (BioLegend #MMS-101P, 1:5000) Secondary antibodies: Donkey anti-Rabbit-HRP (GE Life Sciences #NA934, 1:5,000), Goat anti-mouse-HRP (GE Life Sciences #NA931, 1:5000) Ten micrograms of each sample were electrophoresed through a 12-well, 4-20% SDS-PAGE gel (Bio-Rad #4561096) at 100V for 1hr at RT, transferred onto a nitrocellulose membrane (GE Life Sciences #GE10600002) at 100V for 70min at RT on ice, and blocked with 5% nonfat dry milk in TRIS-buffered saline containing 0.05% Tween-20 (TBST, pH 7.4). Gels were saved for Coomassie staining at RT for 30 min. Membranes were probed with one primary antibody at a time for 24hr at 4°C, then washed four times with TBST at RT before incubating with the corresponding species-specific secondary antibody at RT for 1hr. Membranes were washed with TBST, and then enhanced chemiluminescence (ECL) substrate was added (Thermo Fischer #32109). Membranes were exposed to autoradiography films and scanned with an Epson 1670 at 600dpi. We probed each membrane with one antibody at a time, then stripped the membrane with stripping buffer (Thermo Fischer #21059) for 10min at RT, and then blocked for 1hr at RT before probing with the next antibody. Order of probes: Strumpellin, then β-Tubulin, then WASH1. We determined the optical density of the bands using Image J software (NIH).

Data obtained from three independent experiments were plotted and statistically analyzed using GraphPad Prism (version 8) software.

### Immunoprecipitation

HEK293T cells were transfected with pmCAG-SWIP-WT-HA or pmCAG-SWIP-MUT-HA constructs for three days, as previously described(Mason et al., 2011). Cells were lysed with lysis buffer (25mM HEPES, 150mM NaCl, 1mM EDTA, 1% NonidetP-40, pH 7.4) containing protease inhibitors (5mM NaF, 1mM orthovanadate, 1mM AEBSF, and 2 μg/mL leupeptin/pepstatin) and centrifuged at 1,700g for 5 min. Collected supernatant was incubated with 30µl of pre-washed anti-HA agarose beads (Sigma #A2095) on a sample rotator (15 rpm) for 2 hrs at 4°C. Beads were then washed 3 times with lysis buffer, and sample buffer was added before subjecting to immunoblotting as described above. The protein-transferred membrane was probed individually for WASH1, Strumpellin, and HA. Data were collected from four separate preparations of WT and MUT conditions.

### Electron Microscopy

Adult (7mo) WT and MUT SWIP^P1019R^ mice were deeply anesthetized with isoflurane and then transcardially perfused with warmed heparinized saline (25U/ml heparin) for 4 minutes, followed by ice-cold 0.15M cacodylate buffer pH 7.4 containing 2.5% glutaraldehyde (Electron Microscopy Sciences #16320), 3% paraformaldehyde, and 2mM CaCl_2_ for 15 minutes. Brain samples were dissected and stored on ice in the same fixative for 2 hours before washing in 0.1M sodium cacodylate buffer (3 changes for 15 minutes each). Samples were then post-fixed in 1.0% OsO_4_ in 0.1 M Sodium cacodylate buffer for 1 hour on a rotator. Samples were then washed in 3, 15-minute changes of 0.1M sodium cacodylate. Samples were then placed into *en bloc* stain (1% uranyl acetate) overnight at 4°C. Subsequently, samples were dehydrated in a series of ascending acetone concentrations including 50%, 70%, 95%, and 100% for three cycles with 15 minutes incubation at each concentration change. Samples were then placed in a 50:50 mixture of epoxy resin (Epon) and acetone overnight on a rotator. This solution was then replaced twice with 100% fresh Epon for at least 2 hours at room temperature on a rotator. Samples were embedded with 100% Epon resin in BEEM capsules (Ted Pella) for 48 hours at 60°C. Samples were ultrathin sectioned to 60-70nm on a Reichert Ultracut E ultramicrotome. Harvested grids were then stained with 2% uranyl acetate in 50% ethanol for 30 minutes and Sato’s lead stain for 1 min. Micrographs were acquired using a Phillips CM12 electron microscope operating at 80Kv, at 1700x magnification. Micrographs were analyzed in Adobe Photoshop 2019, using the “magic wand” tool to demarcate and measure the area of electron-dense and electron-lucent regions of interest (ROIs). Statistical analyses of ROI measurements were performed in GraphPad Prism (version 8) software. The experimenter was blinded to genotype for image acquisition and analysis.

### iBioID Sample Preparation

AAV2/9 viral probes, hSyn1-WASH1-BioID2-HA or hSyn1-solubleBioID2-HA, were injected into wild-type CD1 mouse brains using a Hamilton syringe (#7635-01) at age P0-P1 to ensure viral spread throughout the forebrain(Glascock et al., 2011). 15 days post-viral injection, biotin was subcutaneously administered at 24mg/kg for seven consecutive days for biotinylation of proteins in proximity to BioID2 probes. Whole brains were extracted on the final day of biotin injections, snap frozen, and stored in liquid nitrogen until protein purification. Seven brains were used for protein purification of each probe, and each purification was performed three times independently (21 brains total for WASH1-BioID2, 21 for solubleBioID2).

We performed all homogenization and protein purification on ice. A 2ml Dounce homogenizer was used to individually homogenize each brain in a 1:1 solution of Lysis-R:2X-RIPA buffer solution with protease inhibitors (Roche cOmplete tablets #11836153001). Each sample was sonicated three times for 7 seconds and then centrifuged at 5000g for 5min at 4°C. Samples were transferred to Beckman Coulter 1.5ml tubes (#344059), and then spun at 45,000rpm in a Beckman Coulter tabletop ultracentrifuge (TLA-55 rotor) for 1hr at 4°C. SDS was added to supernatants (final 1%) and samples were then boiled for 5min at 95°C. We next combined supernatants from the same condition together (WASH1-BioID2 vs. solubleBioID2) in 15ml conical tubes to rotate with 30µl high-capacity NeutrAvidin beads overnight at 4°C (Thermo #29204).

The following day, all steps were performed under a hood with keratin-free reagents. Samples were spun down at 6000rpm, 4°C for 5min to pellet the beads and remove supernatant. The pelleted beads then went through a series of washes, each for 10 min at RT with 500ul of solvent, and then spun down on a tabletop centrifuge to pellet the beads for the next wash. The washes were as follows: 2% SDS twice, 1% TritonX100-1%deoxycholate-25mM LiCl2 once, 1M NaCL twice, 50mM Ammonium Bicarbonate (Ambic) five times. Beads were then mixed 1:1 with a 2X Laemmli sample buffer that contained 3mM biotin/50mM Ambic, boiled for 5 mins at 95°C, vortexed three times, and then biotinylated protein supernatants were stored at −80°C until LC-MS/MS.

### LC-MS/MS for iBioID

We gave the Duke Proteomics and Metabolomics Shared Resource (DPMSR) six eluents from streptavidin resins (3 x WASH1-BioID2, 3 x solubleBioID2), stored on dry ice. Samples were reduced with 10 mM dithiolthreitol for 30 min at 80°C and alkylated with 20 mM iodoacetamide for 30 min at room temperature. Next, samples were supplemented with a final concentration of 1.2% phosphoric acid and 256 μL of S-Trap (Protifi) binding buffer (90% MeOH/100mM TEAB). Proteins were trapped on the S-Trap, digested using 20 ng/μl sequencing grade trypsin (Promega) for 1 hr at 47°C, and eluted using 50 mM TEAB, followed by 0.2% FA, and lastly using 50% ACN/0.2% FA. All samples were then lyophilized to dryness and resuspended in 20 μL 1%TFA/2% acetonitrile containing 25 fmol/μL yeast alcohol dehydrogenase (UniProtKB P00330; ADH_YEAST). From each sample, 3 μL was removed to create a pooled QC sample (SPQC) which was run analyzed in technical triplicate throughout the acquisition period. Quantitative LC/MS/MS was performed on 2 μL of each sample, using a nanoAcquity UPLC system (Waters) coupled to a Thermo QExactive HF-X high resolution accurate mass tandem mass spectrometer (Thermo) via a nanoelectrospray ionization source. Briefly, the sample was first trapped on a Symmetry C18 20 mm × 180 μm trapping column (5 μl/min at 99.9/0.1 v/v water/acetonitrile), after which the analytical separation was performed using a 1.8 μm Acquity HSS T3 C18 75 μm × 250 mm column (Waters) with a 90-min linear gradient of 5 to 30% acetonitrile with 0.1% formic acid at a flow rate of 400 nanoliters/minute (nL/min) with a column temperature of 55°C. Data collection on the QExactive HF-X mass spectrometer was performed in a data-dependent acquisition (DDA) mode of acquisition with a r=120,000 (@ m/z 200) full MS scan from m/z 375 – 1600 with a target AGC value of 3e^6^ ions followed by 30 MS/MS scans at r=15,000 (@ m/z 200) at a target AGC value of 5e^4^ ions and 45 ms. A 20s dynamic exclusion was employed to increase depth of coverage. The total analysis cycle time for each sample injection was approximately 2 hours.

### LOPIT-DC Subcellular Fractionation

We performed three independent fractionation experiments with one adult SWIP mutant brain and one WT mouse brain fractionated in each experiment. Each mouse was sacrificed by isoflurane inhalation and its brain was immediately extracted and placed into a 2ml Dounce homogenizer on ice with 1ml isotonic TEVP homogenization buffer (320mM sucrose, 10mM Tris base, 1mM EDTA, 1mM EGTA, 5mM NaF, pH7.4 (Hallett et al., 2008)). A cOmplete mini protease inhibitor cocktail tablet (Sigma #11836170001) was added to a 50ml TEVP buffer aliquot immediately before use. Brains were homogenized for 15 passes with a Dounce homogenizer to break the tissue, and then this lysate was brought up to a 5ml volume with additional TEVP buffer. Lysates were then passed through a 0.5ml ball-bearing homogenizer for two passes (14 µm ball, Isobiotec) to release organelles. Final brain lysate volumes were approximately 7.5ml each. Lysates were then divided into replicate microfuge tubes (Beckman Coulter #357448) to perform differential centrifugation, following Geladaki et. al’s LOPIT-DC protocol(Geladaki et al., 2019). Centrifugation was carried out at 4°C in a tabletop Eppendorf 5424 centrifuge for spins at 200g, 1,000g, 3,000g, 5,000g, 9,000g, 12,000g, and 15,000g. To isolate the final three fractions, a tabletop Beckman TLA-100 ultracentrifuge with a TLA-55 rotor was used at 4°C with speeds of: 30,000g, 79,000g, and 120,000g, respectively. Samples were kept on ice at all times and pellets were stored at −80°C. Pellets from seven fractions (5,000g-120,000g) were used for proteomic analyses.

### 16-plex TMT LC-MS/MS

The Duke Proteomics and Metabolomics Shared Resource (DPMSR) processed and prepared fraction pellets from all 42 frozen samples simultaneously (7 fractions per brain from 3 WT and 3 MUT brains). Due to volume constraints, each sample was split into 3 tubes, for a total of 126 samples, which were processed in the following manner: 100µL of 8M Urea was added to the first aliquot then probe sonicated for 5 seconds with an energy setting of 30%. This volume was then transferred to the second and then third aliquot after sonication in the same manner. All tubes were centrifuged at 10,000g and any residual volume from tubes 1 and 2 were added to tube 3. Protein concentrations were determined by BCA on the supernatant in duplicate (5 μL each assay). Total protein concentrations for each replicate ranged from 1.1 mg/mL to 7.8 mg/mL with total protein quantities ranging from 108.3 to 740.81 µg. 60 µg of each sample was removed and normalized to 52.6µL with 8M Urea and 14.6µL 20% SDS. Samples were reduced with 10 mM dithiolthreitol for 30 min at 80°C and alkylated with 20 mM iodoacetamide for 30 min at room temperature. Next, they were supplemented with 7.4 μL of 12% phosphoric acid, and 574 μL of S-Trap (Protifi) binding buffer (90% MeOH/100mM TEAB). Proteins were trapped on the S-Trap, digested using 20 ng/μl sequencing grade trypsin (Promega) for 1 hr at 47°C, and eluted using 50 mM TEAB, followed by 0.2% FA, and lastly using 50% ACN/0.2% FA. All samples were then lyophilized to dryness.

Each sample was resuspended in 120 μL 200 mM triethylammonium bicarbonate, pH 8.0 (TEAB). From each sample, 20µL was removed and combined to form a pooled quality control sample (SPQC). Fresh TMTPro reagent (0.5 mg for each 16-plex reagent) was resuspended in 20 μL 100% acetonitrile (ACN) and was added to each sample. Samples were incubated for 1 hour at RT. After the 1-hour reaction, 5 μL of 5% hydroxylamine was added and incubated for 15 minutes at room temperature to quench the reaction. Each 16-plex TMT experiment consisted of the WT and MUT fractions from one mouse, as well as the 2 SPQC samples. Samples corresponding to each experiment were concatenated and lyophilized to dryness.

Samples were resuspended in 800µL 0.1% formic acid. 400µg was fractionated into 48 unique high pH reversed-phase fractions using pH 9.0 20 mM Ammonium formate as mobile phase A and neat acetonitrile as mobile phase B. The column used was a 2.1 mm x 50 mm XBridge C18 (Waters) and fractionation was performed on an Agilent 1100 HPLC with G1364C fraction collector. Throughout the method, the flow rate was 0.4 mL/min and the column temperature was 55°C. The gradient method was set as follows: 0 min, 3%B; 1 min, 7% B; 50 min, 50%B; 51 min, 90% B; 55 min, 90% B; 56 min, 3% B; 70 min, 3% B. 48 fractions were collected in equal time segments from 0 to 52 minutes, then concatenated into 12 unique samples using every 12th fraction. For instance, fraction 1, 13, 25, and 37 were combined, fraction 2, 14, 26, and 38 were combined, etc. Fractions were frozen and lyophilized overnight. Samples were resuspended in 66 μL 1%TFA/2% acetonitrile prior to LC-MS analysis.

Quantitative LC/MS/MS was performed on 2 μL (1 μg) of each sample, using a nanoAcquity UPLC system (Waters) coupled to a Thermo Orbitrap Fusion Lumos high resolution accurate mass tandem mass spectrometer (Thermo) equipped with a FAIMS Pro ion-mobility device via a nanoelectrospray ionization source. Briefly, the sample was first trapped on a Symmetry C18 20 mm × 180 μm trapping column (5 μl/min at 99.9/0.1 v/v water/acetonitrile), after which the analytical separation was performed using a 1.8 μm Acquity HSS T3 C18 75 μm × 250 mm column (Waters) with a 90-min linear gradient of 5 to 30% acetonitrile with 0.1% formic acid at a flow rate of 400 nanoliters/minute (nL/min) with a column temperature of 55°C. Data collection on the Fusion Lumos mass spectrometer was performed for three different compensation voltages (CV: −40v, −60v, −80v). Within each CV, a data-dependent acquisition (DDA) mode of acquisition with a r=120,000 (@ m/z 200) full MS scan from m/z 375 – 1600 with a target AGC value of 4e^5^ ions was performed. MS/MS scans were acquired in the Orbitrap at r=50,000 (@ m/z 200) from m/z 100 with a target AGC value of 1e^5^ and max fill time of 105 ms. The total cycle time for each CV was 1s, with total cycle times of 3 sec between like full MS scans. A 45s dynamic exclusion was employed to increase depth of coverage. The total analysis cycle time for each sample injection was approximately 2 hours.

Following UPLC-MS/MS analyses, data were imported into Proteome Discoverer 2.4 (Thermo Scientific). The MS/MS data were searched against a SwissProt Mouse database (downloaded November 2019) plus additional common contaminant proteins, including yeast alcohol dehydrogenase (ADH), bovine casein, bovine serum albumin, as well as an equal number of reversed-sequence “decoys” for FDR determination. Mascot Distiller and Mascot Server (v 2.5, Matrix Sciences) were utilized to produce fragment ion spectra and to perform the database searches. Database search parameters included fixed modification on Cys (carbamidomethyl) and variable modification on Met (oxidation), Asn/Gln (deamindation), Lys (TMTPro) and peptide N-termini (TMTPro). Data were searched at 5 ppm precursor and 0.02 product mass accuracy with full trypsin enzyme rules. Reporter ion intensities were calculated using the Reporter Ions Quantifier algorithm in Proteome Discoverer. Percolator node in Proteome Discoverer was used to annotate the data at a maximum 1% protein FDR.

### Mouse Behavioral Assays

Behavioral tests were performed on age-matched WT and homozygous SWIP^P1019R^ mutant littermates. Male and female mice were used in all experiments. Testing was performed at two time points: P42-55 days old as a young adult age, and 5.5 months old as mid-adulthood, so that we could compare disease progression in this mouse model to human patients(Ropers et al., 2011). The sequence of behavioral testing was: Y-maze (to measure working memory), object novelty recognition (to measure short-term and long-term object recognition memory), TreadScan (to assess gait), and steady-speed rotarod (to assess motor control and strength) for 40-55 day old mice. Testing was performed over 1.5 weeks, interspersed with rest days for acclimation. This sequence was repeated with the same cohort at 5.5-6 months old, with three additional measures added to the end of testing: fear conditioning (to assess associative fear memory), a hearing test (to measure tone response), and a shock threshold test (to assess somatosensation). Of note, a separate, second cohort of mice was evaluated for fear conditioning, hearing, and shock threshold testing at adolescence. After each trial, equipment was cleaned with Labsan to remove residual odors. The experimenter was blinded to genotype for all behavioral analyses.

### Y-maze

Working memory was evaluated by measuring spontaneous alternations in a 3-arm Y-maze under indirect illumination (80-90 lux). A mouse was placed in the center of the maze and allowed to freely explore all arms, each of which had different visual cues for spatial recognition. Trials were 5 min in length, with video data and analyses captured by EthoVision XT 11.0 software (Noldus Information Technology). Entry to an arm was define as the mouse being >1 body length into a given arm. An alternation was defined as three successive entries into each of the different arms. Total % alternation was calculated as the total number of alternations/the total number of arm entries minus 2 ×100.

### Novel Object Recognition

One hour before testing, mice were individually exposed to the testing arena (a 48 x 22 x 18cm white opaque arena) for 10min under 80-100lux illumination without any objects. The test consisted of three phases: training (day 1), short-term memory test (STM, day 1), and long-term memory test (LTM, day 2). For the training phase, two identical objects were placed 10 cm apart, against opposing walls of the arena. A mouse was placed in the center of the arena and given full access to explore both objects for 5 min and then returned to its home cage. For STM testing, one of the training objects remained (the now familiar object), and a novel object replaced one of the training objects (similar in size, different shape). The mouse was returned to the arena 30 minutes after the training task and allowed to explore freely for 5 mins. For LTM testing, the novel object was replaced with another object, and the familiar object remained unchanged. The LTM test was also 5 min in duration, conducted 24hr after the training task. Behavior was scored using Ethovision 11.0 XT software (Noldus) and analyzed by a blind observer. Object contact was defined as the mouse’s nose within 1 cm of the object. We analyzed both number of nose contacts with each object and duration of contacts. Preference scores were calculated as (duration contact_novel_ - duration contact_familiar_) / total duration contact_novel+familiar_. Positive scores signified a preference for the novel object; whereas, negative scores denoted a preference for the familiar object, and scores approaching zero indicated no preference.

### TreadScan

A TreadScan forced locomotion treadmill system (CleversSys Inc, Reston, Virginia) was used for gait recording and analysis. Each mouse was recorded walking on a transparent treadmill at 45 days old, and again at 5.5 months old. Mice were acclimated to the treadmill chamber for 1 minute before the start of recording to eliminate exploratory behavior confounding normal gait. Trials were 20 seconds in length, with mice walking at speeds between 13.83 and 16.53 cm/sec (P45 WT average 15.74 cm/s; P45 MUT average 15.80 cm/s; 5.5mo WT average 15.77 cm/s; 5.5mo MUT average 15.85 cm/s). A high-speed digital camera attached to the treadmill captured limb movement at a frame rate of 100 frames/second. We used TreadScan software (CleversSys) and representative WT and MUT videos to generate footprint templates, which were then used to identify individual paw profiles for each limb. Parameters such as stance time, swing time, step length, track width, and limb coupling were recorded for the entire 20 sec duration for each animal. Output gait tracking was verified manually by a blinded experimenter to ensure consistent limb tracking throughout the duration of each video.

### Steady Speed Rotarod

A 5-lane rotarod (Med Associates, St. Albans, VT) was used for steady-speed motor analysis. The rod was run at a steady speed of 32rpm for four, 5-minute trials, with a 40-minute inter-trial interval. We recorded mouse latency to fall by infrared beam break, or manually for any mouse that completed two or more rotations on the rod without walking. Mice were randomized across lanes for each trial.

### Fear Conditioning

Animals were examined in contextual and cued fear conditioning as described by Rodriguiz and Wetsel(Rodriguiz and Wetsel, 2006). Two separate cohorts of mice were used testing the two age groups. A three-day testing paradigm was used to assess memory: conditioning on day 1, context testing 24-hr post-conditioning on day 2, and cued tone testing 48hr post-conditioning on day 3. All testing was conducted in fear conditioning chambers (Med Associates). In the conditioning phase, mice were first acclimated to the test chamber for two minutes under ∼100 lux illumination. Then a 2900Hz, 80dB tone (conditioned stimulus, CS) played for 30 sec, which terminated with a paired 0.4mA, 2 sec scrambled foot shock (unconditioned stimulus, US). Mice were removed from the chamber and returned to their home cage 30 sec later. In the context testing phase, mice were placed in the same conditioning chamber and monitored for freezing behavior for a 5 min trial period, in the absence of the CS and US. For cued tone testing, the chambers were modified to different dimensions and shapes, contained different floor and wall textures, and lighting was adjusted to 50 lux. Mice acclimated to the chamber for 2 min, and then the CS was presented continuously for 3 min. Contextual and cued fear memory was assessed by freezing behavior, captured by automated video software (CleversSys).

### Hearing Test

We tested mouse hearing using a startle platform (Med Associates) connected to Startle Pro Software in a sound-proof chamber. Mice were placed in a ventilated restraint cylinder connected to the startle response detection system to measure startle to each acoustic stimulus. After two minutes of acclimation, mice were assessed for an acoustic startle response to seven different tone frequencies, 2kHz, 3kHz, 4kHz, 8kHz, 12kHz, 16kHz, and 20kHz that were randomly presented three times each at four different decibels, 80, 100, 105, and 110dB, for a total of 84 trials. A random inter-trial interval of 15-60 seconds (average 30sec) was used to prevent anticipation of a stimulus. An animal’s reaction to the tone was recorded as startle reactivity in the first 100msec of the stimulus presentation, which was transduced through the platform’s load cell and expressed in arbitrary units (AU).

### Startle Response (Somatosensation)

Mouse somatosensation was tested by placing mice in a startle chamber (Med Associates) connected to Startle Pro Software. Mice were placed atop a multi-bar cradle within a ventilated plexiglass restraint cylinder, which allows for horizontal movement within the chamber, but not upright rearing. After two minutes of acclimation, each mouse was exposed to 10 different scrambled shock intensities, ranging from 0 to 0.6mA with randomized inter-trial intervals of 20-90 seconds. Each animal’s startle reactivity during the first 100 msec of the shock was transduced through the platform’s load cell and recorded as area under the curve (AUC) in arbitrary units (AU).

## QUANTIFICATION AND STATISTICAL ANALYSIS

Experimental conditions, number of replicates, and statistical tests used are stated in each figure legend. Each experiment was replicated at least three times (or on at least 3 separate animals) to assure rigor and reproducibility. Both male and female age-matched mice were used for all experiments, with data pooled from both sexes. Data compilation and statistical analyses for all non-proteomic data were performed using GraphPad Prism (version 8, GraphPad Software, CA), using a significance level of alpha=0.05. Prism provides exact p values unless p<0.0001. All data are reported as mean ± SEM. Each data set was tested for normal distribution using a D’Agostino-Person normality test to determine whether parametric (unpaired Student’s t-test, one-way ANOVA, two-way ANOVA) or non-parametric (Mann-Whitney, Kruskal-Wallis) tests should be used. Parametric assumptions were confirmed with the Shapiro-Wilk test (normality) and Levine’s test (error variance homogeneity) for ANOVA with repeated measures testing. The analysis of iBioID and TMT proteomics data are described below. All proteomic data and analysis scripts are available online (see Resource Availability).

### Imaris 3D reconstruction

For EEA1+ and CathepsinD+ puncta analyses, coverslips were imaged on a Zeiss LSM 710 confocal microscope. Images were sampled at a resolution of 1024 x 1024 pixels with a dwell time of 0.45µsec using a 63x/1.4 oil immersion objective, a 2.0 times digital zoom, and a z-step size of 0.37 µm. Images were saved as “.lsm” formatted files, and quantification was performed on a POGO Velocity workstation in the Duke Light Microscopy Core Facility using Imaris 9.2.0 software (Bitplane, South Windsor, CT). For analyses, we first used the “surface” tool to make a solid fill surface of the MAP2-stained neuronal soma and dendrites, with the background subtraction option enabled. We selected a threshold that demarcated the neuron structure accurately while excluding background. For EEA1 puncta analyses, a 600 x 800 µm selection box was placed around the soma in each image and surfaces were created for EEA1 puncta within the selection box. Similarly, for CathepsinD puncta analyses, a 600 x 600 µm selection box was placed around the soma(s) in each image for surface creation. The same threshold settings were used across all images, and individual surface data from each soma were exported for aggregate analyses. The experimenter was blinded to sample conditions for both image acquisition and analysis.

### Cleaved Caspase-3 Image Analysis

Z-stack images were acquired on a Zeiss 710 LSM confocal microscope. Images were sampled at a resolution of 1024 x 1024 pixels with a dwell time of 1.58µsec, using a 63x/1.4 oil immersion objective (for cortex, striatum, and hippocampus) or 20x/0.8 dry objective (cerebellum), a 1.0 times digital zoom, and a z-step size of 0.67 µm. Images were saved as “.lsm” formatted files, and then converted into maximum intensity projections (MIP) using Zen 2.3 SP1 software. Quantification of CC3 colocalization with DAPI was performed on the MIPs using the Particle Analyzer function in FIJI ImageJ software. The experimenter was blind to sample conditions for both image acquisition and analysis.

### iBioID Quantitative Analysis

Following UPLC-MS/MS analyses, data was imported into Proteome Discoverer 2.2 (Thermo Scientific Inc.), and aligned based on the accurate mass and retention time of detected ions (“features”) using Minora Feature Detector algorithm in Proteome Discoverer. Relative peptide abundance was calculated based on area-under-the-curve (AUC) of the selected ion chromatograms of the aligned features across all runs. The MS/MS data was searched against the SwissProt *Mus musculus* database (downloaded in April 2018) with additional proteins, including yeast ADH1, bovine serum albumin, as well as an equal number of reversed-sequence “decoys” for false discovery rate (FDR) determination. Mascot Distiller and Mascot Server (v 2.5, Matrix Sciences) were utilized to produce fragment ion spectra and to perform the database searches. Database search parameters included fixed modification on Cys (carbamidomethyl), variable modifications on Meth (oxidation) and Asn and Gln (deamidation), and were searched at 5 ppm precursor and 0.02 Da product mass accuracy with full trypsin enzymatic rules. Peptide Validator and Protein FDR Validator nodes in Proteome Discoverer were used to annotate the data at a maximum 1% protein FDR.

Protein intensities were exported from Proteome Discoverer and processed using custom R scripts. Carboxylases and keratins, as well as 315 mitochondrial proteins(Calvo et al., 2016), were removed from the identified proteins as known contaminants. Next, we performed sample loading normalization to account for technical variation between the 9 individual MS runs. This is done by multiplying intensities from each MS run by a scaling factor, such that the average of all total run intensities are equal. As QC samples were created by pooling equivalent aliquots of peptides from each biological replicate, the average of all biological replicates should be equal to the average of all technical SPQC replicates. We performed sample pool normalization to SPQC samples to standardize protein measurements across all samples and correct for batch effects between MS analyses. Sample pool normalization adjusts the protein-wise mean of all biological replicates to be equal to the mean of all SPQC replicates. Finally, proteins that were identified by a single peptide, and/or identified in less than 50% of samples were removed. Any remaining missing values were inferred to be missing not at random due to the left shifted distribution of proteins with missing values and imputed using the k-nearest neighbors algorithm using the impute.knn function in the R package impute (impute::impute.knn). Normalized protein data was analyzed using edgeR, an R package for the analysis of differential expression/abundance that models count data using a binomial distribution methodology. Differential enrichment of proteins in the WASH1-BioID2 pull-down relative to the solubleBioID2 control pull-down were evaluated with an exact test as implemented by the edgeR::exactTest function. To consider a protein enriched in the WASH interactome, we required that a protein exhibit a fold change greater than 3 over the negative control with an exact test Benjamini Hochberg adjusted p-value (FDR) less than 0.1. With these criteria, 174 proteins were identified as WASH1 interactome proteins. Raw peptide and final normalized protein data as well as the statistical results can be found in Table S1.

Proteins that function together often interact directly. We compiled experimentally-determined protein-protein interactions (PPIs) among the WASH1 interactome from the HitPredict database(López et al., 2015) using a custom R package, getPPIs, (available online at twesleyb/getPPIs). We report PPIs among the WASH1 interactome in Table S1. Bioinformatic GO analysis was conducted by manual annotation of identified proteins and confirmed with Metascape analysis(Zhou et al., 2019) of WASH1-BioID2 enriched proteins using the 2,311 proteins identified in the mass spec analysis as background.

Raw peptide intensities were exported from Proteome Discover for downstream analysis and processing in R. Following database searching, protein scoring using the Protein FDR Validator algorithm, and removal of contaminant species, the dataset retained 86,551 peptides corresponding to the identification of 7,488 unique proteins. These data, as well as statistical results can be found in Table S2.

### TMT Proteomics Quantitative Analysis

Peptide level data from the spatial proteomics analysis of SWIP^P1019R^ MUT and MUT brain were exported from Proteome Discoverer (version 2.4) and analyzed using custom R and Python scripts. Peptides from contaminant and non-mouse proteins were removed. First, we performed sample loading normalization, normalizing the total ion intensity for each TMT channel within an experiment to be equal. Sample loading normalization corrects for small differences in the amount of sample analyzed and labeling reaction efficiency differences between individual TMT channels within an experiment.

We found that in each TMT experiment there were a small number of missing values (mean percent missing = 1.6 +/-0.17%). Missing values were inferred to be missing at random based on the overlapping distributions of peptides with missing values and peptides without missing values. We imputed these missing values using the k-nearest neighbor algorithm (impute::impute.knn). Missing values for SPQC samples were not imputed. Peptides with any missing SPQC data were removed.

Following sample loading normalization, SPQC replicates within each experiment should yield identical measurements. As peptides with irreproducible QC measurements are unlikely to be quantitatively robust, and their inclusion may bias downstream processing (see IRS normalization below), we sought to remove them. To assess intra-batch variability, we utilized the method described by Ping et al., 2019(Ping et al., 2018). Briefly, peptides were binned into 5 groups based on the average intensity of the two SPQC replicates. For each pair of SPQC measurements, the log ratio of SPQC intensities was calculated. To identify outlier QC peptides, we plotted the distribution of these log ratios for each bin. Peptides with ratios that were more than four standard deviations away from the mean of its intensity bin were considered outliers and removed (Total number of SPQC outlier peptides removed = 474).

Proteins were summarized as the sum of all unique peptide intensities corresponding to a unique UniProtKB Accession identifier, and sample loading normalization was performed across all three experiments to account for inter-experimental technical variability. In a TMT experiment, the peptides selected for MS2 fragmentation for any given protein is partially random, especially at lower signal-to-noise peptides. This stochasticity means that proteins are typically quantified by different peptides in each experiment. Thus, although SPQC samples should yield identical protein measurements in each of the three experiments (as it is the same sample analyzed in each experiment), the observed protein measurements exhibit variability due to their quantification by different peptides. To account for this protein-level bias, we utilized the internal reference scaling (IRS) approach described by Plubell *et al*., 2017(Plubell et al., 2017). IRS normalization scales the protein-wise geometric average of all SPQC measurements across all experiments to be equal, and simultaneously adjusts biological replicates. In brief, each protein is multiplied by a scaling factor which adjusts its intra-experimental SPQC values to be equal to the geometric mean of all SPQC values for the three experiments. This normalization step effectively standardizes protein measurements between different mass spectrometry experiments.

The final normalization step was to perform sample pool normalization using SPQC samples as a reference. This normalization step, sometimes referred to as global internal standard normalization, accounts for batch effects between experiments, and reflects the fact that after technical normalization, the mean of biological replicates should be equal to the mean of SPQC replicates.

Before assessing protein differential abundance, we removed irreproducible proteins. This included proteins that were quantified in less than 50% of all samples, proteins that were identified by a single peptide, and proteins that had missing SPQC values. Across all 42 biological replicates, we observed that a small number of proteins had potential outlier measurements that were either several orders of magnitude greater or less than the mean of its replicates. In order to identify and remove these proteins, we assessed the reproducibility of protein measurements within a fraction in the same manner used to identify and filter SPQC outlier peptides. A small number of proteins were identified as outliers if the average log ratio of their 3 technical replicates was more than 4 standard deviations away from the mean of its intensity bin (n=349). In total, we retained 5,897 of the original 7,488 proteins in the final dataset.

Differential protein abundance was assessed using the final normalized protein data for intrafraction comparisons between WT and MUT groups using a general linear model as implemented by the edgeR::glmQLFit and edgeR::glmQLFTest functions(MD et al., 2009). Although this approach was originally devised for analysis of single-cell RNA-sequencing data, this approach is also appropriate for proteomics count data which is over-dispersed, negative binomially distributed, and often only includes a small number of replicates (for an example of edgeR’s application to proteomics see Plubell et al., 2017(Plubell et al., 2017))(McCarthy et al., 2012; MD et al., 2009). For intrafraction comparisons, P-values were corrected using the Benjamini Hochberg procedure within edgeR. An FDR threshold of 0.1 was set for significance for intrafraction comparisons.

We utilized edgeR’s flexible GLM framework to test the hypothesis that the abundance of proteins in the WT group was significantly different from that in the MUT group irrespective of fraction differences (Table S2). For WT vs. MUT contrasts, we considered proteins with an FDR < 0.05 significant (n=687). For plotting, we adjusted normalized protein abundances for fraction differences by fitting the data with an additive linear model with fraction as a blocking factor, as implemented by the removeBatchEffect algorithm from the R limma package(Ritchie et al., 2015).

To construct a protein covariation graph, we assessed the pairwise covariation (correlation) between all 5,897 proteins using the biweight midcorrelation (WGCNA::bicor) statistic(Seyfried et al., 2017), a robust alternative to Pearson’s correlation. The resulting complete, signed, weighted, and symmetric adjacency matrix was then re-weighted using the ‘Network Enhancement’ approach. Network enhancement removes noise from the graph, and facilitates downstream community detection(Wang et al., 2018).

The enhanced adjacency matrix was clustered using the Leiden algorithm(Traag et al., 2019), a recent extension and improvement of the well-known Louvain algorithm(Mucha et al., 2010). The Leiden algorithm functions to optimize the partition of a graph into modules by maximizing a quality statistic. We utilized the ‘Surprise’ quality statistic(Traag et al., 2015) to identify optimal partitions of the protein covariation graph. To facilitate biological inferences drawn from the network’s organization, we recursively split 27 modules that contained more than 100 nodes and removed modules that were smaller than 5 proteins. Initial clustering of the network resulted in the identification of 324 modules.

To reduce the likelihood of identifying false positive modules, we enforced module quality using a permutation procedure (NetRep::modulePreservation)(Ritchie et al., 2016) and removed modules with any insignificant permutation statistics (Bonferroni P-Adjust > 0.05). The following statistics were used to enforce module quality: ‘avg.weight’ (average edge weight), ‘avg.cor’ (average bicor correlation R^2^), and ‘avg.contrib’ (quantifies how similar an individual protein’s abundance profile is to the summary of its module). Proteins which were assigned to modules with insignificant module quality statistics were not considered clustered as the observed quality of their module does not differ from random. After filtering, approximately 85% of all proteins were assigned a cluster. The median percent variance explained by the first principle component of a module (a measure of module cohesiveness) was high (59.8%). After removal of low-quality modules, the analysis retained 255 distinct modules of proteins that strongly covaried together (Table S3).

To evaluate modules that were changing between WT and MUT genotypes, we extended the GLM framework to test for protein differential abundance. Modules were summarized as the sum of their proteins and fit with a GLM, with fraction as a blocking factor. In this statistical design, we were interested in the average effect of genotype on all proteins in a module. For plotting, module abundance was adjusted for fraction differences using the removeBatchEffect function (package: limma). We utilized the Bonferroni method to adjust P-values for 255 module level comparisons and considered modules with an adjusted P-value less than 0.05 were considered significant (n=37).

### Module Gene Set Enrichment Analysis

Modules were analyzed for enrichment of the WASH interactome (this paper), Retriever complex (McNally et al., 2017), CORUM protein complexes (Giurgiu et al., 2019), and subcellular predictions generated by Geladaki *et al*.(Geladaki et al., 2019) using the hypergeometric test with Bonferroni P-value correction for multiple comparisons. The union of all clustered and pathway proteins was used as background for the hypergeometric test. In addition to analysis of these general cellular pathways, we analyzed modules for enrichment of neuron-specific subcellular compartments—this included the presynapse (Takamori et al., 2006), excitatory postsynapse (Uezu et al., 2016), and inhibitory postsynapse (Uezu et al., 2016). These gene lists are available online at https://github.com/twesleyb/geneLists.

### Network Visualization

Network graphs were visualized in Cytoscape (Version 3.7.2). We used the Perfuse Force Directed Layout (weight = edge weight). In this layout, strongly connected nodes tend to be positioned closer together. In some instances, node location was manually adjusted to visualize the module more compactly. Node size was set to be proportional to the weighted degree centrality of a node in its module subgraph. Node size thus reflects node importance in the module. Visualizing co-expression or co-variation networks is challenging because every node is connected to every other node (the graph is complete). To aid visualization of module topology, we removed weak edges from the graphs. A threshold for each module was set to remove the maximal number of edges before the module subgraph split into multiple components. This strategy enables visualization of the strongest paths in a network.

## SUPPLEMENTARY FILES

- **Supplementary File 1:** Representative modules from major subcellular compartments.
- **Supplementary File 2:** Source data for Western blots.
- **Table S1:** WASH iBioID raw and normalized data and the corresponding statistical results.
- **Table S2:** SWIP^P1019R^ TMT raw and normalized proteomics data and the corresponding statistical results.
- **Table S3:** Module-level data and statistical results from network analysis of SWIP^P109R^ proteomics.
- **Table S4:** DNA oligonucleotides used in this study.
- **Table S5:** Key resources.

---

Supplementary File 1. Representative modules from major subcellular compartments.

**Figure 1.**
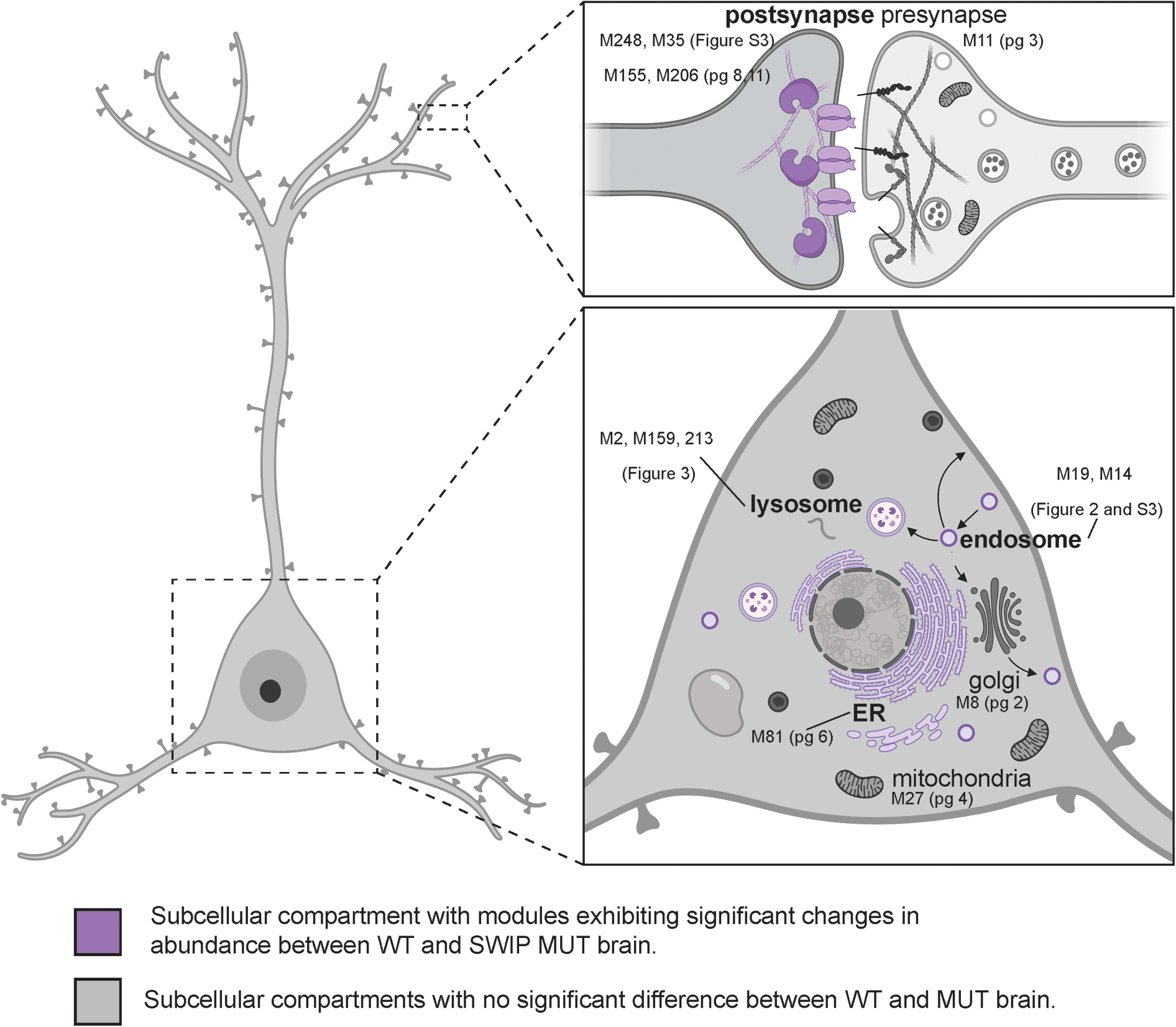
Schematic overview of major subcellular compartments in neurons. SWIP^P1019R^ MUT mouse brain exhibited significant differences in endosomal, lysosomal, ER, and postsynaptic modules compared to WT. Each of the following figures contains (A) normalized protein abundance for all proteins in a module, (B) schematic of the predicted subcellular compartment, and (C) network subgraph vizualization of a module. These modules, which do not exhibit a significant difference between WT and SWIP^P1019R^ brain, support the hypothesis that the primary cellular deficit resulting from SWIP^P1019R^ is dysregulation of endo-lysosomal pathways.

**Figure 2.**
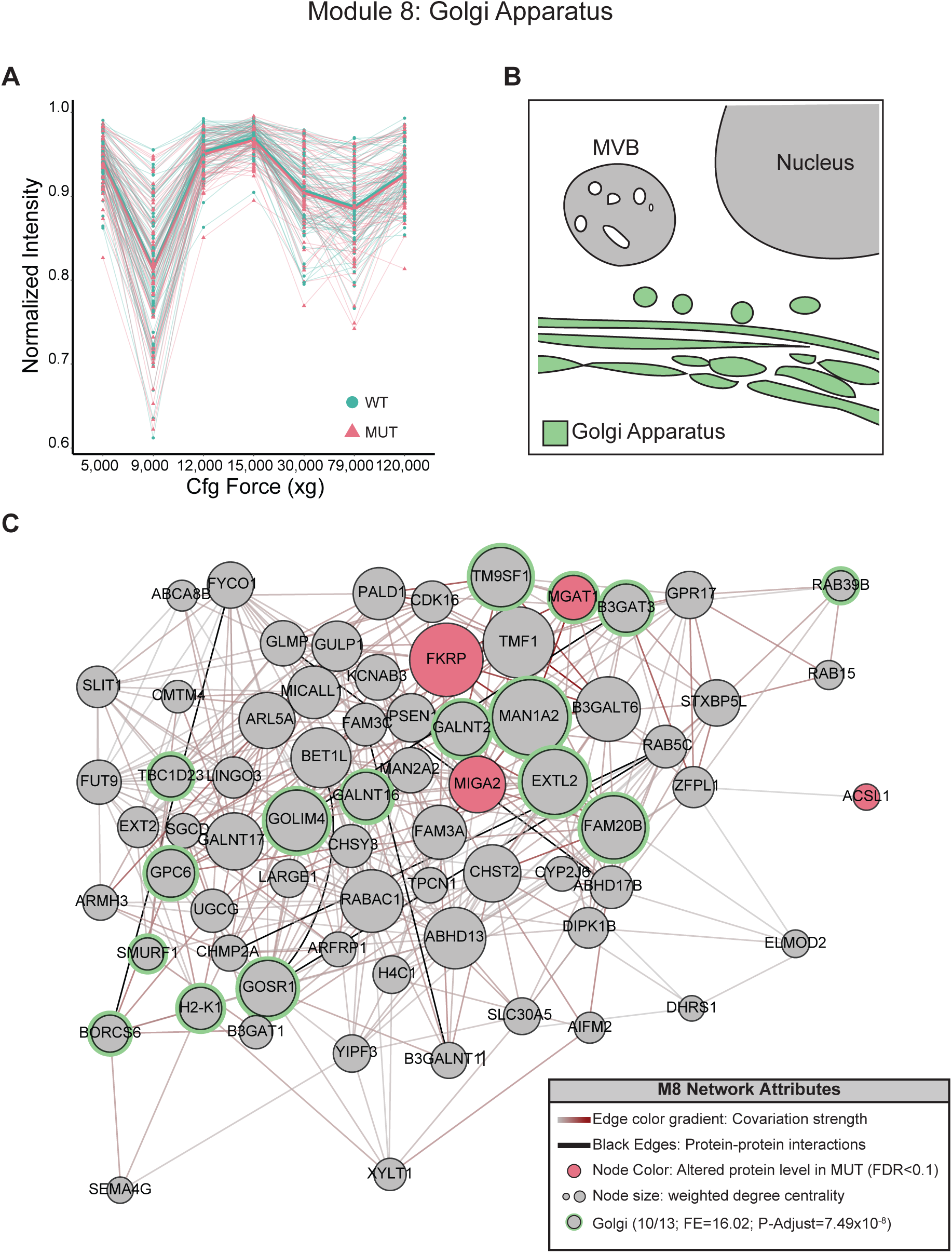
No significant difference in a predicted golgi apparatus module. (A) normalized protein abundance for all proteins in M8, which is predicted to represent the golgi apparatus based on its significant enrichment of golgi proteins from Geladaki *et al*., (2019). (B) schematic representation of the golgi apparatus. (C) M8 subnetwork showing organization of M8 proteins including 10 out of the 13 golgi proteins quantified in this study. Note the presence of MAN1A2 a mannosidase which resides in the golgi and functions in protein glycoslyation.

**Figure 3.**
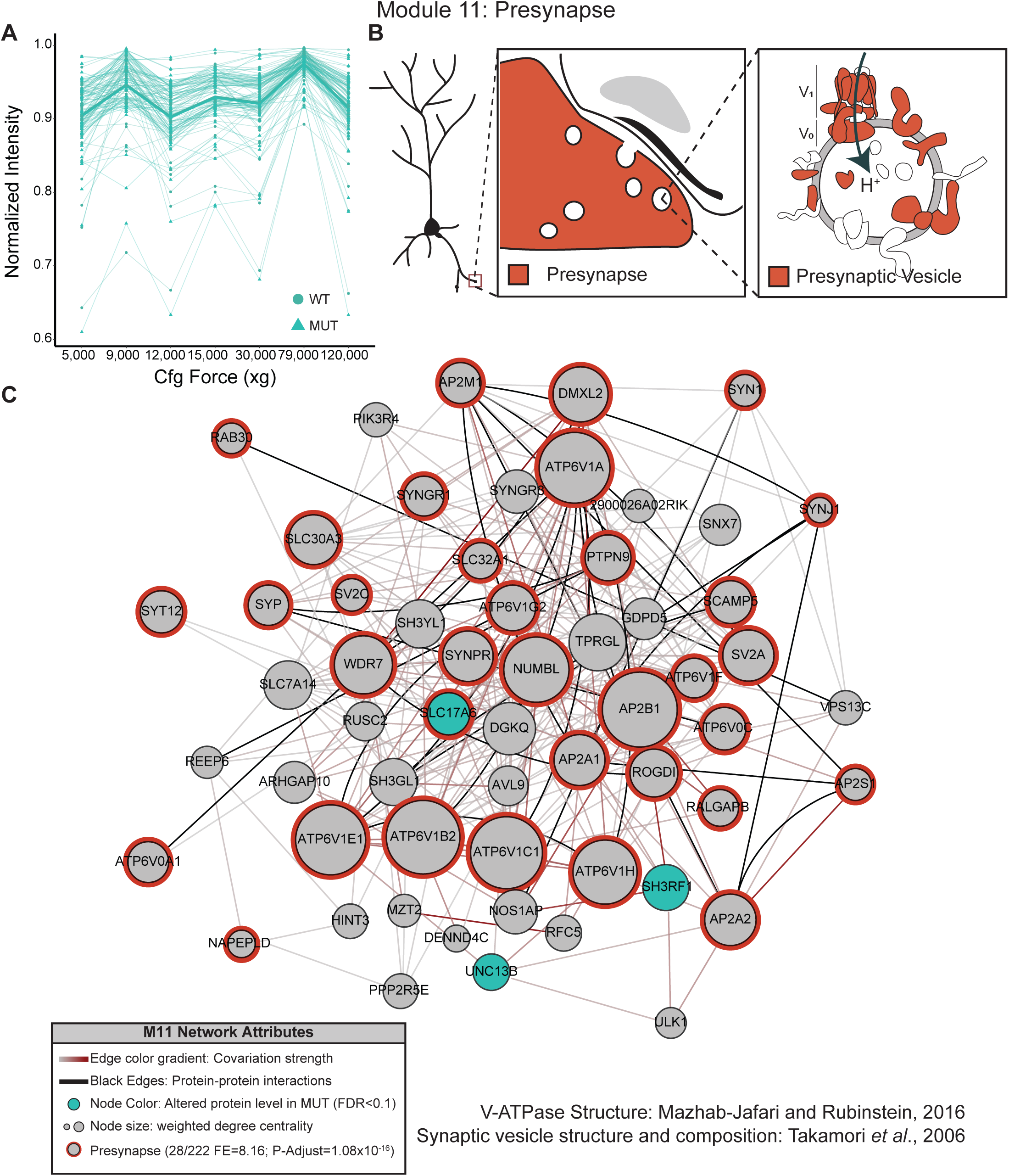
No significant difference in a predicted presynapse module. (A) normalized protein abundance for all proteins in M11, a module which is enriched for proteins known to reside in the presynaptic compartment or in presynaptic vesicles, of neurons (Takamori *et al*., 2006). (B) schematic of the vesicle-enriched presynaptic compartment and inset showing the protein composition of a presynaptic vesicle. (C) M11 network graph.

**Figure 4.**
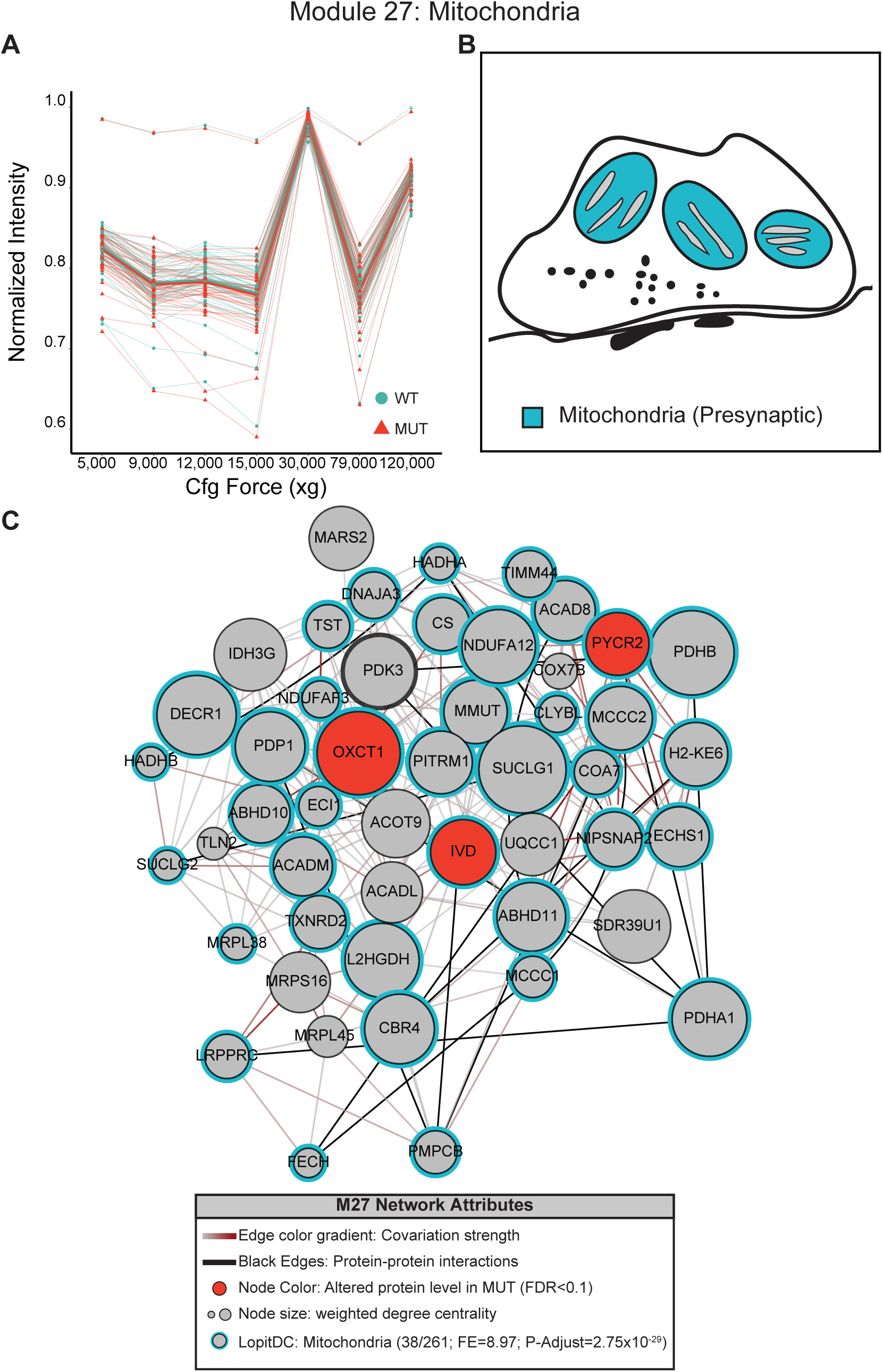
No significant difference in a predicted mitochondrial module. (A) normalized protein abundance for all proteins in M27, predicted to have mitochondrial function by its enrichment of mitochondrial proteins (Geladaki *et al*., 2019). (B) schematic of the neuronal mitochondria found in the presynapse. (C) M27 network which contains 38 of the of the 261 mitochondrial proteins quantified in this study.

**Figure 5.**
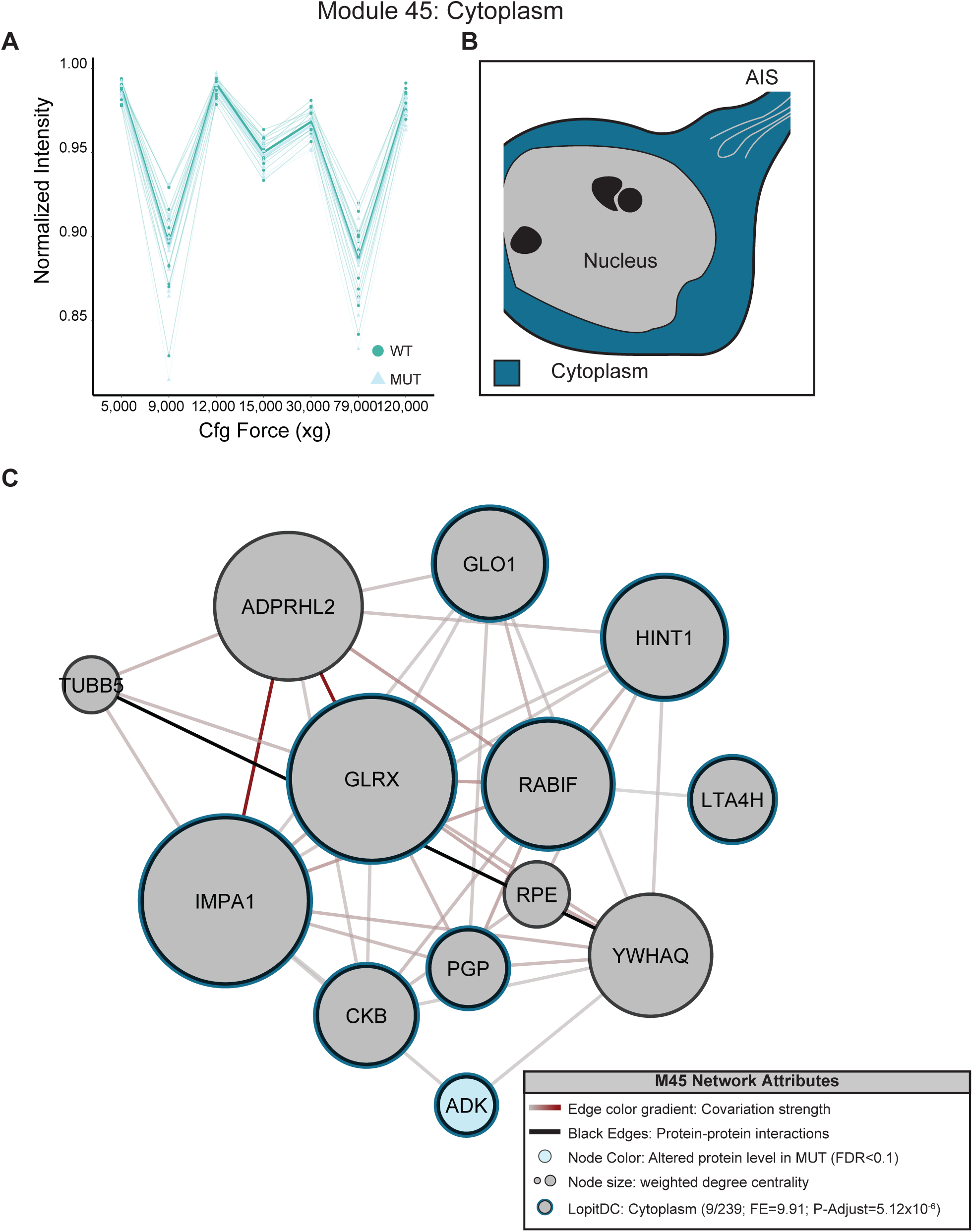
No significant difference in a predicted cytoplasmic module. (A) normalized protein abundance for all proteins in M45, predicted to have cytoplasmic localization. (B) schematic of the predicted cytoplasmic compartment, and (C) M45 network. M45 contained 9 of the 239 cytoplasmic proteins (Geladaki *et al*., 2019) quantified in this study.

**Figure 6.**
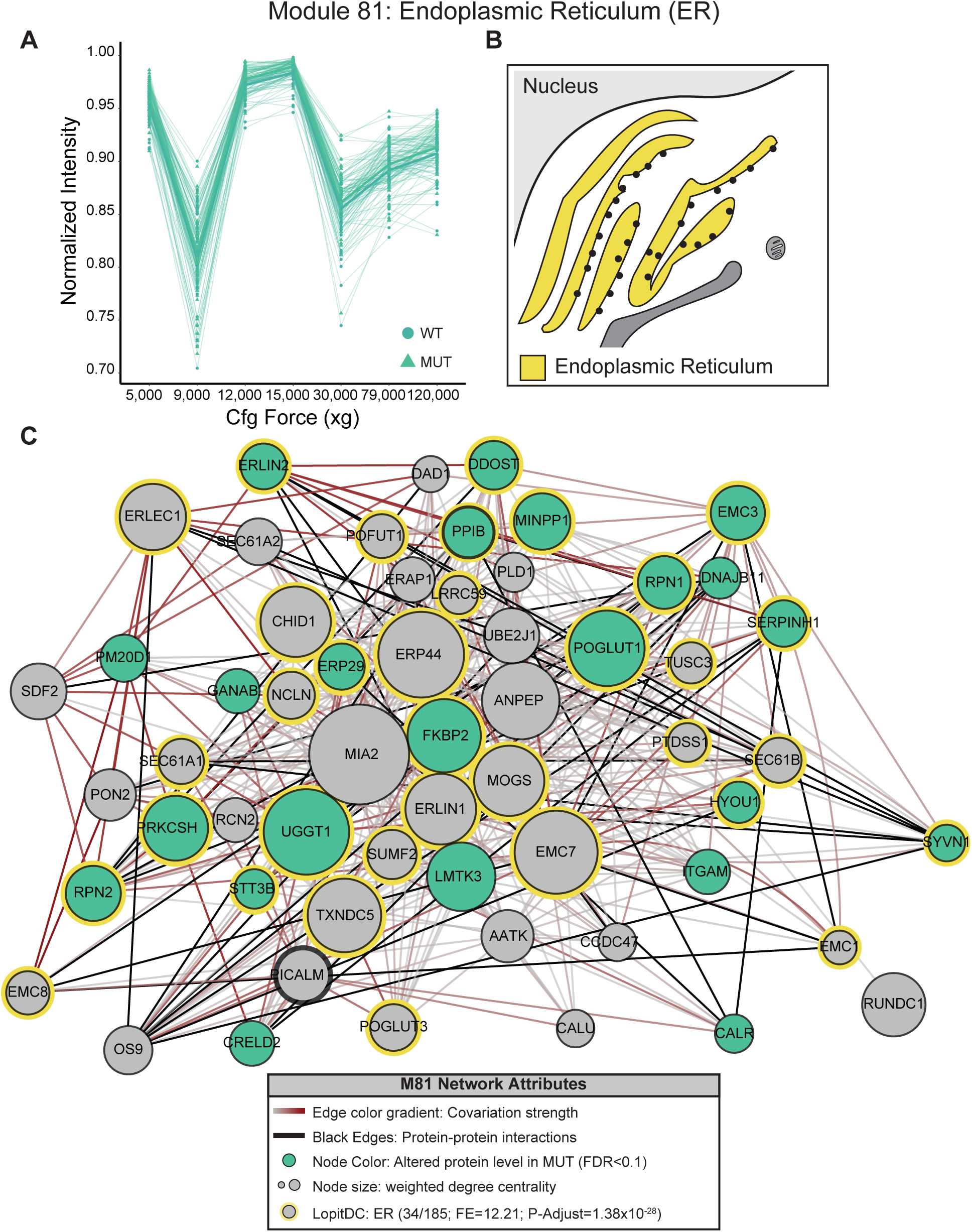
No significant difference in a predicted ER module. (A) normalized protein abundance for all proteins in M81, predicted to represent the endoplasmic reticulum (ER) based on its enrichment of ER proteins (Geladaki et al., 2006). Note that unlike M83 (Fig. S3), there is no significant difference between MUT WT brain. (B) Schematic of the predicted ER compartment, and (C) M81 network. M81 contains 34 ER proteins incuding Erlin1/2 which function in a complex as a part of the ERAD pathway.

**Figure 7.**
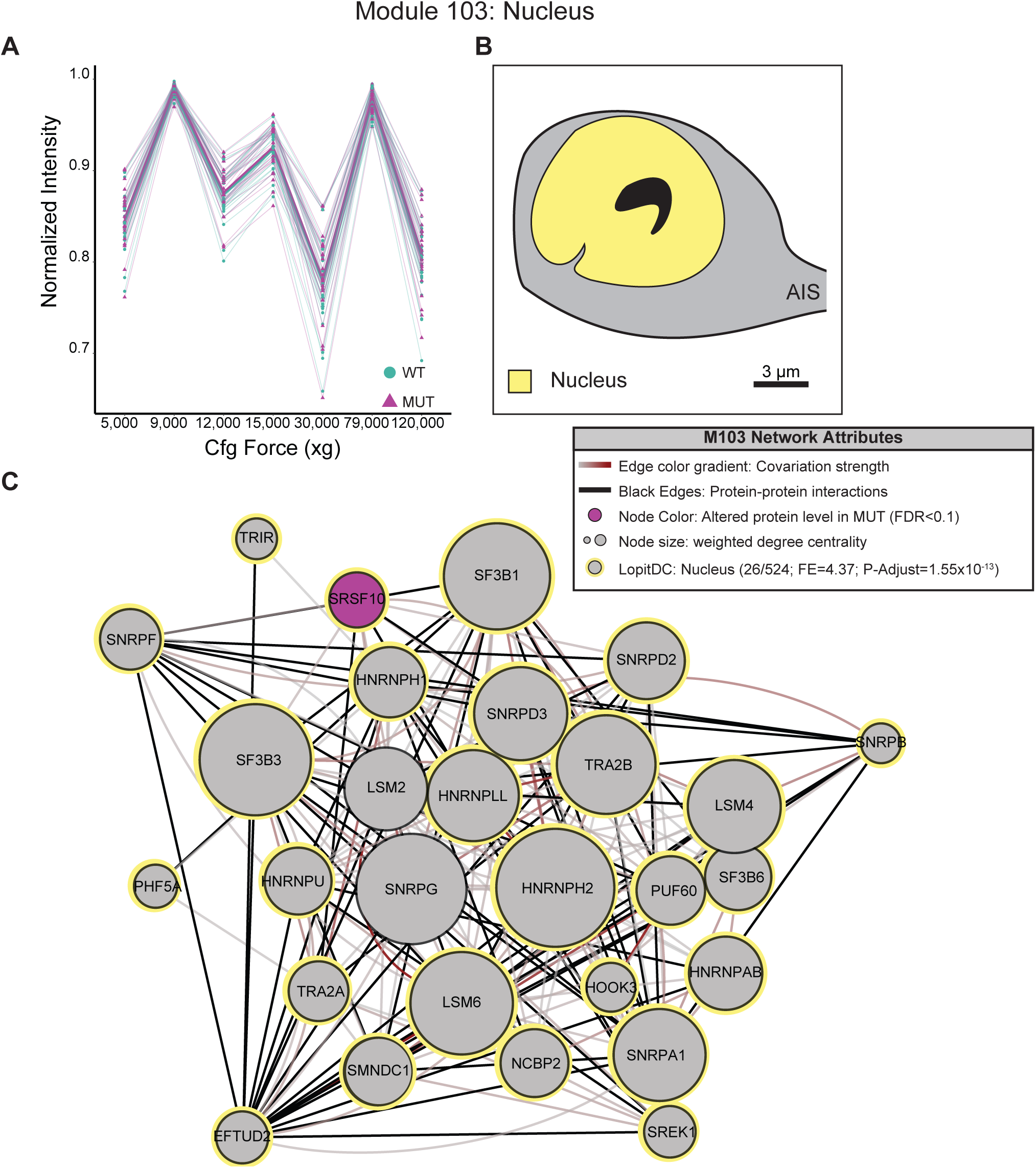
No significant difference in a predicted nuclear module. (A) normalized protein abundance for all proteins in M103, a predicted nuclear module. (B) schematic of the predicted nuclear compartment, and (C) M103 network. M103 contains 26 resident nuclear proteins, including LSM4 and LSM4, which function in spliceosome assembly.

**Figure 8.**
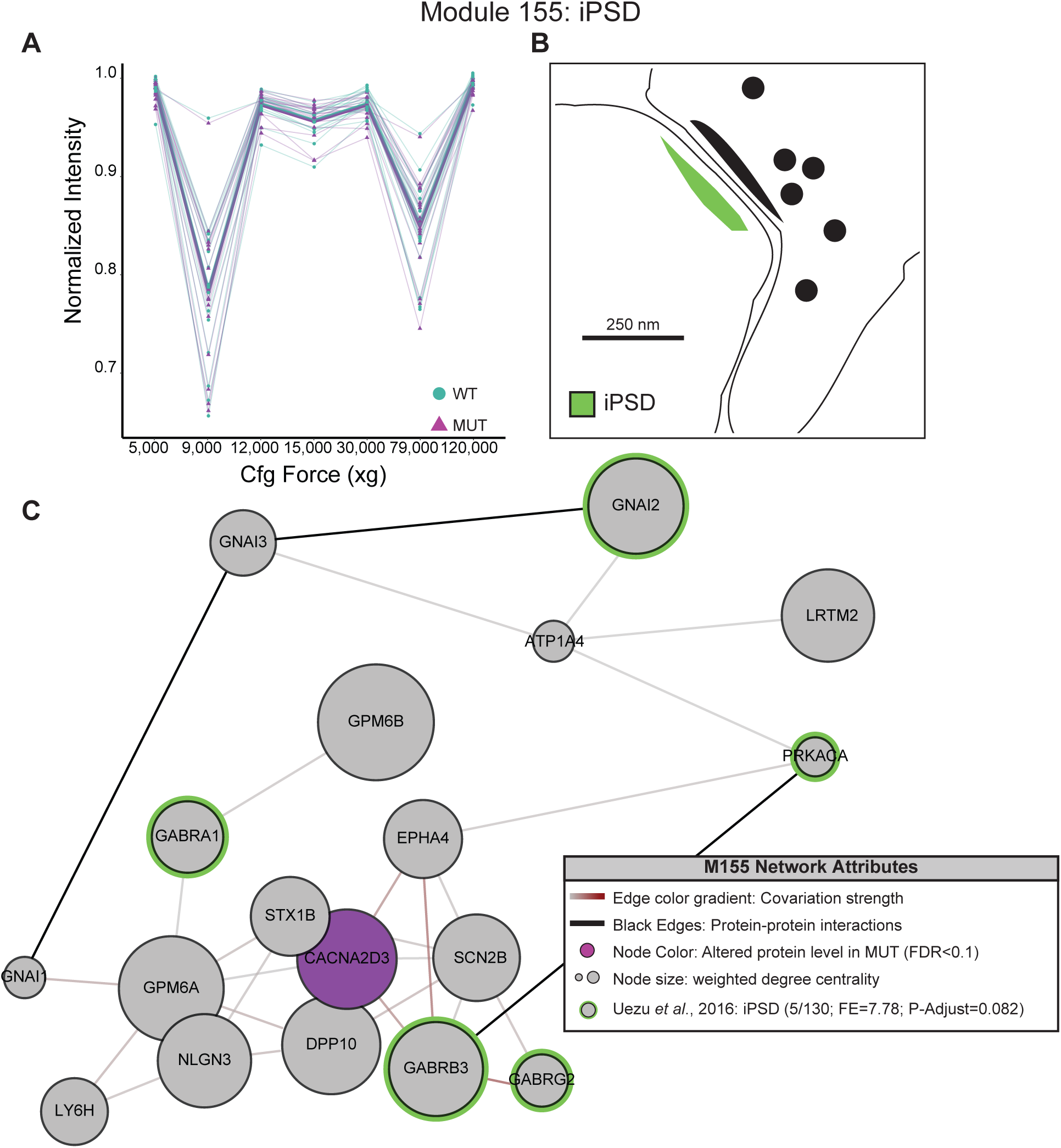
No significant difference in a predicted inhibitory postsynaptic density (iPSD) module. (A) normalized protein abundance for all proteins in M155, a predicted iPSD module. (B) schematic of the predicted iPSD compartment, and (C) M155 network. M155 contains 5 iPSD proteins including two GABA_A_R subunits and one GABA_B_R subunit identified by Uezu *et al*., 2016.

**Figure 9.**
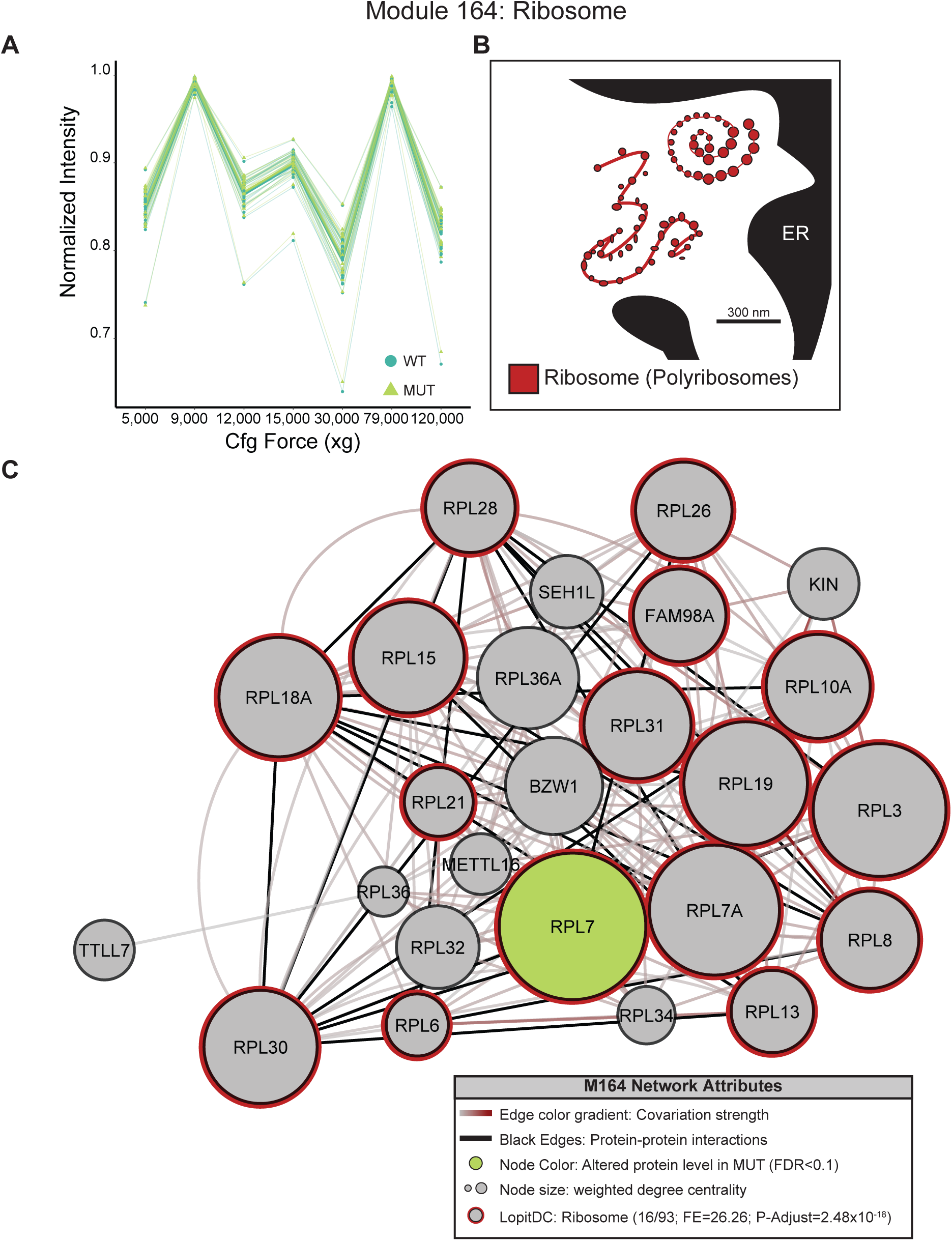
No significant difference in a predicted ribosome module. (A) normalized protein abundance for all proteins in M164, predicted to have ribosomal function by enrichment of ribosome proteins (Geladaki *et al*., 2019). (B) schematic of the predicted ribosomal compartment, and (C) M164 network.

**Figure 10.**
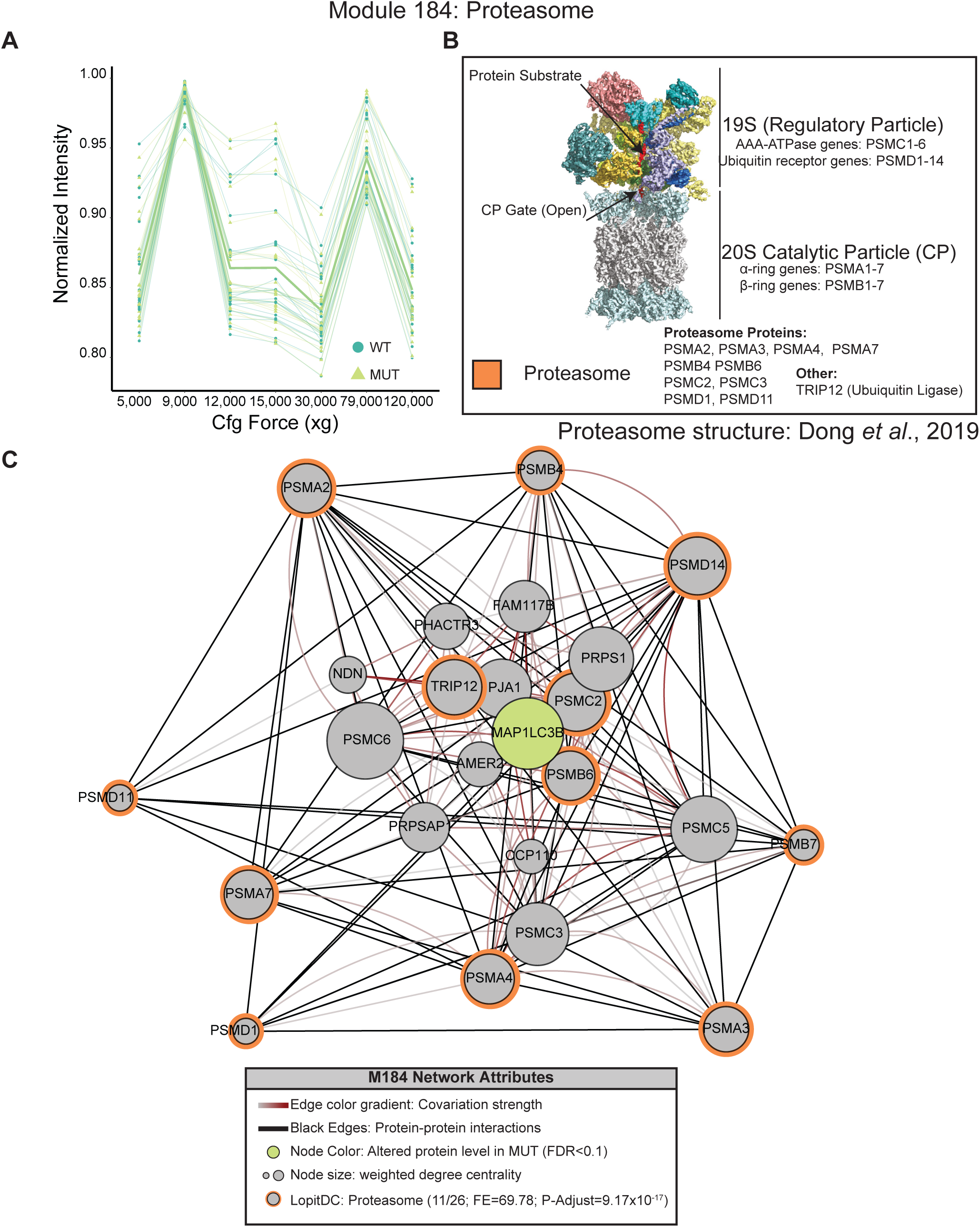
No significant difference in a predicted proteosome module. (A) normalized protein abundance for all proteins in M184, predicted to represent the proteasome. (B) schematic of the predicted proteosomal compartment, and (C) M184 network. M184 contained 11 out of the 26 proteosomal proteins from Geladaki *et al*., (2019) that were quantified in this study.

**Figure 11.**
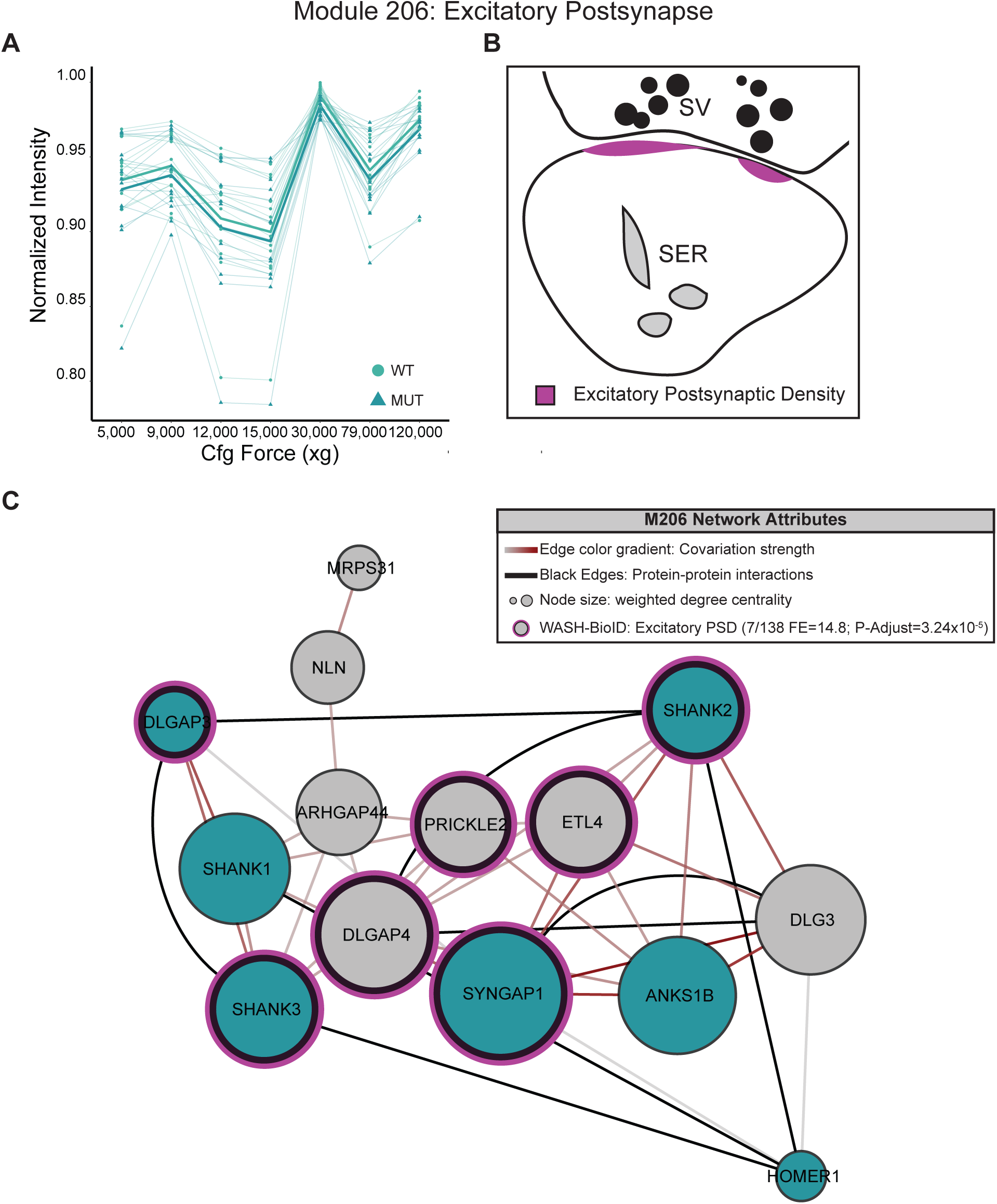
No significant difference in a predicted excitatory postsynapse (ePSD) module. (A) normalized protein abundance for all proteins in M206, predicted to have postsynaptic function by enrichment of proteins known to reside in the excitatory postsynapse. (B) schematic of the predicted ePSD compartment, and (C) M206 network. M206 contained 7 out of the 138 ePSD proteins reported by Uezu *et al*., (2016) and is also enriched for proteins identified by WASH-BioID (this study). In contrast to modules M35 and M143, which exhibited decreased abundance in SWIP MUT brain, M206 does not exhibit a global difference between WT and MUT brain.

**Figure 12.**
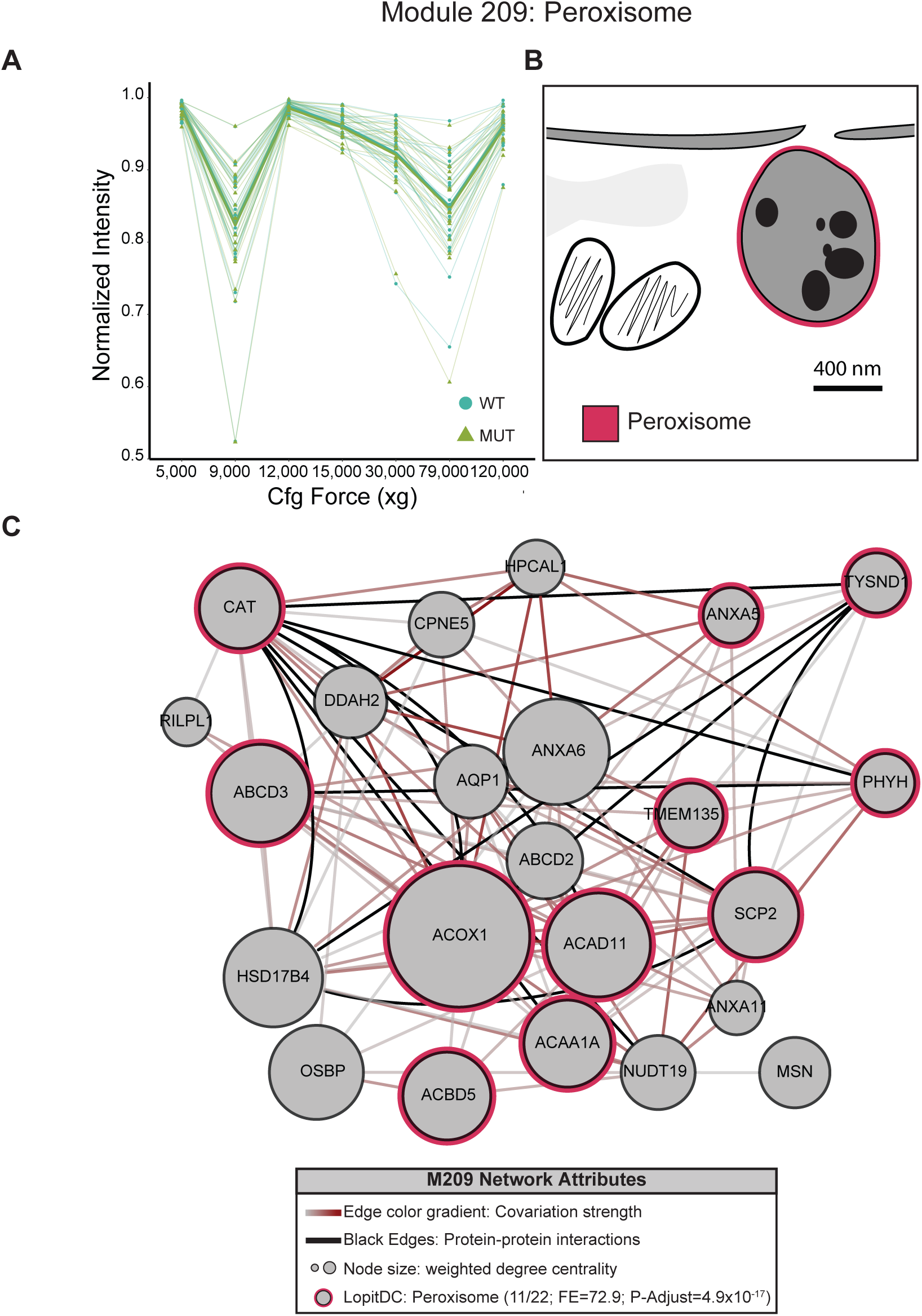
No significant difference in a predicted peroxisome module. (A) normalized protein abundance for all proteins in M209, predicted to represent the peroxisome based on its enrichment of peroxisomal proteins (Geladaki et al., 2019). (B) Schematic of the peroxisome and (C) network of the M209 module which contains 11 of the 22 peroxisomal proteins quantified in this study.

